# Absence of Systematic Effects of Internalizing Psychopathology on Learning Under Uncertainty

**DOI:** 10.1101/2025.05.12.653409

**Authors:** Muhammad H. Satti, Katharina Wille, Matthew R. Nassar, Radoslaw M. Cichy, Nicolas W. Schuck, Peter Dayan, Rasmus Bruckner

## Abstract

Difficulties in adapting learning to meet the challenges of uncertain and changing environments are widely thought to play a central role in internalizing psychopathology, including anxiety and depression. This view stems from findings linking trait anxiety and transdiagnostic internalizing symptoms to learning impairments in laboratory tasks often used as proxies for real-world behavioral flexibility. These tasks typically require learners to adjust learning rates dynamically in response to uncertainty, for instance, increasing learning from prediction errors in volatile environments. However, prior studies have produced inconsistent and sometimes contradictory findings regarding the nature and extent of learning impairments in populations with internalizing disorders. To address this, we conducted eight experiments (*N* = 820) using predictive inference and reversal learning tasks, and applied a bi-factor analysis to capture internalizing symptom variance shared across and differentiated between anxiety and depression. While we observed robust evidence for adaptive learning-rate modulation across participants, we found no convincing evidence of a systematic relationship between internalizing symptoms and either learning rates or task performance. These findings challenge prominent claims that learning difficulties are a hallmark feature of internalizing psychopathology and suggest that the relationship between these traits and adaptive behavior under uncertainty may be more subtle than previously thought.

## Introduction

Learning in complex and changing environments should be calibrated to inherent uncertainties, a process that is especially crucial in aversive contexts. For example, a hiker visiting a predator-rich forest must learn common predator locations to decide which areas to approach with caution. In an area in which bears have previously been sighted, some degree of variability in the exact location and the frequency with which bears appear is expected (aleatoric uncertainty or risk). Therefore, even if the hiker does not encounter a bear on some trips, they should maintain their caution in that area (Fig. 1a). Conversely, if the hiker encounters a bear in a distant and previously safe region, it might indicate a substantial environmental change (epistemic uncertainty), prompting the hiker to update their belief about predator probability more substantially (Fig. 1b). The ability to calibrate learning to the degree of outcome variability and systematic changes is crucial for survival in aversive environments. Notably, this ability has been suggested to be impaired in individuals with high trait anxiety and internalizing psychopathology.

**Figure 1.**
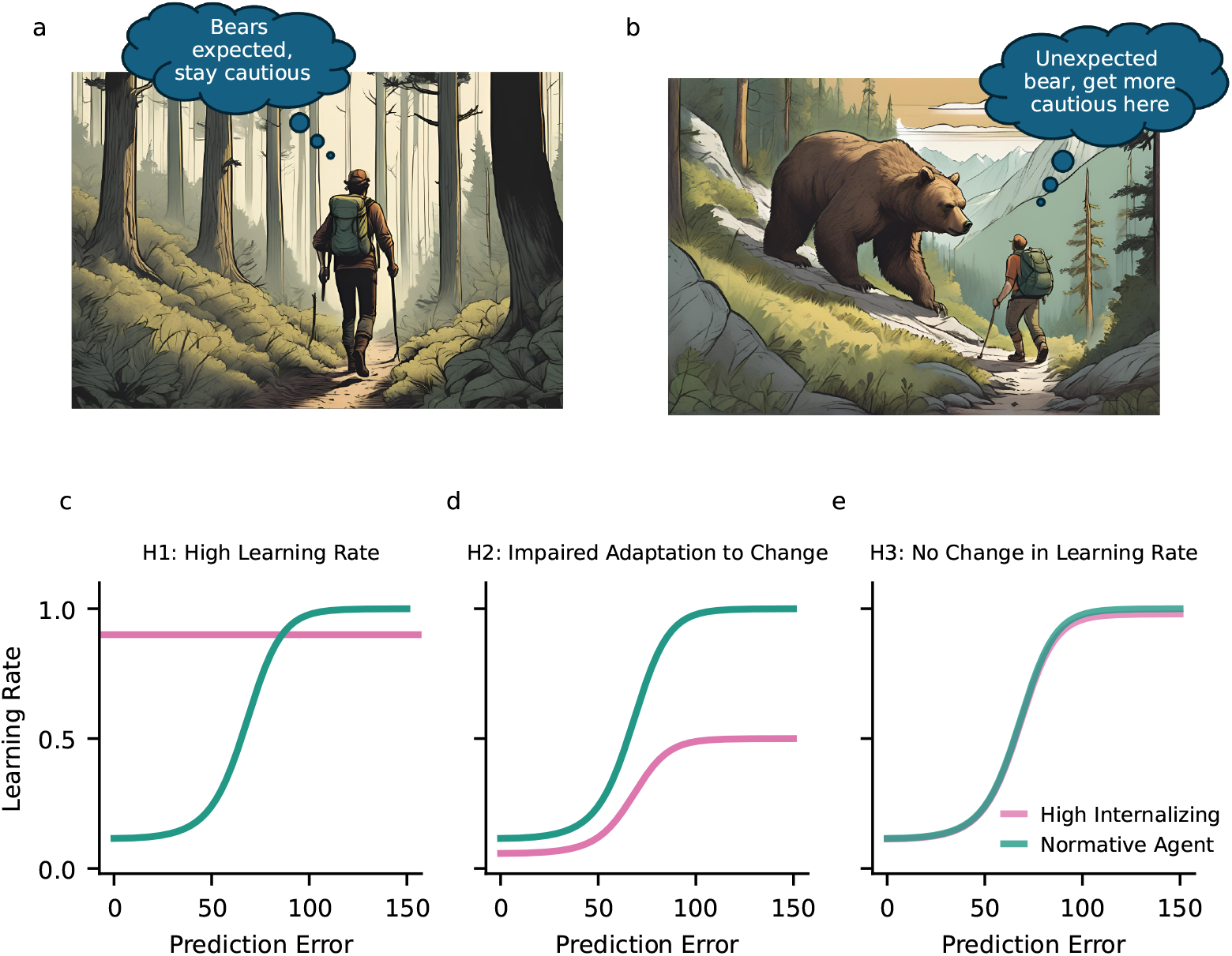
Hypothetical relationship between internalizing psychopathology and learning. To ensure survival, a hiker has to take into account the probability of encountering predators in the forest. Learning the probability that, for example, a bear will suddenly appear requires the ability to regulate the learning rate dynamically, determining the extent of change occasioned by a prediction error (e.g., when the bear appears unexpectedly). **a**| On the one hand, this requires lower learning rates in response to risk due to random outcome variability (aleatoric uncertainty). For example, the hiker should remain cautious in a potentially dangerous area despite not encountering bears every time. **b**| On the other hand, beliefs should be substantially changed via a high learning rate in response to environmental changes that demand new learning (epistemic uncertainty). That is, upon encountering a bear in a previously safe area, the hiker should become more cautious around there. **c-e**| Hypothetical relationships between internalizing psychopathology and the modulation of learning. Green lines illustrate the learning rate simulated from one form of normative Bayesian models called the reduced Bayesian model (details in Methods Reduced Bayesian model). In the range of smaller prediction errors, mainly due to risk, the model uses a lower learning rate. For larger prediction errors related to environmental changes, the model adopts a higher learning rate. Pink lines illustrate hypothetical learning rates in individuals with internalizing psychopathology. **c**| H1: Higher overall learning rate, leading to overlearning from prediction errors. **d**| H2: Impaired learning about environmental changes, selectively concerning larger prediction errors. **e**| H3: No systematic relationship between learning rates and internalizing psychopathology. Graphics in (a) and (b) generated through Canva Magic Media.

From a Bayesian perspective, in environments that can experience sudden and large changes, uncertainty should regulate the learning rate determining how strongly prediction errors (difference between actual outcomes and expected outcomes) are considered for learning (Adams & MacKay, 2007; Bruckner, Heekeren, & Nassar, 2025; Dayan et al., 2000; Mathys et al., 2011; Nassar et al., 2010; O’Reilly, 2013; Piray & Daw, 2021, 2024; Wilson et al., 2010; Yu & Dayan, 2005). When outcomes are corrupted by random environmental variability or risk, one should use a low learning rate to average out prediction errors occurring by chance. In contrast, surprisingly large prediction errors signaling environmental changes should be weighted more heavily by setting a high learning rate. Previous studies that used computational modeling to elucidate modulation in learning duly identified signatures of optimal adaptation to such uncertain and changing environments in healthy younger adults (Behrens et al., 2007; Franklin & Frank, 2015; McGuire et al., 2014; Meyniel & Dehaene, 2017; Nassar, Bruckner, & Frank, 2019; Nassar, McGuire, et al., 2019; Nassar et al., 2016; Payzan-LeNestour & Bossaerts, 2011; Payzan-LeNestour et al., 2013).

Impaired learning-rate adjustments in uncertain and changing environments have been linked to both clinical and trait anxiety (Aylward et al., 2019; Bishop & Gagne, 2018; Browning et al., 2015; Hein et al., 2021; Lamba et al., 2020; Pike & Robinson, 2022; Pulcu & Browning, 2019; Zika et al., 2023). Moreover, recent work provides evidence of a broader relationship between internalizing psychopathology and impaired learning abilities (Fang et al., 2024; Gagne et al., 2020; Wise & Dolan, 2020). These transdiagnostic approaches focus on internalizing symptoms shared across disorders, such as anxiety and depression, which are often correlated (Axelson & Birmaher, 2001; Kalin, 2020; Li et al., 2023).

However, the literature on the exact nature of learning impairments related to trait anxiety and internalizing psychopathology, and whether they are robust in sub-clinical samples, remains inconsistent across studies. Some studies have found that anxious individuals tend to use higher learning rates, particularly in response to negative prediction errors, which can lead to an overestimation of the probability of aversive events (Aylward et al., 2019; Pike & Robinson, 2022; Piray & Daw, 2021; Wise & Dolan, 2020). Other research suggests that individuals with high trait anxiety (Browning et al., 2015) and internalizing psychopathology (Gagne et al., 2020) exhibit less flexible adjustment of their learning rates in volatile environments, indicating a reduced ability to adapt to changing environmental contingencies. Finally, another group of studies has failed to identify any significant anxiety- or internalizing-related impairments in learning (Hitchcock et al., 2024; Jassim et al., 2025; Schindler et al., 2022; Suddell et al., 2024; Ting et al., 2022; Torrents-Rodas et al., 2013).

To address these inconsistencies across previous studies and systematically investigate the relationship between internalizing psychopathology and learning-rate adjustment, we performed a series of eight experiments with *N* = 820 sub-clinical participants. We tested the following three hypotheses derived from prior work. First, internalizing psychopathology might be associated with higher overall learning rates (Fig. 1c). That is, according to this hypothesis, internalizing is related to overlearning from prediction errors. Second, internalizing psychopathology could be selectively associated with impaired adjustments to environmental changes (Fig. 1d). Accordingly, we would expect differences in learning from larger prediction errors, where internalizing would be associated with lower learning rates. The third hypothesis is that internalizing psychopathology and learning rates are not systematically related (Fig. 1e).

To test our hypotheses, we developed an online, game-based, and aversive version of an established predictive inference task (McGuire et al., 2014; Nassar, Bruckner, & Frank, 2019; Nassar et al., 2010) called the predator task. This task allowed us to link adaptive learning abilities and internalizing psychopathology. We systematically investigated different task conditions across six online experiments (*N* = 751) and two in-person laboratory experiments (*N* = 69) with skin conductance response recordings. Additionally, to compare our results across task domains and with previous studies, three of the online experiments (*N* = 179, 94, and 161) featured a probabilistic reversal learning task that partially or fully replicated the task performed in Behrens et al. (2007) and Gagne et al. (2020). To assess internalizing psychopathology, we administered a comprehensive battery of questionnaires and conducted a bi-factor analysis to extract a general internalizing factor representing the shared variance between anxiety and depression (L. A. Clark & Watson, 1991; Gagne et al., 2020; Simms et al., 2008; Watson et al., 1995). Across the eight studies, our findings corroborate the third hypothesis that internalizing psychopathology is not systematically related to learning-rate impairments. These results challenge the idea that internalizing and trait anxiety impair adaptive learning under uncertainty.

## Results

We recruited *N* = 930 participants via the online recruiting platform Prolific (prolific.com) and from the general population of Berlin to participate in online and in-person experiments. After excluding participants based on headphone screening, attention checks, and due to technical difficulties, the final sample consisted of *N* = 820 participants (age range 18 to 69, mean age 32.4, 417 female, 12 non-binary; for details, see Methods Participants).

Participants completed a comprehensive battery of questionnaires before starting the tasks, including the State-Trait Anxiety Inventory - Form Y1, Y2 (STAI-State and STAI-Trait; Spielberger et al., 1983), State-Trait Inventory for Cognitive and Somatic Anxiety (STICSA-Trait; Ree et al., 2008), Intolerance of Uncertainty Scale (IUS-27; Freeston et al., 1994), Beck’s Depression Inventory (BDI; Beck et al., 1961), Penn State Worry Questionnaire (PSWQ; Meyer et al., 1990), Mood and Anxiety Symptom Questionnaire (MASQ; Watson and Clark, 2012) and Internet Gaming Disorder Scale-Short Form (IGDS9-SF; Pontes and Griffiths, 2015).

We conducted a factor analysis to extract a transdiagnostic factor capturing commonalities of anxiety and depressive symptoms. As expected, responses to anxiety and depression items were significantly correlated (Fig. 2a). Following Gagne et al. (2020), we therefore adopted a “bi-factor” analysis approach with a general transdiagnostic factor G explaining shared variance across anxiety and depression, and factors related to anxiety (F1) and depression (F2) (D. A. Clark et al., 1994). We applied this analysis to data from 748 participants (see Methods Factor Analysis and Supplementary Materials (SM) Factor Analysis for participant exclusion criteria and a justification for the number of factors). The results showed that all questionnaire items had moderate-to-large loadings onto the general factor. Additionally, items related to anxiety loaded on F1, while items related to depression loaded on F2 (Fig. 2b).

**Figure 2.**
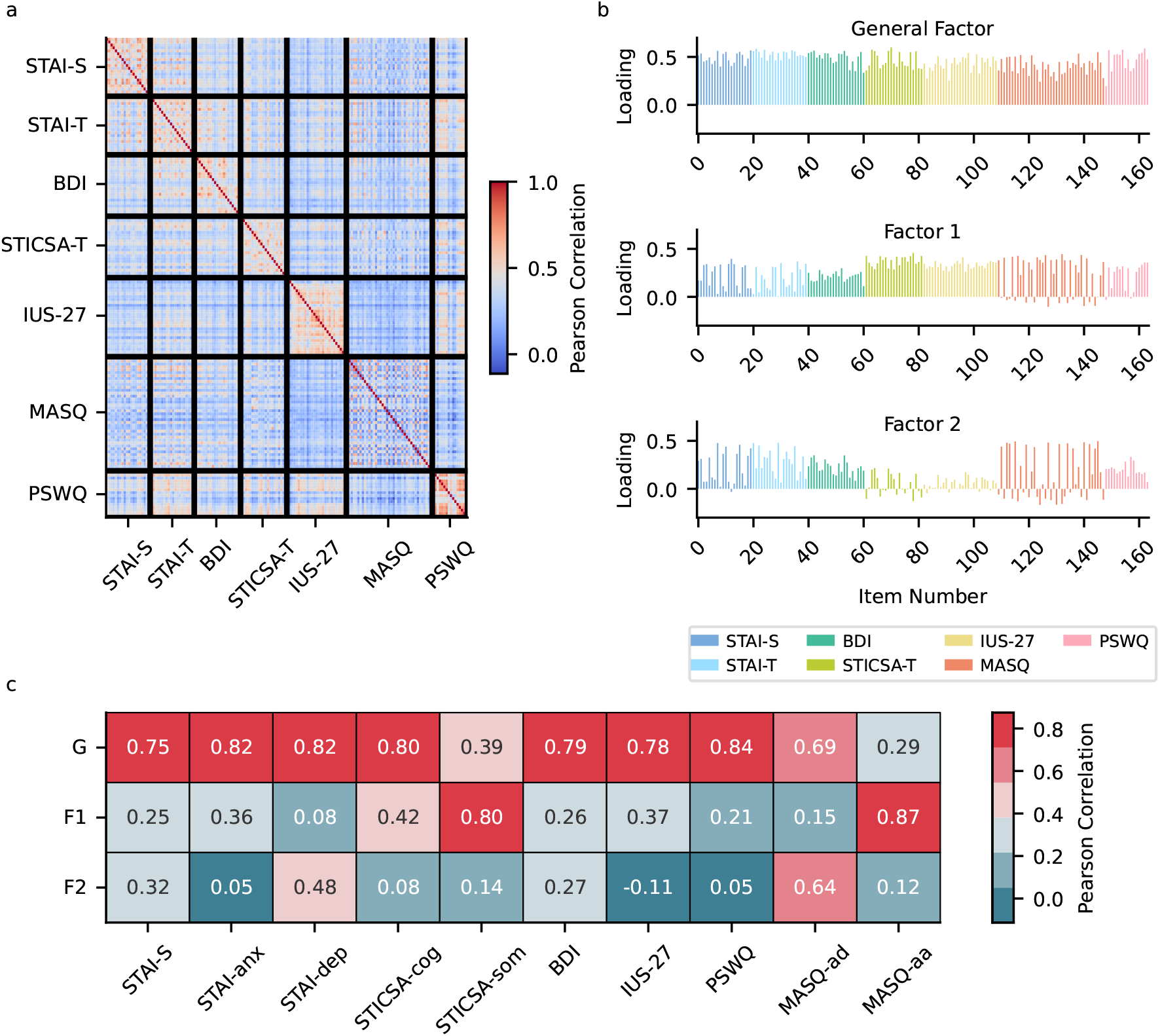
Correlation and factor analysis of questionnaire measures. Questionnaire measures were correlated, so we applied a bi-factor analysis to extract three distinct factors: a general internalizing factor G, an anxiety-related factor F1, and a depression-related factor F2. **a**| Pearson correlation matrix showing the correlation between items of each questionnaire. **b**| Factor loadings of each questionnaire item on the general factor G and the two sub-factors F1 (anxiety related) and F2 (depression related). **c**| The correlation of factor scores with questionnaire sum scores validates the hierarchical 3-factor structure. Based on Cohen’s effect size conventions, the general factor G shows moderate-to-large correlations with all questionnaires and captures variance common to both anxiety and depression, supporting the overall construct validity. F1 has moderate-to-large correlations with the anxiety-related scales MASQ-aa, STICSA-som and STICSA-cog. F2 shows moderate-to-large correlations with depression-related measures like MASQ-ad and STAI-dep. Cohen’s effect size conventions: *r* = 0.10 - small, *r* = 0.30 - moderate, *r* = 0.50 - large. STAI-S: Spielberger State-Trait Anxiety Inventory Y1; STAI-anx: Spielberger State-Trait Anxiety Inventory Y2 - anxiety subscale; STAI-dep: Spielberger State-Trait Anxiety Inventory Y2 - depression subscale; STICSA-cog: State-Trait Inventory for Cognitive and Somatic Anxiety - trait cognitive subscale; STICSA-som: State-Trait Inventory for Cognitive and Somatic Anxiety - trait somatic subscale; BDI: Beck’s Depression Inventory; IUS-27: Intolerance of Uncertainty Scale; PSWQ: Penn State Worry Questionnaire; MASQ-ad: Mood and Anxiety Symptom Questionnaire - anhedonia subscale; MASQ-aa: Mood and Anxiety Symptom Questionnaire - anxious-arousal subscale

We further validated these findings by correlating each participant’s factor scores with summary scores for each questionnaire. For this analysis, the STAI-Trait and MASQ were separated into anxiety- and depression-specific subscales, while the STICSA-Trait was divided into somatic and cognitive anxiety subscales (Fig. 2c). Using Cohen’s conventions for effect sizes (*r* = 0.10 small, *r* = 0.30 moderate, *r* = 0.50 large; Cohen, 1988), we found moderate-to-large correlations of the general-factor scores G with the majority of questionnaires. F1 showed moderate-to-large correlations with anxiety-related measures such as MASQ-anxious arousal (*r* = 0.87, *p* < 0.001), STICSA-somatic (*r* = 0.8, *p* < 0.001), and STICSA-cognitive (*r* = 0.42, *p* < 0.001). F2 showed correlations with the MASQ-anhedonia subscale (*r* = 0.64, *p* < 0.001) and STAI-depression sub-scale (*r* = 0.48, *p* < 0.001). Lastly, we validated our factor structure by comparing our factor scores with those derived using loadings from Gagne et al. (2020), yielding consistent results (see SM Comparing factor structure with Gagne et al. (2020) for details).

### Continuous Predictive Inference Task

#### Task design and model predictions

In the main study, *N* = 563 participants (269 females, 9 non-binary, mean age = 32.3 years, range 18 to 45 years) completed an online, game-based variant of an established predictive inference task called the predator task (Fig. 3a; McGuire et al., 2014; Nassar, Bruckner, & Frank, 2019; Nassar et al., 2010). In this task, participants controlled a flame to defend an avatar against an attacking predator positioned at the center of the screen. The predator attack locations varied on a circle in two critically different ways. First, the attack locations were clustered around a mean position, subject to a circular Gaussian distribution whose variance represented risk.

**Figure 3.**
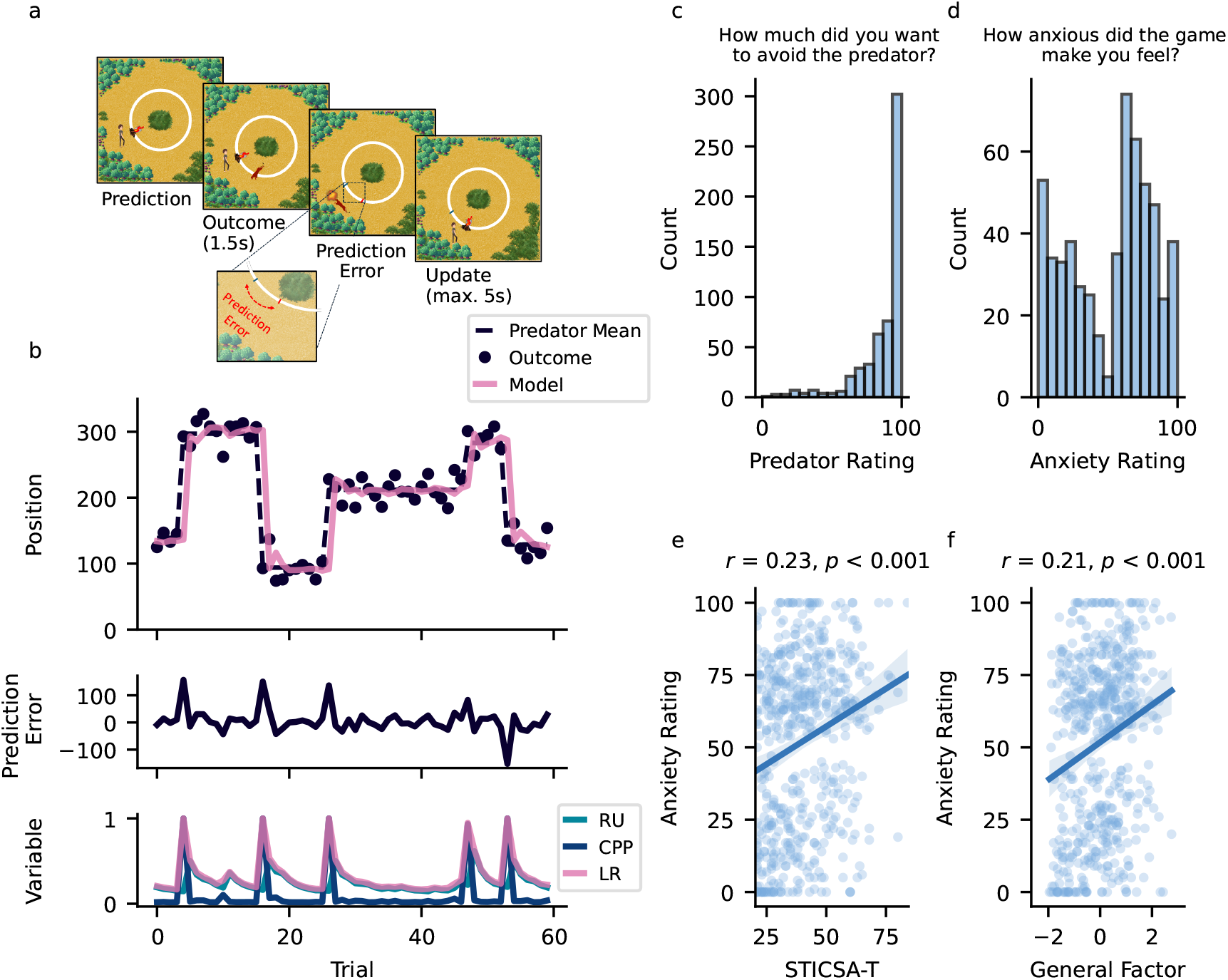
Predator task and reduced Bayesian model. **a**| The goal of participants was to scare away as many attacking predators as possible. To do so, participants placed a flame in the predicted attack location of the predator. When the flame was placed accurately, the predator was scared away. However, when predictions were inaccurate, the predator caught the participants. In each trial, the participants first placed the flame in the expected predator position (Prediction). The predator then started the attack (Outcome), revealing a prediction error (difference between expected and actual location). Finally, participants updated the location of their flame (Update). **b**| The predator’s average attack location (dashed line) is mostly stable but shifts occasionally at change points. Actual attack locations (outcomes, shown by black dots) are corrupted by random variability. We used a reduced Bayesian model with near-optimal learning performance. The model (pink line) calculates prediction errors (second panel) to update its predictions. The learning rate (LR) that controls the influence of prediction errors on belief updating is influenced by change-point probability (CPP) and relative uncertainty (RU). **c**| Histogram showing participants’ responses to the question “How much did you want to avoid the predator?” (task motivation). **d**| Histogram showing participants’ responses to the question “How anxious did the game make you feel?” (task-induced anxiety). **e**| Correlation between trait-anxiety scores (STICSA-T) and task-induced anxiety. **f**| Correlation between the general-factor score and task-induced anxiety. STICSA-T: State-Trait Inventory for Cognitive and Somatic Anxiety - trait subscale

Second, the mean occasionally shifted, without warning, inducing the sort of change points that should inspire more substantial adaptation. Since any attack would be arrested if the flame was placed between the avatar and the predator, participants had to learn the mean of the predator’s attack distribution to protect themselves most effectively. They earned 10 points for each trial in which they successfully defended themselves, and heard an aversive scream when they were caught by the predator.

To test the hypotheses about the relationship between internalizing psychopathology and difficulties in adapting to environmental variability (also referred to as stochasticity or risk) and volatility (the rate of environmental change), we conducted eight experiments using different variations of the predator task. Across these experiments, we systematically manipulated outcome type, variability, volatility, pre-task training, and whether the study was conducted online or in the laboratory. In experiments 1-4, the task included blocks with low and high levels of variability and changes in the mean location of the predator, to assess whether anxiety and internalizing affect task performance under different environmental conditions. Experiments 5 and 6 served as control conditions, with experiment 5 featuring a single variability and volatility level with minimal training and experiment 6 including a full training phase. Experiments 7 and 8 were conducted in the laboratory to validate the aversiveness of screams compared to electric shocks in the predator task. For brevity, we present the combined results from experiments 1 to 4 for the predator task in the Results section, along with results from the laboratory study in experiment 7. We refer to SM for the results from experiments 5,6, and 8 (SM Predator Task). Participants demonstrated a clear understanding of the task. We assessed this through a post-training quiz consisting of three questions completed before starting the main task (details in SM Training Quiz). The majority of participants (70.7%) answered all three quiz questions correctly, 24.7% answered two questions correctly, and 4.3% answered only one question correctly. Only one participant failed to answer any question correctly (Fig. S3). To assess subjective anxiety and whether participants were sufficiently motivated to perform the task, we asked how much they wanted to avoid the predator and how anxious the task made them feel at the end of the task, using a scale from 0 to 100. Participants demonstrated high motivation to avoid the predators (median = 95.0, inter-quartile range (IQR) 81.0 to 100.0; Fig. 3c) and reported high overall subjective anxiety (median = 62.0, IQR 23.0 to 75.0; Fig. 3d). Anxiety ratings were significantly correlated with both trait anxiety (STICSA-T, *r* = 0.23, *p* < 0.001; Fig. 3e) and general-factor scores (*r* = 0.21, *p* < 0.001; Fig. 3f), after controlling for age and gender, indicating that the task sufficiently induced anxiety for studying aversive learning behavior. Moreover, single-trial learning rates in the task exhibited qualitative signatures of adaptive learning (Fig. 4a). Participants used lower learning rates in the range of smaller prediction errors that were primarily driven by random outcome variability, where a low learning rate helped them to average out randomness. In contrast, participants used higher learning rates for larger prediction errors, which enabled them to adjust predictions to change points. This behavioral pattern is consistent with findings from previous studies using similar tasks (Bruckner, Nassar, et al., 2025; Fromm et al., 2025; Jepma et al., 2018; McGuire et al., 2014; Nassar, Bruckner, & Frank, 2019; Nassar et al., 2012, 2016; Piray & Daw, 2024; Vaghi et al., 2017). Therefore, altogether, these results indicate that our task is well-suited to examine threat learning and internalizing psychopathology.

**Figure 4.**
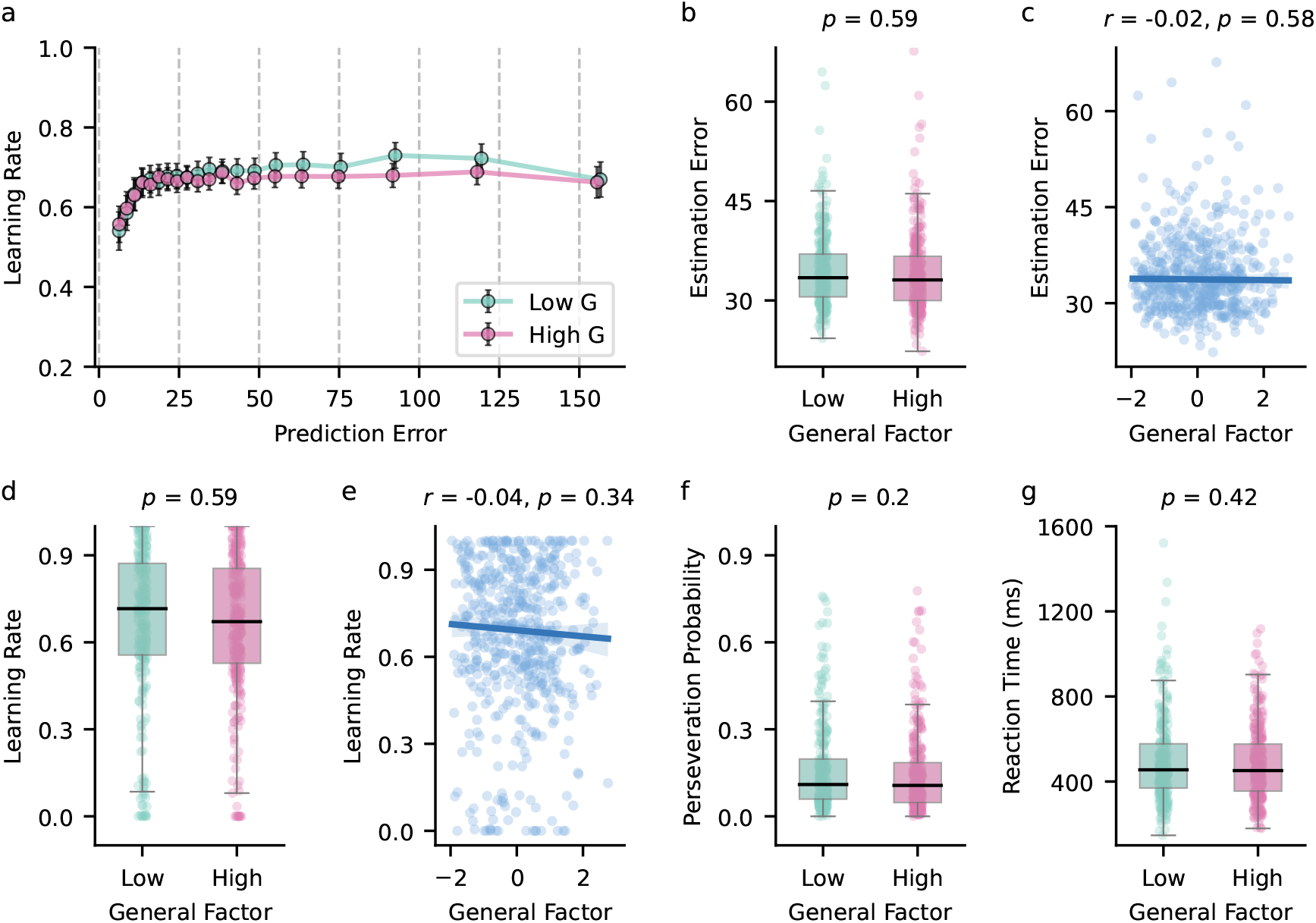
Behavioral results from the predator task. The model-agnostic analysis did not reveal a significant association of internalizing psychopathology (general factor) with behavior in the task. Participants are split into low- and high-internalizing groups for illustration purposes. The low-internalizing group includes participants with a general-factor score below the mean, while the high-internalizing group includes those with a score above the mean. **a**| Learning rates plotted as a function of prediction errors. We divided prediction errors into 20 quantiles showing the mean *±* 95% confidence interval. The plot suggests a similar increase in the learning rate as a function of the prediction error for both groups. **b**| Estimation errors (absolute difference between predator mean and participant prediction), used as a measure of performance in the task, were not found to differ significantly between the two groups. **c**| Analyzing estimation errors across the full spectrum of general-factor scores did not reveal any significant associations. **d**| Average single-trial learning rates were not found to differ significantly between the two groups. **e**| We did not find any significant association between average singletrial learning rates and general-factor scores. **f**| Participants’ perseveration probability (repeating the previous prediction) did not differ significantly between the two groups. **g**| Average reaction times of participants did not differ significantly between the groups.

#### Task learning and performance are not systematically related to internalizing psychopathology

Across several performance measures and ways to quantify learning, we did not find a systematic relationship between internalizing psychopathology and learning impairments. To assess the strength of evidence for the null hypothesis, we calculated Bayes factors (*BF*_01_) based on the corresponding *t*-statistics. For interpretation, we adopted the Jeffreys (1961) classification scheme, where Bayes factors between 1 and 3 indicate anecdotal evidence, between 3 and 10 suggest moderate evidence, and values greater than 10 provide strong evidence for the null hypothesis (Aczel et al., 2018; Lee & Wagenmakers, 2014). To assess participant performance in the task, we analyzed the average accuracy of their beliefs about the predator’s mean location, expressed as the estimation error. The estimation error represents the absolute difference between the participant’s predictions and the mean predator location, and lower estimation errors reflect higher accuracy. To examine the impact of internalizing psychopathology on performance, we divided participants into two groups based on their general-factor scores (low vs. high internalizing).

Subjects of the low-internalizing group had general-factor scores below the mean, and vice versa for the high-internalizing group. Comparisons of estimation errors between the low- (median = 33.43, IQR 30.55 to 36.99) and high-internalizing groups (median = 33.1, IQR 30.0 to 36.66) did not reveal a significant difference (Welch’s two-tailed *t*-test *t*_551.55_ = 0.53, *p* = 0.59, *BF*_01_ = 9.259; Fig. 4b). Similarly, we did not find a significant association between average estimation errors and general-factor scores across participants (*r*(554.0) = -0.02, *p* = 0.58, *BF*_01_ = 17.857; Fig. 4c).

We next examined empirical single-trial learning rates and did not find a systematic relationship between learning behavior and internalizing psychopathology. The empirical learning rate expresses the influence of the prediction error on the belief update, which, in our task, corresponds to how much a participant updates the flame in response to a prediction error. A Welch’s two-tailed *t*-test, which accounts for unequal variances between groups, did not reveal a significant difference in the learning rate between the low-(median = 0.72, IQR 0.56 to 0.87) and high-internalizing groups (median = 0.67, IQR 0.53 to 0.85, *t*_534.6_ = 0.54, *p* = 0.59, *BF*_01_ = 9.174; Fig. 4d). Similarly, the test did not reveal a significant association between learning rates and general-factor scores across participants (*r*(554.0) = -0.04, *p* = 0.34, *BF*_01_ = 13.158; Fig. 4e).

Finally, we examined the perseveration probability, which indicates the probability that participants repeat their predictions in subsequent trials. We did not find significant differences between the low-(median = 0.11, IQR 0.06 to 0.2) and high-internalizing groups (median = 0.11, IQR 0.05 to 0.18, *t*_540.21_ = 1.29, *p* = 0.2, *BF*_01_ = 4.695; Fig. 4f). Similarly, initiation reaction times indicating when the flame was initially moved were not found to be significantly different between the low-(median = 455.0, IQR 369.75 to 577.25) and high-interalizing group (median = 451.5, IQR 355.25 to 575.75, *t*_516.98_ = 0.81, *p* = 0.42, *BF*_01_ = 7.692; Fig. 4g). Together, these analyses suggest comparable performance and learning-rate dynamics across the internalizing spectrum.

These analyses suggest that learning behavior and internalizing are not systematically related to each other, yet one potential concern is that the low- and high-internalizing groups might be best described by different latent phenotypes (Gagne et al., 2020; Wise & Dolan, 2020). To examine this, and to gain a finer-scale, computationally more transparent view of performance, we extracted parameters from a reduced Bayesian model using participant prediction errors – a method extensively validated in prior research (Bruckner, Nassar, et al., 2025; McGuire et al., 2014; Nassar, Bruckner, & Frank, 2019; Nassar et al., 2010). The reduced Bayesian model adjusts the learning rate on each trial in association with the prediction errors (Fig. 3b). This model is formulated in line with the error-correcting delta rule commonly applied in reinforcement learning studies (Dayan & Daw, 2008; Sutton & Barto, 2018). Specifically, the model forms a belief about the predator’s attack location based on the prediction error, expressing the difference between the actual and predicted outcome. The model then scales the prediction error according to a dynamically adjusted learning rate to update beliefs from trial to trial. The dynamic adjustment of the learning rate depends on two parameters, which quantify relative uncertainty and the change-point probability. Relative uncertainty reflects the model’s uncertainty about the predator’s attack location relative to its total uncertainty (sum of environmental variability and uncertainty about attack location). Relative uncertainty is high when only a few trials have been observed, and it decreases over stable trials when the mean location does not move as the model becomes more certain. The change-point probability specifies how likely it is that the mean location of the predator might move between trials. When this happens, prior beliefs are rendered obsolete, relative uncertainty increases, and re-learning is required. Larger prediction errors, signaling potential change points, increase change-point probability and drive higher learning rates for more substantial updates. In contrast, smaller prediction errors, likely resulting from risk, are given a lower weight, leading to subtle adjustments in the expected location of the predator (see Methods Reduced Bayesian model).

Using these extracted parameters, we applied linear regression to quantify the extent to which participants relied on fixed and adaptive learning rates (see Methods First-Level Regression Model). Adaptive learning rates are driven by change-point probability and relative uncertainty, computed by the reduced Bayesian model. In particular, the adaptive learning rate is characterized by larger updates in periods of uncertainty about the predator’s location and following larger prediction errors that are likely due to environmental changes. In contrast, the fixed learning rate reflects a constant influence of the prediction error on the belief update and corresponds to a heuristic learning strategy (Fig. 5a). We assessed the split-half reliabilities of these parameters and found good reliabilities for all parameters, with the fixed learning rate having a Spearman *ρ* = 0.81 and the adaptive learning rate having *ρ* = 0.6 (see SM Split-Half Reliability). These split-half reliabilities indicate that the task and regression model provide a reasonable methodology for inferring the underlying differences in learning and adjustment thereof.

**Figure 5.**
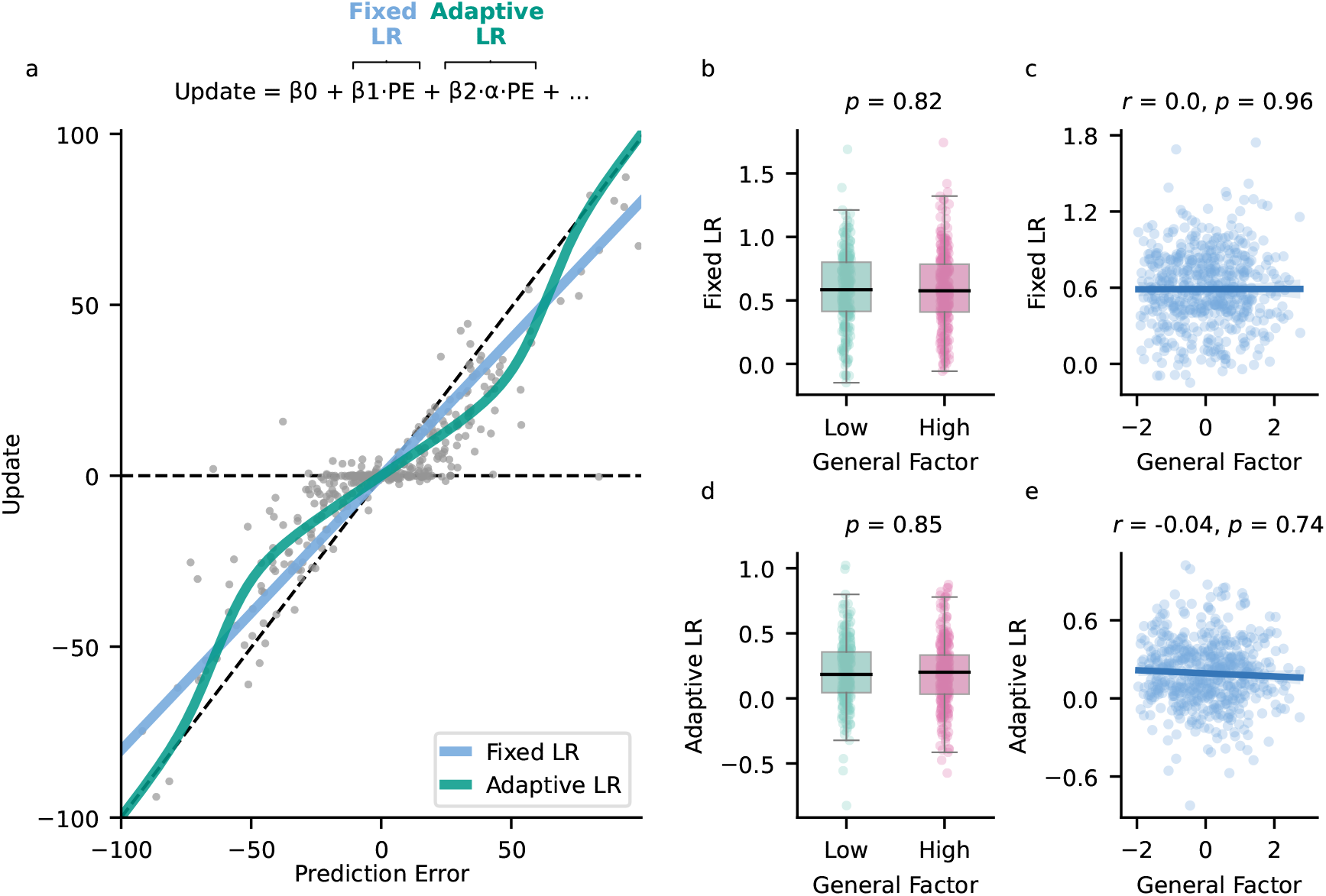
Regression using the reduced Bayesian model reveals no significant effect of internalizing on learning rates in the predator task. **a**| Illustration of the fixed and adaptive learning rate (LR) using example data from one participant. Each gray dot represents a single trial. The fixed learning rate represents a heuristic learning strategy through which the prediction error has a constant effect on the belief update, i.e., there is a linear relationship between update (difference between current and previous flame location) and prediction error. The adaptive learning rate reflects an approximately normative learning strategy, i.e., a non-linear relationship where larger prediction errors increase the learning rate to adaptively respond to change points. Human participants typically show a mixture of fixed and adaptive learning rates. **b**| The low- and high-internalizing groups used fixed learning rates to update their beliefs. However, we did not find significant group differences. **c**| A regression analysis did not yield a significant association between internalizing and fixed learning rates (controlling for age and gender). **d**| Participants also relied on adaptive learning rates, taking into account the underlying environmental dynamics. However, the test did not reveal significant differences between the groups. **e**| Regression analysis of adaptive learning rates similarly did not indicate a significant relationship with internalizing (controlling for age and gender).

Across participants, we found that learning behavior was characterized by a mixture of fixed (median = 0.58, IQR 0.41 to 0.8, Wilcoxon signed-rank *W* = 188.0, *p <* 0.001) and adaptive learning rates (median = 0.2, IQR 0.04 to 0.35, *W* = 19384.0, *p <* 0.001; Fig. S9a). To correct for multiple comparisons when statistically testing this and the following regression parameters, we applied the false discovery rate (FDR) correction (details in Methods False Discovery Rate Correction). The result that participants rely on both fixed and adaptive learning rates is consistent with previous work using similar tasks (Bruckner, Nassar, et al., 2025; McGuire et al., 2014; Nassar, Bruckner, & Frank, 2019; Nassar et al., 2016) and suggests that the task is well-suited for examining the relation between adaptive learning and internalizing.

Analyses examining the link between learning rates and internalizing psychopathology did not indicate any significant association (Fig. 5). We first categorized participants into two groups based on their general-factor scores (low vs. high internalizing) and compared their reliance on the fixed learning rate. A Welch’s two-tailed *t*-test did not reveal a significant difference in fixed learning rates between the low- (median = 0.58, IQR 0.41 to 0.8) and high-internalizing groups (median = 0.58, IQR 0.41 to 0.78, *t*_550.62_ = -0.22, *p* = 0.82, *BF*_01_ = 10.309; Fig. 5b). Similarly, we did not find a significant relationship between fixed learning rates and general-factor scores across participants (*r*(554.0) = 0.003, *p* = 0.959, *BF*_01_ = 20.833; Fig. 5c).

Moreover, we did not find a significant relationship between adaptive learning rates and internalizing psychopathology in the low-(median = 0.18, IQR 0.04 to 0.36) compared to the high-internalizing group (median = 0.2, IQR 0.03 to 0.33, *t*_551.94_ = 0.19, *p* = 0.85, *BF*_01_ = 10.417; Fig. 5d). Additionally, the test did not indicate a significant relationship between adaptive learning rates and internalizing psychopathology across participants (*r*(554.0) = -0.039, *p* = 0.744, *BF*_01_ = 13.333; Fig. 5d). Finally, we did not find any significant association between the model parameters and the general factor G, the anxiety-related factor F1, or the depression-related factor F2 (SM Fig. S9, Table S2).

Taken together, our analysis did not provide convincing evidence for an association between learning impairments and internalizing psychopathology in both model-agnostic and model-based analyses of the predator task. Furthermore, Bayes factors supported the null hypothesis, providing evidence against a relationship between internalizing and learning.

One potential concern was that the extensive task training or the inclusion of multiple variability and hazard-rate conditions might mask the presence of potential impairments. To address this, we conducted an experiment (*N* = 76) using a predator-task version with minimal task training and a single variability and hazard-rate condition (Methods Experimental Details, Experiment 5). Consistent with our previous findings, we did not find any significant learning impairment associated with internalizing psychopathology (in fact, there was some evidence for increased adaptability in this condition; SM Minimal-Training Version Fig. S12). These results were further supported by a control experiment (*N* = 89) in which we used a version of the predator task with a single variability and hazard-rate condition but retaining the full training protocol (Methods Experimental Details, Experiment 6; SM Control Version Fig. S13).

#### Results replicate in a laboratory study with electric shocks and screams as aversive stimuli

A potential concern that still remained was that impairments might only emerge with more intense aversive stimuli, such as electric shocks, as screams may not be sufficiently aversive to reveal these effects compared to shocks. Although existing literature supports the efficacy of screams as effective aversive stimuli (Beaurenaut et al., 2020; Seow & Hauser, 2022), we conducted an in-person laboratory study to address this issue more comprehensively.

In this study, *N* = 29 participants (22 females, mean age = 25.76 years, age range 20 to 41 years) completed four blocks of the predator task. The task consisted of a single variability and hazard-rate condition, with aversive stimuli alternating between shocks (two blocks) and screams (two blocks), counterbalanced across participants (Methods Experimental Details; Experiment 7).

We did not find a significant difference in performance, quantified by estimation errors, between screams and shocks (paired-sample *t*-test *t*_28.0_ = -0.6, *p* = 0.55, *BF*_01_ = 3.236; Fig. 6a). Similarly, we did not find significant differences in single-trial learning rates between scream and shock blocks (*t*_28.0_ = -0.53, *p* = 0.6, *BF*_01_ = 3.344; Fig. 6b). We then used the reduced Bayesian model combined with our regression approach to estimate fixed and adaptive learning rates. Consistent with our results above, the test did not yield a significant difference in fixed learning rates between blocks featuring shocks and screams (*t*_28.0_ = 0.68, *p* = 0.5, *BF*_01_ = 3.096). Similarly, we did not find a significant relationship between general-factor scores and fixed learning rates in the two conditions (*r*_*screams*_(29.0) = 0.08, *p*_*screams*_ = 0.68, *BF*_01−*screams*_ = 4.673, *r*_*shocks*_(29.0) = -0.01, *p*_*shocks*_ = 0.95, *BF*_01 *shocks*_ = 5.051; Fig. 6c). In line with these results, the test did not yield a significant difference in adaptive learning rates between screams and shocks (*t*_28.0_ = -0.27, *p* = 0.79, *BF*_01_ = 3.65), and regression analyses did not reveal significant associations between adaptive learning rates and general-factor scores in either condition (*r*_*screams*_(29.0) = -0.0, *p*_*screams*_ = 0.99, *BF*_01−*screams*_ = 5.076, *r*_*shocks*_(29.0) = 0.23, *p*_*shocks*_ = 0.26, *BF*_01_− _*shocks*_ = 2.809; Fig. 6d).

**Figure 6.**
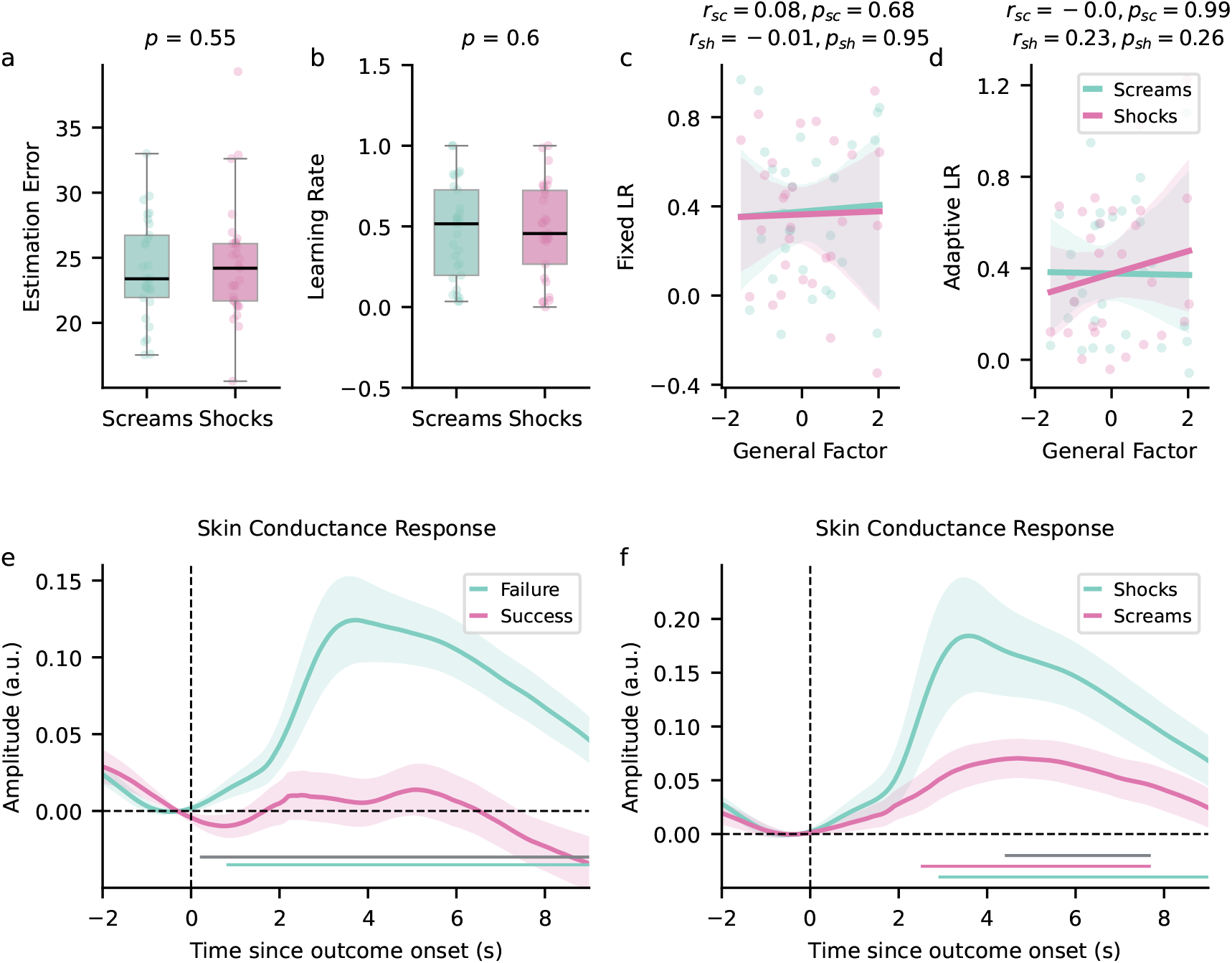
Results from the laboratory study comparing electric shocks and screams as aversive stimuli in the predator task. This study examined the effects of screams and electric shocks as aversive stimuli. **a**| We did not find a significant difference in estimation errors between blocks with screams and shocks. **b**| Analyzing single-trial learning rates similarly did not reveal a significant difference between the two types of aversive stimuli. **c**| Analyzing fixed learning rates based on our regression model did not reveal a significant association between fixed learning rates and internalizing psychopathology for screams or shocks. **d**| Similarly, analyzing adaptive learning rates based on the regression model did not yield significant associations with internalizing psychopathology for both types of stimuli. **e**| Baseline-corrected time course of mean ± standard error of the mean ± (SEM) skin conductance response (SCR) for successful and failed trials. SCR was significantly higher for failed trials compared to successful trials. The horizontal lines at the bottom of the plot show time points where SCRs for failed trials (pink) were significantly different from zero after permutation testing. The gray line shows the significant differences between the two conditions. **f**| For both screams (green) and shocks (pink), the SCR was significantly different from zero. Additionally, SCRs for screams were significantly lower than for shocks (gray line).

The analysis above suggests that behavioral responses elicited by screams are comparable to those elicited by shocks. To further validate the efficacy of screams as aversive stimuli, we analyzed skin conductance responses, correcting for multiple comparisons using cluster-based permutation testing (details in Methods Cluster-Based Permutation Testing). First, we observed a significant increase in the skin conductance response following outcome onset for failed trials (the predator caught the participant) compared to successful trials (pink curve compared to green curve, *p* = 0.001; Fig. 6e). Next, we compared the skin conductance response for the failed trials in shock and scream blocks. This yielded a significant increase after outcome onset for both screams (green curve, *p <* 0.001) and shocks (pink curve, *p <* 0.001, Fig. 6f), confirming that screams effectively function as aversive stimuli. Moreover, in line with the intuition that shocks are more aversive than screams, our comparison between the two types of stimuli revealed that the skin conductance response for screams was significantly lower than that for shocks (horizontal gray line in Fig. 6f, *p* = 0.023). One potential explanation for our comparable behavioral effects despite the differences in the skin conductance response is that the perceived aversiveness that ultimately affects behavior might be similar between the conditions. These findings were further supported by an independent pilot study (Methods Experimental Details, Experiment 8 and SM Fig. S14), which employed a different version of the predator task while maintaining the aversive structure with screams and shocks in separate blocks.

In summary, we did not find significant learning impairments associated with internalizing psychopathology in either model-agnostic or model-based analyses of the predator task. These results were consistent across different task versions and aversive stimuli.

#### Binary Reversal Learning Task

One potential limitation is that our findings may be task-specific and not generalizable to other paradigms commonly used in the literature, most importantly, reversal learning. To address this concern and further test the three hypotheses outlined above, we next relied on three different versions of the established binary reversal learning task (Behrens et al., 2007; Browning et al., 2015; Gagne et al., 2020).

#### Task design and model predictions

To examine reversal learning across the internalizing spectrum, we performed a study including a simplified version of the task in Behrens et al. (2007) and the predator task with *N* = 179 participants (87 females, mean age = 31.68 years, range = 19 to 45 years). In the reversal learning task, participants were required to learn the underlying reward probabilities associated with two fractals to maximize their rewards (Fig. 7a). After making their choice, they received feedback indicating whether the chosen fractal resulted in a reward. Participants used this feedback to update their predictions and gradually learn the reward contingencies of both fractals as the task progressed. The task had two phases: a stable phase and a volatile phase (but with no signal for the transition). During the stable phase, the reward probabilities for both fractals remained consistent, while in the volatile phase, the reward probabilities switched every 20 trials, requiring participants to adapt to the changing contingencies (Fig. 7b). The order of these phases was counterbalanced across participants. This version of the task did not include reward magnitudes for the fractals, focusing solely on assessing participants’ ability to learn the underlying reward probabilities and detect shifts in these probabilities. Results from this version are presented here, as including reward magnitudes can influence behavior in ways that reflect differences in decision strategies rather than differences in learning (Fang et al., 2024). Findings from the two task versions, which incorporated reward magnitudes for each fractal (similar to Browning et al., 2015; Gagne et al., 2020), are detailed in the SM (Task Version With Reward Magnitudes, Task Version with Outcome Magnitudes – Reward and Loss Domain).

**Figure 7.**
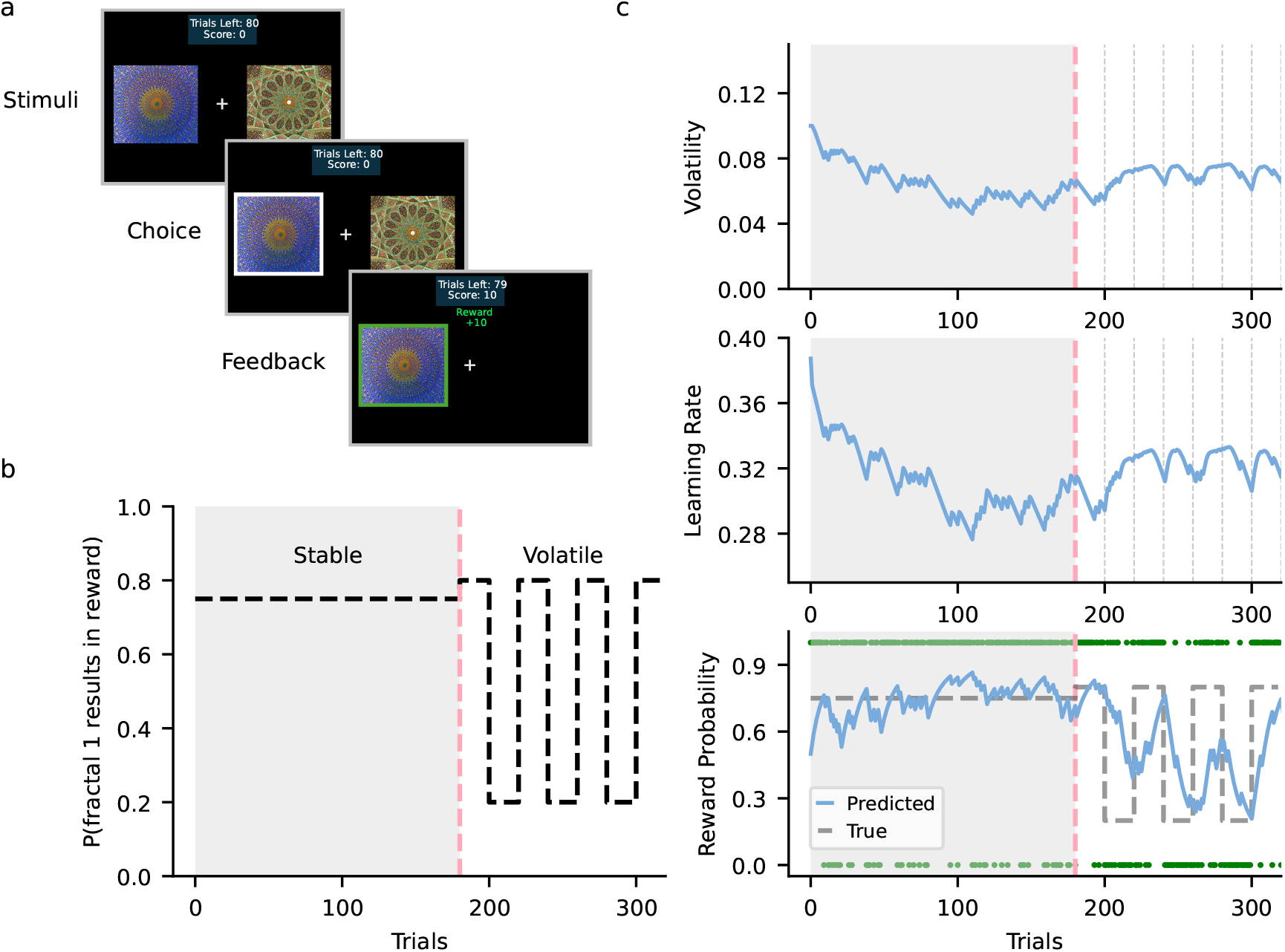
Probabilistic reversal learning task. **a**| The participants’ goal in the task was to maximize rewards. They chose between two fractals on each trial, each associated with a different reward probability. Only one fractal resulted in a reward per trial, and participants were required to learn the reward probability of each fractal. **b**| The task comprised stable and volatile phases. During the stable phase, one fractal had a 75% reward probability and the other 25%. During the volatile phase, reward probabilities switched between 80% and 20% every 20 trials. **c**| From the perspective of a volatile Kalman filter (VKF), learning in the task involves tracking environmental volatility (top), which dynamically adjusts the learning rate (middle). In the stable phase, low volatility results in a lower learning rate, leading to stable reward probability estimates. In the volatile phase, volatility increases after each reversal, raising the learning rate and enabling faster adaptation to shifting reward contingencies (bottom). In the bottom panel, the blue line represents the predicted reward probability of fractal 2, the black dashed line denotes the true reward probability, and the green circles indicate actual outcomes. VKF simulation parameters: *λ* = 0.1, *v*_0_ = 0.1, *ω* = 0.05.

Similar to the predator task, optimal learning behavior in this task can be modeled using a Bayesian inference framework, where the learning rate dynamically adjusts based on the environment’s volatility or rate of change (Behrens et al., 2007; Bruckner, Heekeren, & Nassar, 2025; Piray & Daw, 2020). The volatile Kalman filter (VKF; Piray and Daw, 2020) formalizes this process by continuously tracking environmental volatility and incorporating it into the learning rate in the error-correcting rule of the Kalman filter, allowing flexible adaptation to changing conditions (Fig. 7c). Specifically, in the stable task phase with low volatility, the learning rate remains lower. Conversely, in the volatile phase, frequent reversals in outcome contingencies increase the model’s estimates of volatility, leading to a higher learning rate that facilitates more rapid adaptation to the changing environment.

#### No systematic learning impairments associated with internalizing psychopathology

There was no significant difference between the low- and high-internalizing groups in the performance on this task (Fig. 8a). Similar to the above, we categorized participants into low-internalizing (*N* = 87) and high-internalizing groups (*N* = 92) based on their general-factor scores. Performance was quantified as the proportion of trials where participants chose the highly rewarding or “correct” fractal. Across both the stable and volatile phases of the task, the test did not show significant differences in performance between the high-internalizing group (median stable performance = 0.79, median volatile performance = 0.66) and the low-internalizing group (median stable performance = 0.78, median volatile performance = 0.66; Welch’s *t*-test - stable phase *t*_172.84_ = -1.78, *p* = 0.08, *BF*_01_ = 1.425, volatile phase *t*_169.94_ = -1.72, *p* = 0.09, *BF*_01_ = 1.57; for more details see SM Task Version Without Reward Magnitudes).

**Figure 8.**
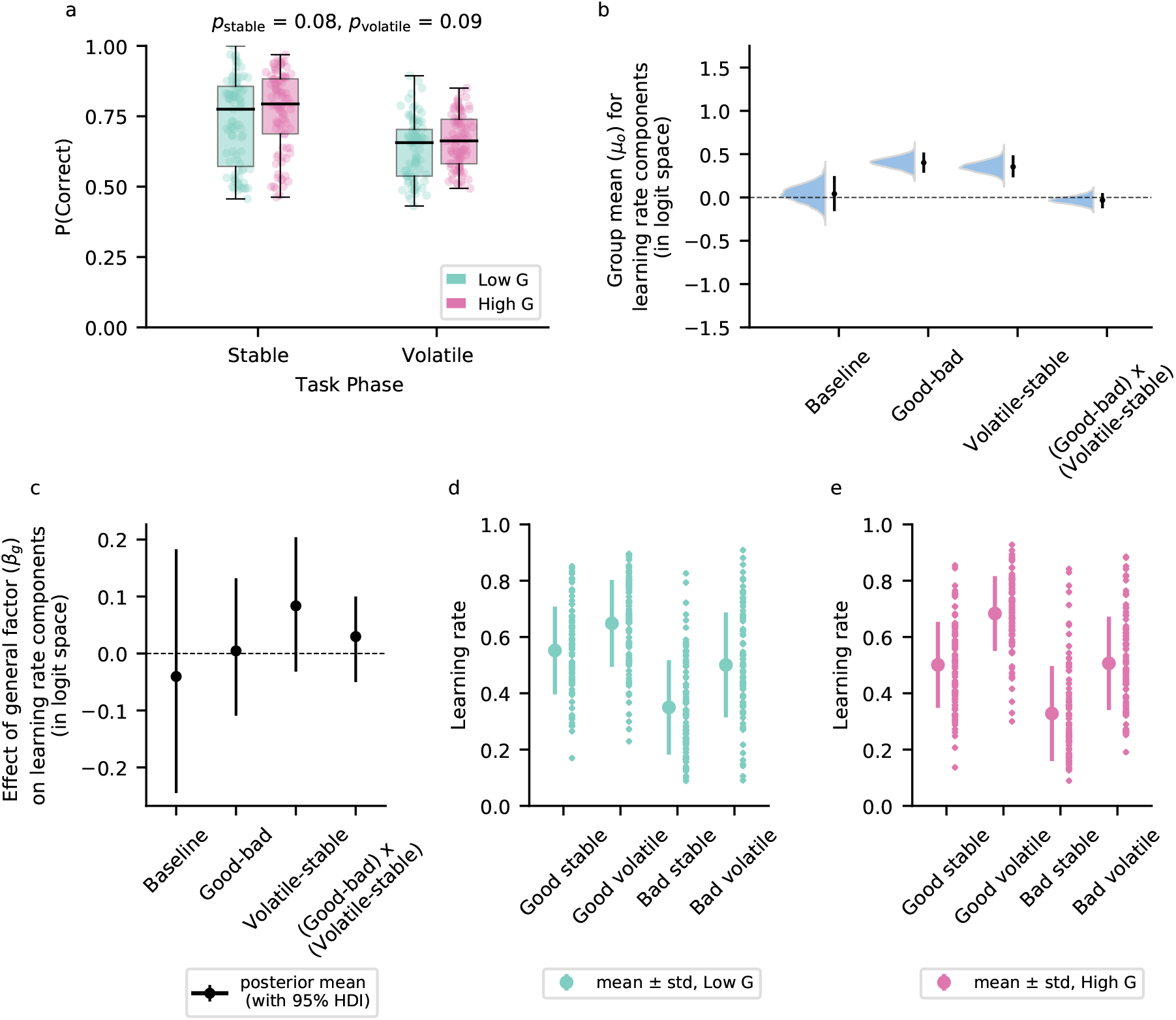
Results from the reversal learning task. **a**| We did not find a significant difference in the probability of choosing the correct fractal between the high- and low-internalizing groups in either the stable or the volatile phase. **b**| Posterior distributions for the mean parameter component of the learning-rate *µ*_0_, with errorbars representing the mean value and 95% highest density intervals (HDI) for each distribution. We found significant effects of outcome valence (good vs. bad) and task phase (volatile vs. stable). **c**| Posterior means and 95% HDIs for the effect of the general factor on learning rates. No significant effect of the general factor was observed, as all HDIs included zero. **d**,**e**| Learning rates depending on outcome valence and task phase plotted separately for the low-internalizing group (**d**) and high-internalizing group (**e**). Small markers show individual participant learning rates, while larger markers represent the average across participants, with the error bars showing the standard deviation.

The results from the previous paragraph suggest comparable performance between low- and high-internalizing groups, contrary to the hypothesis that higher internalizing psychopathology is associated with impaired learning. However, as examined for the predator task, low and high internalizing might be associated with different computational phenotypes. To investigate this possibility, we fitted the VKF (see methods Binary Volatile Kalman Filter) and multiple variants of the canonical Rescorla-Wagner model inspired by Gagne et al. (2020, see methods Model Space) to participants’ learning behavior. The results of the Rescorla-Wagner models are presented here, while qualitatively similar results from the VKF are presented in Fig. S22.

The versions of the Rescorla-Wagner model differed in how their parameters, such as the learning rate and inverse temperature, were further decomposed into components. For instance, in some models, the learning rate was divided into components based on environmental volatility (*α*_*V olatile*_ −_*stable*_), outcome valence (*α*_*Good*_ −_*bad*_), and other factors. Specifically, *α*_*V olatile*_ −_*stable*_ captured the relative change in learning rate across task phases, with positive values indicating increased learning rates in the volatile phase compared to the stable phase. Meanwhile, *α*_*Good*_ −_*bad*_ represented the relative difference in learning rates when learning from rewards versus no rewards. The models were fitted using hierarchical Bayesian estimation, where participants’ factor scores were incorporated in the model parameters (details in Methods Hierarchical model estimation). The population means for each model parameter were composed of an overall mean *µ*_0_ and the three factor-score-specific parameters *B*_*g*_, *B*_1_, and *B*_2_. We assessed and compared models using leave-one-out cross-validation (LOO-CV), estimated through the Pareto-smoothed importance-sampling method (PSIS-LOO). The best-performing model, model 6, incorporated parameters and their interactions related to environmental volatility, outcome valence, and a choice kernel (details in Methods Model Space, SM Table S4). We assessed the reliability of model parameters using a split-half analysis, which revealed poor-to-moderate reliability for the learning-rate components (*α*_*Baseline*_ Spearman *ρ* = 0.5, *α*_*Good*−*bad*_ Spearman *ρ* = 0.23, *α*_*V olatile stable*_ Spearman *ρ* = 0.14, *α*(*Good*− *bad*)x(*V olatile* −*stable*) Spearman *ρ* = -0.01; for details see SM Split-Half Reliability).

We found that, independent of internalizing, learning rates across participants were influenced by environmental volatility and outcome valence. Fig. 8b shows the posterior distributions of the mean *µ*_0_ parameter for the different learning-rate components, where an effect is considered significant if the 95% highest-density interval (HDI) does not include zero. Participants learned more after good outcomes compared to bad outcomes (*α*_*Good* − *bad*_ *µ*_0_ = 0.4, 95% HDI = [0.29, 0.52]). Additionally, learning rates were higher in the volatile phase compared to the stable phase (*α*_*V olatile*−*stable*_ *µ*_0_ = 0.35, 95% HDI = [0.23, 0.49]). We did not find a significant interaction of outcome valence and task phase in relation to the learning rate (*α*(*Good* −*bad*)x(*V olatile*− *stable*) *µ*_0_ = -0.03, 95% HDI = [-0.12, 0.05]). These results are in line with previous research using reversal learning tasks (Behrens et al., 2007; Gagne et al., 2020) and support its suitability to further investigate the role of internalizing psychopathology in learning impairments.

Therefore, we next examined group differences based on our model, but, consistent with the earlier analyses, we did not find a significant effect of internalizing on the learning rate. In particular, we investigated whether internalizing was related to the consideration of outcome valence, volatility, and their interaction. The analysis did not yield a significant impact of internalizing on the consideration of valence (*α*_*Good*−*bad*_ *β*_*g*_ = 0.0, 95% HDI = [-0.11, 0.13]), volatility (*α*_*V olatile*−*stable*_ *β*_*g*_ = 0.08, 95% HDI = [-0.03, 0.2]), and the interaction between the two (*α*(*Good bad*)x(*V olatile* − *stable*) *β*_*g*_ = 0.03, 95% HDI = [-0.05, 0.1], with all 95% HDIs containing zero). Additionally, neither the anxiety-related (F1) nor depression-related (F2) factors were significantly associated with learning-rate components (SM Fig. S19).

While previous analyses did not indicate any significant differences in learning between individuals with low and high internalizing symptoms, we did observe an association between internalizing and choice stochasticity. Internalizing had a significant effect on the baseline inverse temperature parameter, with higher internalizing linked to an increase in the parameter value, leading to less choice stochasticity (*B*_*Baseline*_ *β*_*g*_ = 0.19, 95% HDI = [0.05, 0.34]). Similarly, the depression-related factor (F2) was significantly associated with an increase in the baseline inverse temperature (*B*_*Baseline*_ *β*_2_ = 0.15, 95% HDI = [0.01, 0.28]). These results suggest that higher internalizing symptoms are associated with more deterministic choice behavior. However, this pattern appears to reflect a general decision-making heuristic, rather than differences in learning, and did not lead to statistically significant differences in task performance.

## Discussion

Impaired learning under uncertainty has long been associated with trait anxiety and internalizing psychopathology. However, existing evidence on this relationship is inconsistent between studies. To address this, we conducted eight comprehensive online and in-person experiments, incorporating multiple tasks and physiological measurements. Our findings offer reasonably strong support for the absence of a systematic relationship between internalizing psychopathology, trait anxiety, and impaired learning abilities.

Prior work suggested three potential relationships between internalizing psychopathology and learning behavior. First, it might be associated with higher learning rates, specifically in response to negative outcomes (Aylward et al., 2019; Pike & Robinson, 2022; Wise & Dolan, 2020). Second, it could be linked to a selective impairment in responding to environmental changes (Browning et al., 2015; Gagne et al., 2020). Third, and ultimately corroborated by our results, internalizing and learning impairments might not be systematically related to each other (Hammond et al., 2023; Hitchcock et al., 2024; Suddell et al., 2024; Ting et al., 2022).

There are a number of potential reasons for the discrepancies in prior work that we aimed to address in our study. Inconsistencies may stem from insufficient reliability of tasks and computational assays, compromising the generalizability of findings. This is an important concern for computational psychiatry as a whole (Enkavi et al., 2019; Hedge et al., 2018; Karvelis et al., 2023). For instance, while behavioral measures in reversal learning tasks can exhibit good reliability, the reliability of model parameters varies considerably, with some evidence suggesting that hierarchical models help improve reliability (Schaaf et al., 2024; Waltmann et al., 2022). Similarly, behavioral measures of predictive inference tasks generally show good reliability, but computational measures yielded mixed results in a recent study (Loosen et al., 2024).

In the present study, we observed moderate-to-good split-half reliabilities for all model parameters in the predator task. Moreover, to address the potential influences of practice on reliability (Karvelis et al., 2023), we incorporated an extensive training phase before the task and conducted a separate experiment without training to confirm that training did not bias the main results. Additionally, given the increased reliability of hierarchical models in reversal learning tasks, we used hierarchical Bayesian estimation methods to model behavior in these tasks. Nevertheless, our reliability analysis of the reversal learning task revealed mixed results, with learning-rate parameters exhibiting poor-to-moderate reliability, consistent with prior findings (Suddell et al., 2024). This variability in parameter reliability raises concerns about the robustness and generalizability of previously reported effects in the literature.

Another potential source of the discrepancies in results reported in the literature stems from the heterogeneity of tasks and computational models employed (Nair et al., 2020). Many studies rely on findings derived from bespoke learning tasks, which limits the comparability of results across studies. Additionally, the computational models used to analyze behavioral data vary widely, and researchers often employ different versions of the Rescorla-Wagner model or Bayesian frameworks to estimate learning rates (Pike & Robinson, 2022). Recent work by Eckstein et al. (2022) highlights the problems with comparing model parameters across tasks, showing that parameters – particularly those related to learning rates – are highly context-dependent and exhibit limited generalizability across tasks. This context dependence likely contributes to the inconsistent findings regarding the effects of trait anxiety and internalizing on learning. To address this issue, we investigated learning dynamics across two distinct tasks using multiple model families, including Bayesian models, hierarchical Rescorla-Wagner models, and volatile Kalman filters. These computational models were designed to be flexible, capturing variance in learning behavior driven by various factors such as fixed biases in learning, differential learning from positive versus negative outcomes, and changes in learning rates in response to environmental uncertainty. Importantly, these models were grounded in established methodologies (Bruckner, Nassar, et al., 2025; Gagne et al., 2020; Nassar, Bruckner, & Frank, 2019; Piray & Daw, 2020), ensuring robust analyses while maintaining relevance and comparability to prior work. However, these models still do not encompass the entire range of possible models. Potential future work can explore the impact of different analyses and model specifications on results obtained from the same datasets (e.g., Orben & Przybylski, 2019).

Moreover, some of the previous studies on anxiety and learning relied on small to modest sample sizes ranging from 30 to 147 participants (Aylward et al., 2019; Browning et al., 2015; Gagne et al., 2020; Hammond et al., 2023; Hein et al., 2021; Huang et al., 2017), which limits statistical power and reduces the ability to detect effects reliably (Button et al., 2013; Gelman & Carlin, 2014; Suddell et al., 2024). This is of particular importance given that recent work suggests these effects to be small and inconsistent (Hammond et al., 2023). Additionally, because small samples are only sufficiently powered to detect larger effects, their combination with field-wide biases, such as publication bias and the file-drawer effect, leads to selective reporting of significant findings. This inflates effect sizes, making them difficult to replicate and contributing to inconsistencies in the literature (Button et al., 2013; Ioannidis, 2005). When combined with the flexibility in data collection and analysis, this increases the risk of false positives (Orben & Przybylski, 2019; Simmons et al., 2011; Simonsohn et al., 2019), ultimately adding to the inconsistencies observed in the field.

To ensure sufficient power to detect medium-sized effects, we recruited a large sample (*N* = 820) across multiple experiments, including 563 participants in the main predator task and 179 participants in the reversal learning task. This design provides at least an 80% power to detect medium effects (*r* ≥ 0.25), should they exist. Despite this, we did not find convincing evidence of impaired learning under internalizing – aligning with a recent high-powered study with clinical samples that similarly failed to detect such effects (Suddell et al., 2024). These findings suggest that any effects, should they exist, are likely small (*r*≤ 0.25), a range where our study lacks sufficient power. If effects are small, then, even if they can reach statistical significance in large samples, they may not translate into meaningful behavioral differences (Gelman & Stern, 2006). Addressing this issue in future research could take two complementary approaches. From a clinical perspective, it would be valuable to determine the magnitude of learning-rate differences that manifest as observable behavioral changes. From a computational perspective, simulations could be used to model behavior under varying parameter values, helping to identify the point at which differences in learning parameters produce meaningful shifts in decision-making. Together, these approaches could clarify what the true effect, if it exists, should look like to be considered an impairment.

A potential concern with online participant recruitment is data quality, which depends on participants’ attention and motivation (Jun et al., 2017). One promising approach to improving data quality is task gamification – designing tasks to be more engaging and immersive (Long et al., 2023). Gamified tasks not only enhance participant motivation but also create decisionmaking environments that more closely resemble real-world contexts, making them intuitive and interactive (Allen et al., 2024; Benrimoh et al., 2023). In line with this, the predator task was designed as a game in which participants had to defend themselves against attacking predators, with the environment structured to incorporate principles of predictive inference. This approach proved effective, as participants reported high task motivation. Additionally, to minimize the risk of spurious correlations arising from inattentive responses to psychological questionnaires (Zorowitz et al., 2023), we excluded participants who failed attention checks embedded in the questionnaires. We also excluded participants who failed the headphone checks, thus filtering out potentially unmotivated participants who may have disregarded instructions. These methodological considerations aimed to increase the robustness of our findings. Furthermore, our results were consistent with those obtained in a laboratory setting, both in behavior and task aversiveness, as well as with results that we obtained from a reversal learning task previously used in the literature, adding further credibility to our conclusions.

A recent large-scale study also did not find convincing evidence of learning impairments associated with internalizing in a clinical sample with anxiety and depression (Suddell et al., 2024). However, it reported poor reliability of model parameters in the reversal learning task, leaving open the question of whether inconsistencies in the literature stem from unreliable parameters or a genuine absence of an effect of internalizing on learning. In our study, the predator task yielded more reliable model parameters than the reversal learning task, alleviating concerns that our findings are merely driven by reliability issues. Additionally, to our knowledge, our study is the first to systematically examine the impact of internalizing on learning in a large subclinical sample across two distinct tasks, both online and in-person, and across multiple task variants. The consistency of our findings across tasks, coupled with their alignment with Suddell et al. (2024), provides further support for the hypothesis that internalizing is not systematically linked to impaired learning under uncertainty. Nonetheless, an important limitation of our study is that we did not include clinical subjects. Therefore, our results might not generalize to clinical samples, and future work could address this comprehensively. Additionally, impairments related to internalizing and social anxiety may be more pronounced in social learning contexts (Lamba et al., 2020; Müller-Pinzler et al., 2019), which were not tested in our study and remain an important direction for future work.

In conclusion, despite several influential studies reporting a link between internalizing psychopathology and impaired learning under uncertainty, our findings provide no convincing evidence for the existence of a systematic relationship. By leveraging two well-established tasks – spanning both continuous and discrete binary learning domains – our study offers one of the most comprehensive and large-scale investigations of this question to date. These results help clarify a previously inconsistent literature and suggest that the relationship between internalizing traits and adaptive learning may be weaker or less robust than previously assumed.

## Methods

### Participants

In total, we recruited *N* = 872 participants (Table 1) from Prolific (https://www.prolific.com) for the online studies (experiment 1-6). Eligibility was limited to residents of the UK and USA with a Prolific approval rating of 94 or above and fluency in English. Screening tests ensured a gender-balanced sample. Participants who failed the in-game sound checks or who failed more than 2 attention-check questions out of 5 were excluded from further analysis. In experiments 2, 3, and 4, half of the participants were required to have experienced symptoms of anxiety in the past. This was done to ensure a wider spectrum of participants and symptomatologies. Participants provided informed consent for data collection and storage before starting the studies. For experiments 7 and 8, *N* = 76 participants were recruited from Berlin’s student and general population to participate in the laboratory, where the predator task included physiological measurements and electric shock delivery. Participants were required to be between the ages of 18 to 45. They provided written informed consent to take part in the studies. The experimental protocols were approved by the ethics committee of Freie Universität Berlin (“Adaptives Lernen unter Unsicherheit und unterschiedlichen Bedrohungslagen”, protocol number: 013/2023).

**Table 1.** Basic task and demographic details of all conducted experiments. Predator Task*: Version of predator task with differing variability and hazard-rate conditions; results of this task version are reported in the main analysis.

| Experiment | Tasks | Total Participants | Number After Questionnaire Checks | Number After All Exclusions (# females) | Mean Age ± sd |
| --- | --- | --- | --- | --- | --- |
| 1 | Predator Task* | 175 | 173 | 134 (62) | 33.84 ± 6.97 |
| 2 | Predator Task* + Reversal Learning | 210 | 205 | 189 (90) | 31.53 ± 6.66 |
| 3 | Predator Task* + Reversal Learning (Reward Magnitudes) | 103 | 100 | 95 (47) | 32.18 ± 6.73 |
| 4 | Predator Task* + Reversal Learning (Reward & Loss Version) | 194 | 188 | 169 (85) | 31.97 ± 6.83 |
| 5 | Predator Task (Minimal Instructions) | 93 | 91 | 76 (36) | 35.76 ± 12.05 |
| 6 | Predator Task (Control Study) | 97 | 96 | 89 (43) | 35.5 ± 11.72 |
| 7 | Predator Task (Laboratory Study - with shocks) | 35 | 35 | 29 (22) | 25.76 ± 5.01 |
| 8 | Predator Task (3 predators) (Laboratory Study - with shocks) | 41 | 41 | 40 (32) | 24.95 ± 4.83 |

### Psychological Questionnaires

Participants completed a battery of widely used psychological questionnaires before starting the tasks. These questionnaires were selected to measure symptoms related to anxiety, intolerance of uncertainty, and depression and included the State-Trait Anxiety Inventory - Form Y1, Y2 (STAI-State, STAI-Trait; Spielberger et al., 1983), State-Trait Inventory for Cognitive and Somatic Anxiety (STICSA-Trait; Ree et al., 2008), Intolerance of Uncertainty Scale (IUS-27; Freeston et al., 1994), Beck’s Depression Inventory (BDI; Beck et al., 1961) and Internet Gaming Disorder Scale - Short Form (IGDS9-SF; Pontes and Griffiths, 2015). In experiments 1-5, the Penn State Worry Questionnaire (PSWQ; Meyer et al., 1990) and Mood and Anxiety Symptom Questionnaire (MASQ; Watson and Clark, 2012) were also included to align more closely with the measures used in Gagne et al. (2020). The administration of the questionnaires and data storage were managed using SocSci Survey (https://www.soscisurvey.de).

### Exclusion Criteria

#### Questionnaire Attention Checks

Inattentive responses to questionnaire items can result in spurious correlations between task behavior and symptom measures (Meade & Craig, 2012; Zorowitz et al., 2023). To identify and control for inattentive participants, we embedded five infrequency items (e.g., “I work fourteen months in a year”) into our questionnaires. These questions were designed to have only one or two plausible correct answers. Participants who failed more than two of these infrequency items were excluded from the data analysis.

#### Headphone Screening

To ensure participants used headphones and maintained appropriate sound levels necessary to hear the aversive screams, we administered headphone screening checks at the beginning and midpoint of the predator task (Milne et al., 2021). For each headphone check, participants underwent three trials where they had to identify a beep among three consecutively presented tones. Two of these tones were white noise, while the tone containing the beep was designed to be audible only through headphones, not loudspeakers. Participants needed to correctly identify the beep in at least two out of three trials across two attempts. If they failed to meet this criterion either at the start or during the task, their data were excluded from further analysis. Exclusion rates for each experiment are detailed below.

#### Performance-Based Exclusion

Participants who did not complete all blocks of a task were excluded from further analysis. In the probabilistic reversal learning task, participants who did not make any choice for more than 15% of the trials or who consistently selected images from only one side of the screen (either left or right) across both blocks demonstrated either a poor understanding of the task or low motivation. As a result, their data were excluded from further analysis.

#### Factor Analysis

We conducted an exploratory bi-factor analysis on item-level responses from 164 questionnaire items, belonging to STAI-S, STAI-T, STICSA-T, BDI, IUS-27, MASQ, and PSWQ. The analysis included data from *N* = 758 participants across experiments 1-5 who completed all questionnaires and passed the Questionnaire Attention Checks. To improve the interpretability of factor loadings, we applied reverse scoring to flip the response scale of negatively phrased items, ensuring that higher scores consistently reflect the same meaning across all questions.

Following Gagne et al. (2020), we determined the number of factors using a combination of methods: prior research, visual inspection of the scree plot (Cattell, 1966), and parallel analysis (Horn, 1965). Previous studies (Chorpita et al., 2000; L. A. Clark & Watson, 1991; Gagne et al., 2020; Simms et al., 2008) support a three-factor hierarchical structure for anxiety and depression, comprising a general internalizing factor that captures shared variance, and two sub-factors: an anxiety-related factor and a depression-related factor. Consistent with these findings, the scree plot indicated three eigenvalues above the elbow. Parallel analysis, which compared the eigenvalues from our dataset with those generated from a resampled version of the original dataset, further validated the three-factor structure (see SM Factor Analysis).

Next, we performed factor analysis using polychoric correlations, specifying the pre-determined number of factors. An oblique rotation and a Schmid-Leimann transformation (Schmid & Leiman, 1957) were applied to extract two lower-level factors, F1 and F2, and a higher-level general factor G. G exhibited moderate-to-high loadings across all items, reflecting shared variance. F1 and F2 exhibited loadings for anxiety-related and depression-related questionnaire items, respectively. Factor scores for all participants were then calculated using the Anderson-Rubin method, ensuring orthogonality of the factor structure (Anderson & Rubin, 1956).

For experiments 6 and 7, where MASQ and PSWQ were not included in the questionnaire battery, factor scores were derived using factor loadings identified in the above analysis for common questionnaire items.

The factor analyses were carried out in *R* (R Core Team, 2021) using the Psych library, with the fa.parallel function used for the parallel analysis and the omega function used for the factor analysis.

#### Experimental Tasks

We first provide intuitive explanations of our two tasks (predator task and reversal learning task), and then explain the formal task models in detail.

#### Predator Task

The predator task is an online, game-based variant of an established predictive inference task (Fig. 3a) (McGuire et al., 2014; Nassar, Bruckner, & Frank, 2019; Nassar et al., 2010). The participants’ aim in the task was to protect themselves from attacking predators. The task’s cover story set the scene in a jungle clearing where participants gathered gold while predators lurked among the bushes at the center of the screen. At the beginning of each trial, an audio tone alerted participants to an impending predator attack. The predators started off hidden in the forest and typically ran in a specific direction, though with some variability. Using their mouse, participants initiated and controlled a fire, which could be moved in a circle around the center of the screen and placed appropriately to attempt to fend off the predator. After the warning tone, participants needed to predict the predators’ attack direction based on previous trials, and place the fire at that location. Once placed, the fire’s position was fixed for the trial, and its size varied randomly (Fig. 3a, first panel). The predator then revealed itself by charging in a specific direction. If the fire was correctly placed in its path, the predator was warded off and kept confined to the center, with the participants gaining 10 points. If the fire failed to align with the predator’s path, the predator attacked and caught the participants, triggering an aversive scream (Fig. 3a, second panel). If the participants failed to initiate the fire within the allocated time limit (5s), they additionally lost 10 points. Following this outcome, participants received feedback displayed as markers indicating the predator’s actual attack location (in red) and their own prediction (in blue). These markers remained visible while participants made their next prediction, allowing them to see the previous prediction errors (Fig. 3a, third panel). This design aimed to reduce the influence of working memory differences on learning behavior, encouraging participants to adjust their predictions based solely on prediction errors.

The predator attack locations were sampled from a von Mises (circular Gaussian) distribution, with the mean representing the most probable attack direction and the standard deviation indicating the inherent variability in the attack direction. To maximize their chances of survival, participants had to learn the mean of the distribution and place the fire there, while disregarding the errors caused by the outcome variability. After participants placed the fire, the flame size varied randomly on each trial, sampled from a truncated exponential distribution. A fixed flame size could introduce performance biases – if too large, it would always deter predators, while if too small, it would never provide protection. By varying flame sizes, we eliminated this confound, decorrelating torch size from performance while also encouraging participants to make the best possible predictions on each trial. Occasionally, the predator’s attack direction shifted entirely due to a change in the mean of the Gaussian distribution, representing an environmental change point. In the instance of a change point, participants needed to quickly update their beliefs to relearn the new attack direction. Thus, on each trial, prediction errors could result either from random outcome variability or from a genuine change point. The core challenge of the task was to differentiate between prediction errors caused by systematic changes and those stemming from outcome variability and to adaptively use these errors for belief updating.

Before starting the predator task, participants underwent a comprehensive training session, which included a final quiz to assess their understanding of the task and to provide feedback about their understanding (details in SM Training Quiz). The task was programmed in JavaScript, mainly using the Phaser 3 game development framework (https://phaser.io). Google Firebase services were utilized for both hosting the game and managing data storage (https://firebase.google.com/).

#### Task Variants

- **Different variability and hazard rate**: The main version of the predator task (experiments 1,2,3 and 4) employed a 2-by-2 design with 4 distinct conditions, varying in levels of variability (low standard deviation *σ* = 20, high *σ* = 30) and hazard rate (low *h* = 0.1, high *h* = 0.16), presented across 4 blocks. While variability and hazard rate differed between blocks, they remained consistent within each block. In the **low-variability, low-hazard-rate condition**, outcomes were weakly corrupted by variability, making it relatively easier to identify change points. Here, an ideal agent would learn more from large prediction errors, as they likely signal change points. In contrast, in the **high-variability, low-hazard-rate condition**, outcomes were more heavily influenced by variability, making change points harder to detect. In this scenario, an ideal observer would adopt a lower learning rate to avoid overreacting to variability. In the **low-variability, high-hazard-rate condition**, the mean of the predator attack distribution changed frequently. A higher learning rate is advantageous in this condition to cope with frequent changes. Finally, in the **high-variability, high-hazard-rate condition**, the combination of high variability and frequent changes made it difficult to distinguish random variability from true change points, necessitating a lower learning rate. The order of conditions was counterbalanced across participants to control for order effects. This experiment allowed us to analyze the impact of varying variability and hazard rates on learning, and to investigate whether trait anxiety and internalizing psychopathologies are associated with impaired adaptation to variability and environmental changes (Piray & Daw, 2021, 2024). All participants underwent task training with one condition (low variability and low hazard rate) before starting the task.
- **Control task - minimal instructions**: To evaluate whether task training influenced learning, experiment 5 used a task version that provided participants with only minimal instructions and training before they began. In this version, participants completed 6 blocks of the predator task, with each block consisting of 80 trials. All blocks had the same levels of variability (low variability: *σ* = 20 degrees) and hazard rate (low hazard rate *h* = 0.1), but participants received minimal training beforehand. Comparing the results of this task to the main task allowed us to determine whether the lack of observed effects of internalizing psychopathology on learning could be attributed to the comprehensive training phase in the main task.
- **Control task - one condition**: As a control for the minimal-training task version, in experiment 6 we used a task version with a similar structure but included the full training before starting the task. In this version, participants completed 6 blocks of the predator task, where each block consisted of 65 trials. All blocks had the same levels of variability (low variability: *σ* = 20 degrees) and hazard rate (low hazard rate *h* = 0.1). All participants underwent comprehensive task training before starting the task. This design acted as a control condition for the previous experiment, having minimal task training, and also allowed us to analyze learning across a substantial number of trials for a single condition

#### Binary Reversal Learning Task

In this study, we also used different variants of a binary reversal learning task (Fig. 7a; Behrens et al., 2007; Browning et al., 2015; Gagne et al., 2020) along with the predator task. Here, we describe the basic version used in experiment 2, with modifications for experiments 3 and 4 detailed below. On each trial, participants chose between two fractals presented on either side of the computer screen, separated by a fixation cross in the center (Fig. 7a, top panel). Each fractal was probabilistically associated with the receipt of a reward, and participants aimed to select the fractal most likely to yield a reward on that trial (Fig. 7a, middle panel). After making their selection, participants were provided with feedback in which only the rewarding fractal remained on screen, and participants received 10 points if this was the fractal they selected (Fig. 7a, bottom panel). Participants had 6 seconds to make a choice, and if they did not respond within this time, the trial was marked as ‘failed’, and 10 points were deducted from their total number of points. Feedback then indicated the rewarding fractal for that trial. To control for location-related choice biases, we randomized the fractal location on the screen. The main goal of the task was for the participants to learn the underlying reward probabilities of each fractal and maximize their points by consistently choosing the fractal with the highest probability of reward.

The task was divided into two distinct phases: a stable phase and a volatile phase, each consisting of 2 blocks with 80 trials per block. In the stable phase, the reward probabilities remained constant throughout; one fractal had a 75% chance of delivering a reward, while the other fractal had a 25% chance. In contrast, in the volatile phase, the highest reward probability was increased to 80%, but the fractal associated with the highest probability changed every 20 trials. This phase required participants to continually update their beliefs in response to shifts in reward contingencies. Participants were not informed about these phases or their distinct structures. To control for potential order effects, we counterbalanced the order of the stable and volatile phases across participants.

#### Task Variants

1. **Reversal learning task**: Reversal learning task described above.
2. **Reversal learning task with outcome magnitudes**: In the reversal learning task in experiment 3, each fractal was also associated with reward magnitudes, displayed within a circle above each fractal. These magnitudes were independent of the reward probabilities but determined the number of points earned by the participants if they chose the correct fractal. Reward magnitudes varied randomly from 1 to 99 for each fractal and changed from trial to trial. Participants were instructed to consider both the reward probabilities and reward magnitudes of the fractals while making a choice on each trial. The design of the reversal learning task with reward magnitudes aligns with previously established task versions in the literature (Behrens et al., 2007; Browning et al., 2015; Gagne et al., 2020).
3. **Reversal learning task with outcome magnitudes - reward and loss domain**: In the version of the task in experiment 4, participants completed both a reward-based and a loss-based reversal learning task with outcome magnitudes. In the reward version, reward magnitudes were displayed above the fractals, and participants earned the corresponding rewards for selecting the correct fractal. Conversely, in the loss version, participants started with 10,000 points and lost the corresponding magnitude displayed above the fractal when they chose the incorrect one. While the tasks differed in their framing (reward vs. loss), the underlying structure was the same. Each task further consisted of a stable and a volatile block, with 90 trials in each block. The order of the two tasks was counterbalanced across participants, and each participant was randomly assigned to one of two schedules, which predetermined the outcomes and magnitudes. This design, consistent with prior research (Gagne et al., 2020), enabled us to investigate whether the outcome domain (reward vs. loss) influences learning in individuals with internalizing psychopathology.

#### Experimental Details

**Experiment 1** In this experiment, *N* = 175 participants (84 females, 2 non-binary, mean age = 32.86 years, age range = 18 to 45 years) were recruited from Prolific to complete 4 blocks of the predator task with differing variability and hazard rate conditions, each block consisting of 120 trials. Participants were compensated £8 for their participation in the study, with the opportunity to earn an additional 50 pence bonus for each block in which their score exceeded 550 points. Data from 1 participant were excluded for failing the questionnaire attention checks. Data from 23 participants were excluded from subsequent analysis for failing the headphone sound checks, while data from 17 participants were additionally excluded for incorrect ID assignments. The final participant sample consisted of 134 participants (62 females, 2 non-binary, mean age = 33.84 years, age range = 18 to 45 years).

**Experiment 2** In this experiment, *N* = 210 participants (100 females, 5 non-binary, mean age = 31.63 years, age range = 19 to 45) completed four blocks of the predator task (with differing variability and hazard-rate conditions) and four blocks of the probabilistic reversal learning task, with each block consisting of 80 trials. The predator-task blocks varied in variability and hazard rate, similar to those in experiment 1. Participants were compensated £8 for their participation in the study and had the opportunity to earn an additional 50 pence bonus for each block in which their score exceeded 550 points. Data from 5 participants were excluded for failing the questionnaire attention checks, and 16 participants were excluded for failing the initial headphone sound check. The final participant sample consisted of 189 participants (90 females, 5 non-binary, mean age = 31.53 years, age range = 19 to 45 years). In this final sample of 189 participants, 179 participants (87 females, 4 non-binary, mean age = 31.68 years, age range = 19 to 45 years) completed the reversal learning task, 175 participants (82 females, 5 non-binary, mean age = 31.37 years, age range = 19 to 45 years) completed the predator task, while 165 participants (79 females, 4 non-binary, mean age = 31.52 years, age range = 19 to 45 years) completed both tasks.

**Experiment 3** Following up on experiment 2, *N* = 103 participants (51 females, 2 non-binary, mean age = 32.06, age range = 18 to 45 years) completed four blocks of the predator task (with differing variability and hazard-rate conditions), along with four blocks of the probabilistic reversal learning task with reward magnitudes. Participants received £8 for their participation in the study, with additional bonuses based on performance: £3 for the top 5% of participants, £1 for the top 10%, and 25 pence for those in the top 50%. Data from 2 participants were excluded for failing the questionnaire attention checks, and 7 participants were excluded for failing the initial headphone sound check. The final participant sample consisted of 95 participants (47 females, 2 non-binary, mean age = 32.18, age range = 18 to 45 years). In this final sample of 95 participants, 94 participants (46 females, 2 non-binary, mean age = 32.1 years, age range = 18 to 45 years) completed the reversal learning task, 92 participants (44 females, 2 non-binary, mean age = 32.03 years, age range = 18 to 45 years) completed the predator task, while 91 participants (43 females, 2 non-binary, mean age = 31.95 years, age range = 18 to 45 years) completed both tasks.

**Experiment 4** In this experiment, *N* = 194 participants (96 females, 0 non-binary, mean age = 32.02 years, age range = 18 to 45) completed four blocks of the predator task (with differing variability and hazard-rate conditions) and four blocks of the reversal learning task, where outcome magnitudes were displayed above the fractals. The reversal learning task was divided into reward- and loss-domain blocks, where the order was counterbalanced. Participants received £8 for their participation in the study, with additional bonuses based on performance: £3 for the top 5% of participants, £1 for the top 10%, and 25 pence for those in the top 50%. Data from 6 participants were excluded due to failed questionnaire attention checks, 19 were excluded for failing the initial headphone sound check, and 1 subject was excluded from the reversal learning task for not making a selection in more than 15% of trials. The final participant sample consisted of 169 participants (85 females, 0 non-binary, mean age = 31.97 years, age range = 18 to 45 years). In this final sample of 169 participants, 161 participants (80 females, 0 non-binary, mean age = 31.87 years, age range = 18 to 45 years) completed the reversal learning task, 161 participants (81 females, 0 non-binary, mean age = 32.01 years, age range = 18 to 45 years) completed the predator task, while 153 participants (76 females, 0 non-binary, mean age = 31.91 years, age range = 18 to 45 years) completed both tasks.

**Experiment 5** In this experiment, *N* = 93 participants (48 females, 1 non-binary, mean age = 36.05 years, age range = 20 to 69 years) were recruited from Prolific to complete six blocks of the predator task, each consisting of 80 trials. The task featured a single variability and hazard-rate condition, and participants underwent minimal task training. Participants were compensated £9 for their participation in the study, with the opportunity to earn an additional 50 pence bonus for each block in which their score exceeded 450 points. Data from 2 participants were excluded for failing the questionnaire attention checks, and 15 participants were excluded for failing the headphone sound checks. The final participant sample consisted of 76 participants (36 females, 1 non-binary, mean age = 35.76 years, age range = 20 to 69 years).

**Experiment 6** In experiment 6, *N* = 97 participants (48 females, 1 non-binary, mean age = 36.71 years, age range = 18 to 69 years) were recruited from Prolific to complete 6 blocks of the predator task, with each block consisting of 65 trials. The task structure was similar to experiment 5, but all participants underwent comprehensive task training before starting the task. Participants were compensated £9 for their participation in the study, with the opportunity to earn an additional 50 pence bonus for each block in which their score exceeded 400 points. Data from 1 participant were excluded for failing the questionnaire attention checks, and 8 participants were excluded for failing the headphone sound checks. The final participant sample consisted of 89 participants (43 females, 1 non-binary, mean age = 35.5 years, age range = 18 to 69 years).

**Experiment 7** In this experiment, *N* = 35 participants (28 females, 1 non-binary, mean age = 25.91 years, age range = 20 to 41 years) were recruited from the general population in Berlin to participate in an in-person laboratory study involving the predator task with a single variability and hazard-rate condition. The skin conductance response was recorded throughout the task. Due to missing data or technical issues, data from 6 participants were excluded from the final analysis. The final participant sample consisted of 29 participants (22 females, 1 other, mean age = 25.76 years, age range = 20 to 41 years).

**Experiment 8** In the final experiment, *N* = 41 participants (33 females, 0 non-binary, mean age = 24.85 years, age range = 18 to 39 years) were recruited from the general population in Berlin to participate in an in-person study using a pilot version of the predator task. This version featured three distinct predators and was divided into blocks where screams or shocks served as aversive stimuli. Skin conductance responses were recorded throughout the task, and data from 1 participant were excluded from the data analysis due to technical malfunctions. The final participant sample consisted of 40 participants (32 females, 0 non-binary, mean age = 24.95 years, age range = 18 to 39 years).

### Predator Task

#### Task Model

For the purpose of computational modeling, we present a formal description of the predator task:

- 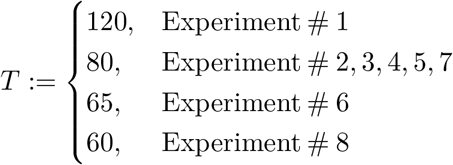 denotes the number of trials in a single block for the different experiments, indexed as *t* ∈ 𝒯 = {1, 2, …, *T*},
- *x* ∈ *X* = [0, 359]*°* denotes the set of predator attack directions,
- *µ* ∈ *M* = [0, 359]*°* is the set of the mean parameters of the Gaussian distribution that generates outcomes,
- *σ* ∈ *S* = {20, 30} *°* is the set of standard deviation parameters of the Gaussian distribution that generates the outcomes, where *σ* = 20 in the low-variability condition, and *σ* = 30 in the high-variability condition,
- *c*_*t*_ ∈ *C* = {0, 1}, *t* ∈ 𝒯 is the set of change points in the outcome contingencies, where *c*_*t*_ = 0 denotes that outcome contingencies are unchanged, and *c*_*t*_ = 1 indicates a change point,
- *r*_*t*_ ∈*R* = {−10, 0, 10}, *t* ∈ 𝒯 refers to the set of rewards in the task, where *r*_*t*_ = −10 for trials where the flame is not turned on, *r*_*t*_ = 0 for unsuccessful trials with the flame turned on and *r*_*t*_ = 10 for successful trials,
- *b*_*t*_ ∈ *B* = [0, 359]*°, t* ∈ 𝒯 denotes the set of participants’ flame positions on each trial, *h* ∈*H* = {0.1, 0.16} denotes the hazard rate that determines the frequency of change points, with *h* = 0.1 in the low-hazard-rate condition and *h* = 0.16 in the high-hazard-rate condition,
- *v*_*t*_ ∈ *V* = {−1, 1}, *t*∈ 𝒯 denotes the valence set, where *v*_*t*_ = 1 if the trial results in a successful defense (good outcome) and *v*_*t*_ = − 1 if the trial results in failure to save oneself (bad outcome),
- 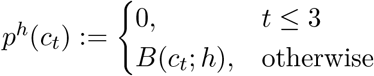 is the distribution that generates change points, i.e., change points do not occur for trials *t* ≤ 3 and in all other cases according to the Bernoulli distribution, which depends on the hazard rate *h*,
- 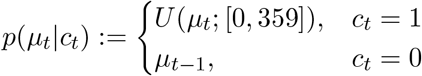 is the change-point-conditional distribution of *µ*_*t*_,
- 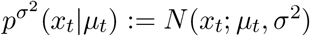 is the outcome-generating Gaussian distribution, where *x_t_ <* 0 and *x*_*t*_ *>* 359 were adjusted on the circular axis,
- 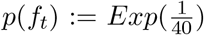 is the exponential distribution that generates the flame size *f*_*t*_ on each trial; the distribution is truncated to the range [15, 50] degrees.

#### Data preprocessing

Additional variables of interest for the analyses were participants’ estimation errors, prediction errors, prediction updates, and perseveration trials.

- *δ*_*t*_ ∈ *D* = [−180, 179]*°, t* ∈ 𝒯 refers to the set of prediction errors, defined as *δ*_*t*_ := circular(*x*_*t*_ − *b*_*t*_), where circular(·) is defined as the operation that ensures the result lies within [−180, 179]*°* to account for the periodic nature of angular measurements,
- *e*_*t*_ ∈ *E* = [0, 179]*°, t* ∈ 𝒯 is the set of estimation errors, defined as *e*_*t*_ := |circular(*µ*_*t*_ −*b*_*t*_) |, i.e., the absolute distance between the mean predator attack location and the prediction,
- *a*_*t*_ ∈ *A* = [−180, 179]*°, t* ∈ 𝒯 refers to the set of trial-by-trial prediction updates, which are defined as *a*_*t*_ := circular(*b*_*t*+1_ − *b*_*t*_),
- *n*_*t*_ ∈ *N* = {0, 1}, *t* ∈ 𝒯 refers to the set of perseveration trials, which are defined as 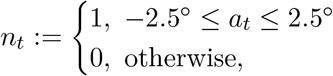 i.e., trials in which predictions were not updated were categorized as perseveration trials, with a tolerance of ± 2.5 degrees to account for minimal, potentially unintentional mouse movements caused by motor noise.

#### Reduced Bayesian model

The predator task can be modeled using Bayesian change-point-detection algorithms that predict future outcomes based on all previous outcomes (Adams & MacKay, 2007; Wilson et al., 2010). While theoretically robust, these algorithms require tracking all possible change-point combinations, posing computational challenges that may be unrealistic for human cognition. To address this, Nassar and colleagues proposed a reduced Bayesian model that approximates the solution of the full Bayesian model while remaining computationally feasible (Nassar et al., 2010, 2012; Wilson et al., 2010).

In the reduced Bayesian model, the learning rate is dynamically regulated by two parameters: relative uncertainty (RU) and change-point probability (CPP). RU quantifies the estimation uncertainty in the model’s belief about the predator’s attack location relative to environmental variability and estimation uncertainty. RU is high at the start (or after a known change point) and decreases as the model observes more trials and becomes more certain about the predator’s location. When a change point occurs, previously learned estimates are no longer predictive, requiring the model to relearn the new predator attack locations. Since the occurrence of a change point is unknown to the model, it infers it based on the computed CPP parameter, which estimates the probability of a change point based on the most recent prediction error. The learning rate of the reduced Bayesian model is a weighted combination of both RU and CPP. When CPP is high, the most recent outcome primarily determines the learning rate, while earlier outcomes are forgotten. Conversely, when CPP is low, learning is primarily driven by RU, and the model considers information from multiple past outcomes to predict the next predator location. We assume the following variables:

- *µ*_0_ := 180*°* is the model’s initial estimate of the *µ* parameter,
- 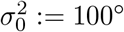 is the model’s initial estimation uncertainty over the *µ* parameter.

To infer the *µ*_*t*+1_ parameter, the reduced Bayesian model combines the latest outcome *x*_*t*_ and the previous prediction *µ*_*t*_. This combination is achieved using an error-driven sequential updating rule

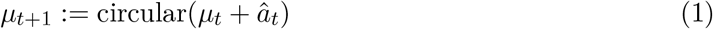

where

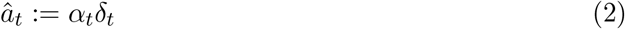

and where

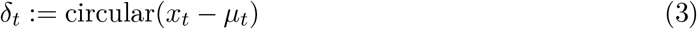

refers to the prediction error that expresses the difference between the last outcome *x*_*t*_ and the last prediction *µ*_*t*_. The weight of this combination is determined by the learning rate *α*_*t*_. The reduced Bayesian model uses a dynamical learning rate that depends on the two factors change-point probability *ω*_*t*_ and relative uncertainty *τ*_*t*_:

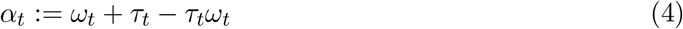

The change-point probability *ω*_*t*_ is computed as a function of the current prediction error *δ*_*t*_, the hazard rate of the task block *h* and the total uncertainty about the outcome 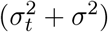,i.e.,

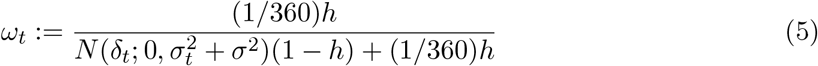

Next, to determine relative uncertainty *τ*_*t*+1_ for the next trial, the model first computes estimation uncertainty about the predator attack location, defined as

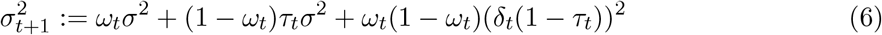

Subsequently, the model computes the fraction of estimation uncertainty relative to its total uncertainty about the next outcome, which is the sum of estimation uncertainty and the variability or risk in the environment *σ*^2^:

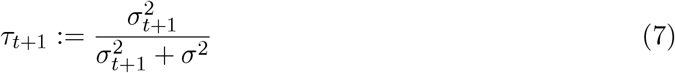

#### First-Level Regression Model

To understand the factors influencing trial-by-trial updates for each participant, we applied regression models incorporating both normative and non-normative parameters. This approach enabled us to assess the extent to which participants relied on an adaptive learning rate derived from the reduced Bayesian model and a fixed learning rate modeling a static influence of the prediction error on the belief update. To account for block-specific effects, the regression models also included coefficients for the variability and hazard-rate conditions, with the equation of the best-fitting model given as:

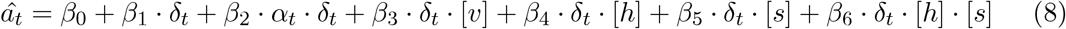

where *β*_0_ is the intercept. The *β*_1_ coefficient models the average effect of prediction error *δ*_*t*_ on the update and is referred to as the fixed learning rate. The *β*_2_ coefficient models the extent to which participants rely on an adaptive learning rate *α*_*t*_ derived from the reduced Bayesian model (eq. (4)). The *β*_3_ coefficient models the influence of valence on the updates, with the categorical variable *v* coded as 1 for a successful and 1 for an unsuccessful trial. *β*_4_ captures the influence of the hazard-rate level *h*, which is a categorical variable coded as *h* = − 1 for the high hazard-rate and *h* = − 1 for the low-hazard-rate condition. *β*_5_ captures the influence of the variability level *s*, which is a categorical variable coded as *s* = 1 for the high-variability and *s* = − 1 for the low-variability condition. The *β*_6_ coefficient captures the interaction of the variability and hazard-rate levels on the updates, accounting for how these factors jointly influence updates. That is, the inclusion of the hazard-rate level *h*, variability level *s*, and their interaction term accounts for differences in block types across experiments 1 to 4.

#### Evaluation and Estimation

We fitted the regression model to participant data. The probability of each trial-by-trial update was assumed to be sampled from a von Mises distribution centered on the model-predicted update *â*_*t*_, with concentration *κ*_*t*_ to capture noise in the updates

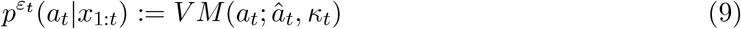

The concentration parameter *κ*_*t*_ is the inverse of the noise parameter *ε*_*t*_, expressed as:

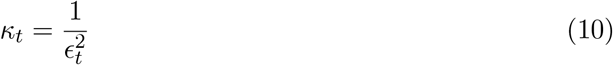

where *ϵ*_*t*_ is defined as:

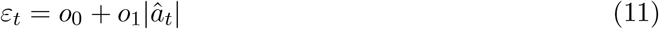

where *o*_0_ is the motor-noise component that models inaccuracies in flame placement that result from imprecision in the motor system (i.e., hand controlling the mouse in our case) for each participant. *o*_1_ is the learning-rate noise component and accounts for inaccuracies in flame placement that increase as a function of the update magnitude (i.e., larger updates tend to be noisier).

#### Model Comparison

To identify the model that best accounted for participants’ behavior, we compared Bayesian Information Criterion (BIC; Schwarz, 1978) scores summed across participants for several regression models. BIC was computed separately for each participant using the following equation:

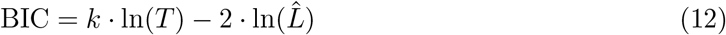

where *k* denotes the number of free parameters, *T* is the number of trials and 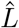 is the maximum likelihood of the model.

The candidate models included a model with only a fixed learning rate, a model with a fixed learning rate and valence component, a model incorporating both fixed and adaptive learning rates, and the full model specified in eq. (8), which includes fixed and adaptive learning rates as well as valence. All models accounted for hazard-rate and stochasticity levels. The model that achieved the lowest summed BIC score, indicating the best overall fit, included the fixed learning rate, adaptive learning rate, and valence parameters (Fig. S6).

#### Second-Level Regression With Factor Scores

To investigate how internalizing psychopathology influences learning behavior, we conducted a second-level robust regression analysis explaining the first-level regression parameters described above. This analysis incorporated participants’ factor scores as predictors while controlling for age and gender, and the regression equation corresponded to

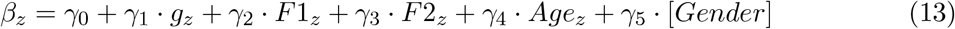

where *β*_*z*_ represents the *z*-scored parameter of interest from the first-level regression. The *γ*_0_ coefficient is the intercept; *γ*_1_, *γ*_2_, and *γ*_3_ capture the influence of the general factor G, anxiety-related factor F1, and depression-related factor F2, respectively. *γ*_4_ and *γ*_5_ account for the effects of age and gender. Gender was coded as 1 for males and − 1 for females, and all continuous predictors were *z*-scored to facilitate interpretability.

Bayes factors corresponding to *t*-statistics were computed in *R* using the ttest.tstat function from the BayesFactorpackage (Morey & Rouder, 2024), with the alternative hypothesis specified as a two-tailed Cauchy distribution with a medium scale parameter 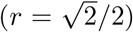.Multiple comparisons were corrected using the Benjamini–Hochberg false discovery rate (FDR) procedure (Benjamini and Hochberg, 1995; see False Discovery Rate Correction). Robust regression and FDR correction were performed in *Python* using the statsmodelslibrary.

### In-Person Study

#### Stimulus Calibration and Skin Conductance Response Measurements

In the in-person laboratory study, participants completed a version of the predator task that used either electrical shocks or screams as aversive stimuli. Electrical shocks were delivered as three 2-ms pulses, each separated by 15 ms, using a Digitimer DS5 (Digitimer Limited, United Kingdom) isolated bipolar constant current stimulator. The DS5 received an analog voltage input from the presentation computer via a National Instruments Data Acquisition System (National Instruments, USA), which was converted into isolated constant-current electrical shocks. These shocks were administered through a Digitimer wasp stimulation electrode attached to the participant’s lower shin, delivering a sharp and uncomfortable stimulus. Skin conductance responses (SCR) were recorded using two electrodes attached to the index and middle fingertips of the left hand (Gershman & Hartley, 2015), a position particularly suitable for SCR recording (Ojala & Bach, 2020). The SCR data were recorded through a 16-channel BrainAmp ExG amplifier (Brain Products GmbH, Germany) with the BrainVision Recorder software.

The SCR data, along with task triggers for trial and outcome onset, were stored in the BrainVision core data format (CDF) and preprocessed in Python using the MNE-Python package (Gramfort et al., 2013). SCR preprocessing steps included applying a first-order Butterworth low-pass filter with a cutoff frequency of 5 Hz, downsampling the data from 5000 Hz to 100 Hz, *z*-scoring, and further downsampling to 10 Hz (Bach et al., 2009; Gerster et al., 2018). Task events were epoched from 2 seconds before to 9 seconds after outcome presentation, with baseline correction applied using a 0.5-second window before outcome onset. To prepare the SCR data for analysis, all participants’ data were combined and averaged across epochs for each combination of timestamps, participants, and task-relevant variables.

To cater to individual differences in shock perception across participants, the pulse amplitude was individualized using a staircase procedure (Aylward et al., 2019; Wise et al., 2019). The procedure began with a pulse amplitude of 100 mV, which was increased in 100 mV increments. At each step, participants rated the perceived intensity of shocks on a scale from 1 (minimal pain) to 10 (maximum tolerable pain for the task). This process continued until a rating of 10 was achieved or the maximum amplitude of 3.5 V was reached. This procedure was repeated three times, and the average amplitude corresponding to the rating of 10 across the three runs was calculated. The shock intensity for the task was set at 80% of this average. Finally, participants received seven test shocks at this chosen intensity, delivered at random intervals ranging from 3 to 7 s, to confirm that the intensity was tolerable.

Aversive screams were delivered using headphones, and a similar staircase procedure was used to calibrate the sound intensity for each participant in experiment 7. Here, participants rated the volume intensity of the screams from 1 (minimum sound intensity) to 10 (maximum tolerable sound intensity), while the volume level increased by 3% on each step, with the procedure repeated three times. The average sound intensity corresponding to the rating of 10 across the three runs was calculated, with the final sound intensity chosen as 80% of this maximum value. Participants then received seven test screams at this volume, delivered at random intervals, to ensure tolerability.

#### Cluster-Based Permutation Testing

To control for multiple comparisons in analyzing skin conductance responses across time bins, we applied cluster-based permutation testing (Maris, 2012; Maris & Oostenveld, 2007). For each time bin, we computed the test statistic (*t*-value) by comparing conditions of interest in two-tailed tests or by comparing one set of values to zero in one-tailed tests. Adjacent time bins with *t*-values exceeding a predefined threshold (corresponding to *α* = 0.05) were then grouped into clusters, and the cluster mass was calculated as the sum of *t*-values within each cluster. Next, to assess the significance of the observed cluster masses, we created a null distribution by randomly shuffling condition labels within participants to generate pseudo-datasets. For each pseudo-dataset, we recalculated the *t*-values, identified clusters surpassing the threshold, and recorded the maximum cluster mass. This permutation process was repeated 1,000 times to construct a null distribution of cluster masses. Finally, we compared the observed cluster masses to this null distribution to calculate *p*-values, representing the probability of observing a cluster mass as large as or larger than the observed one under the null hypothesis.

#### Probabilistic Reversal Learning Task Task Model

The formal model of the probabilistic reversal learning task is as follows:

- *f* ∈ *F* = {1, 2} denotes the set of fractals, where *f* = 1 corresponds to the fractal 1 and *f* = 2 corresponds to fractal 2,
- *B* := 4 denotes the number of blocks in the task, indexed as *b* ∈ ℬ = {1, 2, …, *B*},
- 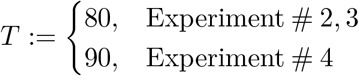 represents the number of trials per block, indexed as *t* ∈ 𝒯 = {1, 2, …, *T*},
- *a*_*t*_ ∈ *A* = {1, 2}, *t* ∈ *𝒯* is the set of actions, where *a*_*t*_ = 1 if fractal 1 is chosen and *a*_*t*_ = 2 if fractal 2 is chosen on the current trial.
- *u* ∈*U* = {−1, 1} is the set of task phases, where *u* = −1 corresponds to the stable phase and *u* = 1 corresponds to the volatile phase; each phase comprises two consecutive blocks in the task version with reward magnitudes and one block in the task version with reward and loss domains,
- *o*∈ *p*(*o*) := *B*(0.5) denotes the distribution that determines the order of phase presentation for each participant; *o* = 0 represents the stable phase followed by the volatile phase, while *o* = 1 represents the volatile phase followed by the stable phase,
- *p*(*f*) := *B*(0.5) is the distribution that selects whether fractal 0 or fractal 1 will be the dominant fractal in each block,
- *r*_*u*_∈ *R* = {0.75, 0.8}, *u* ∈*U* refers to the set of the dominant fractals’ reward probabilities, where *r*_*u*_ = 0.75 for the stable phase and *r*_*u*_ = 0.8 for the volatile phase,
- *p*(*x*_*t*_ | *r*_*u*_) := *B*(*r*_*u*_) defines the outcome-generating distribution for the dominant fractal on each trial,
- *p*(*l*) := *B*(0.5) is the distribution that determines whether fractal 1 or fractal 2 appears on the left side of the screen on each trial,
- *x*_*t*_∈ *X* = {0, 1}, *t* ∈ 𝒯 is the set of outcomes, where *x*_*t*_− _1_ = 1 if fractal 2 is chosen and followed by a reward or fractal 1 is chosen and followed by no reward; *x*_*t*_ − _1_ = 0 encodes the opposite case,
- *v*_*t*_ ∈ *V* = {−1, 1}, *t* ∈ 𝒯 refers to the set of valence, where *v*_*t*_ = 1 if the trial results in a reward (good outcome) and *v*_*t*_ = −1 if the trial results in no reward (bad outcome).

#### Model Space

The participant behavior on the binary reversal learning task was modeled using different versions of the canonical Rescorla-Wagner (RW) model. Each model’s estimated choice probabilities were based on a learning-rate parameter and an inverse temperature parameter capturing choice stochasticity. Following the approach of Gagne et al. (2020), we sub-divided parameters into components that could capture different task and behavioral dynamics. Model fits for the different model versions were compared using Pareto-smoothed importance-sampling-based approximation of leave-one-out cross-validation (PSIS-LOO), and the model with the lowest PSIS-LOO value was chosen as the best-fitting model.

#### Model 1: Canonical Rescorla-Wagner model

The first model that we fitted to the data was the canonical Rescorla-Wagner model, which includes learning-rate and inverse temperature parameters. We applied the model to both phases of the task by incorporating phase-specific parameters for learning rate (*α*_*V olatile*_ − _*stable*_) and inverse temperature (*B*_*V olatile*_ − _*stable*_). *α*_*V olatile*_− _*stable*_ captures the relative difference in learning rate between volatile and stable phase, whereas *B*_*V olatile*_ − _*stable*_ captures the relative difference in choice stochasticity between the two phases. The formal model is expressed as:

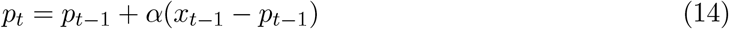

where *p*_*t*_ is the probability estimate of fractal 2. The learning rate *α* is defined as:

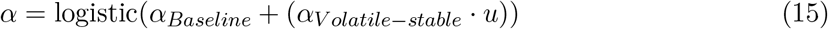

where *u* represents the task phase and the logistic function constrains the learning rate between 0 and 1. The model relies on a softmax action-selection rule, where the probability of choosing fractal 2 is given by:

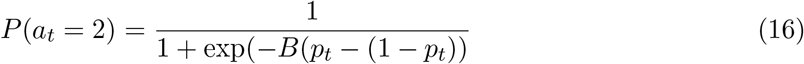

where *a*_*t*_ represents the fractal chosen, while the inverse temperature parameter *B* is a combination of a baseline component and a task-specific component, and is transformed using an exponential function to ensure a positive value:

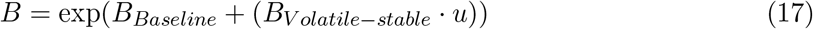

#### Model 2: Incorporating valence into learning

In model 2, we extended the canonical Rescorla-Wagner model by incorporating the valence of outcomes, allowing us to quantify how participants learn from rewards versus punishments.

Valence describes whether the trial resulted in a reward (*v*_*t*_ = 1) or no reward (*v*_*t*_ = − 1). In this model, the learning rate had an additional valence component *α*_*Good*_− _*bad*_ that adjusts the learning rate accordingly. The updated learning rate in this model is defined as:

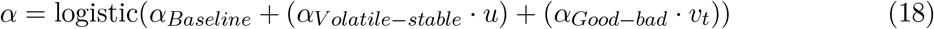

#### Model 3: Incorporating valence into choice stochasticity

To determine whether outcome valence influences choice stochasticity, we extended model 2 by adding a valence-specific parameter to the inverse temperature of the softmax function. This parameter captures whether participants’ choice consistency changes after experiencing good or bad outcomes. The inverse temperature parameter for this extended model is defined as:

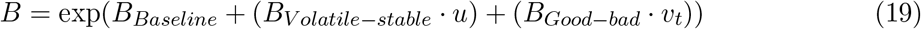

#### Model 4: Incorporating interaction of valence and task phase into learning

To investigate whether learning from good versus bad outcomes varied across different task phases, we extended the learning-rate parameter of model 3 by introducing an interaction parameter between valence and task phase. That way, the model can assess whether the impact of good or bad outcomes on the learning rate differs depending on whether the environment is stable or volatile:

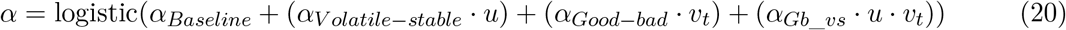

#### Model 5: Incorporating interaction of valence and task phase into choice stochasticity

We further extended model 4 by introducing an interaction term between valence and task phase into the inverse temperature parameter. This addition allowed us to explore whether participants made more stochastic choices in the stable or volatile phase depending on outcome valence. The inverse temperature parameter of this model is given as:

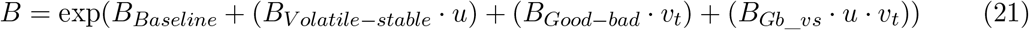

#### Model 6: Adding a choice kernel

To account for the possibility that participants may repeat their previous choices independently of the associated reward probabilities, we extended model 5 by integrating a choice kernel *k*_*t*_. The choice kernel tracks the tendency to choose a specific fractal based on the recent choice history, where the choice-kernel learning rate *α*_*ck*_ determines the influence of past choices on the current decision. The updating rule of the choice kernel is given as:

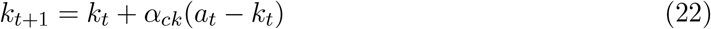

where *a*_*t*_ represents the fractal chosen on the previous trial.

This choice kernel is then integrated into the decision-making process through an inverse choice-kernel temperature parameter *B*_*ck*_ as follows:

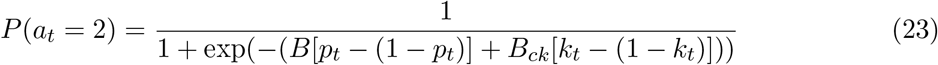

where *p*_*t*_ represents the probability of choosing fractal 2.

#### Model 7: Independent probability estimates for each fractal

In this model, we explored the possibility that participants learn independent reward probabilities for each fractal, despite the actual reward probabilities for both fractals being yoked together. To account for this, we extended model 5 by introducing separate Rescorla-Wagner update equations for both fractals, updating only the chosen fractal on each trial. In this model, 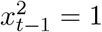 if fractal 2 is chosen and followed by a reward, whereas 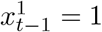 if fractal 1 is chosen and followed by a reward; otherwise, these variables are set to 0. The update equation for the reward probability estimate of each fractal is given as:

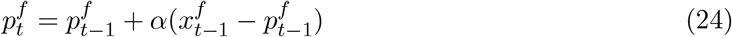

where *f* denotes the fractal (1 or 2), and 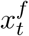 is the observed outcome (1 for a reward, 0 otherwise) for the corresponding fractal. Following each update, a decay parameter *δ* controls the amount of decay to 0.5 for both estimates, where a higher *δ* leads to a stronger pull towards 0.5:

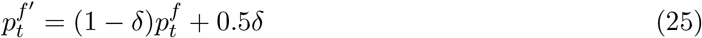

The decay parameter itself is dynamically adjusted based on both a baseline component and a task-phase component, reflecting different levels of decay depending on whether the task is in a stable or volatile phase:

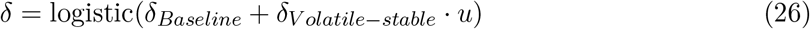

The probability of choosing fractal 2 is determined using the softmax function:

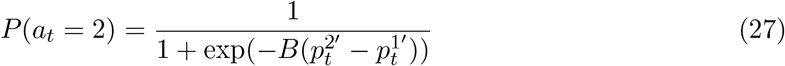

#### Model 8: Combining a choice kernel and independent probability estimates

We extended model 7 by incorporating a choice kernel to account for the possibility that participants may have a tendency to repeat their previous choices, in addition to maintaining separate reward probability estimates for each fractal. The choice-kernel update and decision rule are applied as described in eqs. (22) and (23).

#### Hierarchical model estimation

To incorporate the effects of the general factor and the two sub-factors into the model, we employed a hierarchical Bayesian approach based on Gagne et al. (2020). This approach allowed us to estimate distributions over individual-level Rescorla-Wager model parameters using population-level parameters, where participant scores on the three factors were incorporated at the population level for all models. Consequently, we were able to capture the variance in behavior attributable to the three factors of interest.

Specifically, each individual-level parameter was assigned a population-level normal distribution shared across participants. The mean *µ* of this population distribution was specified by a linear model estimated separately for each parameter

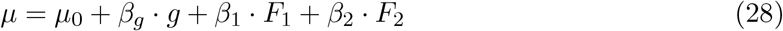

where *µ*_0_ represents the overall mean of the parameter across all participants, while the weights *β*_*g*_, *β*_1_, *β*_2_ encapsulate the deviation in the parameter value from the overall mean based on each participant’s score on the three factors. The variance *σ*^2^ of the population-level distribution for each parameter was also estimated separately.

The hyperpriors for the population-level parameters *µ*_0_, *β*_*g*_, *β*_1_, *β*_2_ were specified as uninformative normal distributions with mean = 0 and standard deviation = 10, while the hyperpriors for the population-level variance *σ*^2^ followed HalfCauchy distributions with *γ* = 2.5. Models were fitted using PyMC3(Abril-Pla et al., 2023) in *Python*. We applied a Hamiltonian Monte-Carlo method with 4 chains to sample from the full posterior, where each chain had 1200 tuning steps and 2000 samples. Convergence was assessed by visual inspection of traces and by analyzing the Gelman-Rubin statistics (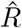;Gelman and Rubin, 1992), where 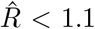 indicates good convergence.

#### Model comparison

For the probabilistic reversal learning task, we evaluated fits of different models using PSIS-based computation of LOO-CV (Gagne et al., 2020; Vehtari et al., 2017). While traditional LOO-CV methods are effective for comparing Bayesian models based on their out-of-sample prediction accuracy, they are computationally expensive. PSIS-LOO offers a more efficient approximation of LOO-CV accuracy, delivering performance comparable to the widely applicable information criterion (WAIC; Watanabe, 2013). The model with the lowest value of PSIS-LOO was chosen as the best-fitting model for the data.

#### Parameter Recovery

To assess the identifiability of the winning model’s parameters, we conducted a parameter recovery analysis. We first simulated choice data for five datasets using the subject-specific posterior means of all parameter components from the winning model. The model was then fitted to these simulated datasets, and Spearman’s rank correlation was computed between the original (ground truth) and estimated (recovered) parameter values for each dataset. Fig. S16 shows parameter recovery for one of these simulated datasets.

To further assess parameter identifiability across a wider parameter range, we generated an additional dataset with 500 simulated subjects, where learning-rate parameters were sampled from a uniform distribution (*U* (− 2, 2)). The winning model was then fitted to this dataset, and Spearman’s rank correlation was calculated between the ground truth and recovered parameters (see SM Parameter Recovery).

#### Binary Volatile Kalman Filter

As an additional analysis, we fitted a binary volatile Kalman filter (VKF; Piray and Daw, 2020) to participant data from the reversal learning task. Kalman filters are widely used to infer hidden environmental states, especially when the environment is gradually changing. The binary VKF is specifically designed for environments with binary observations and changing volatility. It tracks environmental volatility on each trial, which directly influences the learning rate in the update rule (for details, see Piray and Daw, 2020).

The Kalman gain *k*_*t*_ on each trial is given as:

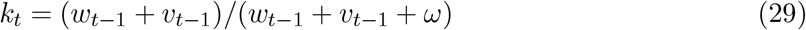

where *k*_*t*_ depends on the posterior variance on each trial *w*_*t*_, the inferred volatility on each trial *v*_*t*_, and a noise parameter *ω*.

The learning rate *α*_*t*_ on the current trial is given as:

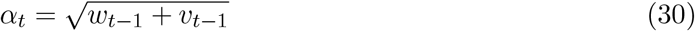

which is then used to update the posterior mean *m*:

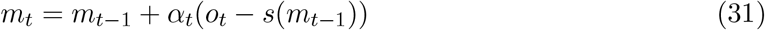

where s(·) is the sigmoid function and *o*_*t*_ is the observed outcome. On every trial, the posterior variance also gets updated depending on the Kalman gain and the volatility estimate:

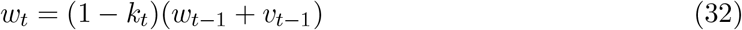

with the autocovariance between consecutive states given as:

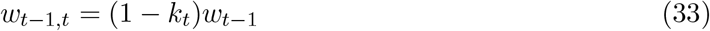

The volatility *v*_*t*_ is tracked using a volatility-update-rate parameter *λ*, where higher *λ* values allow for a higher speed of update:

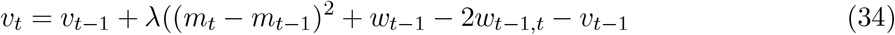

To model choice behavior, the probability of selecting fractal 2 (*a*_*t*_ = 2) was computed using a softmax function with an inverse temperature parameter *B*, based on the difference in estimated reward probabilities *c*_*t*_:

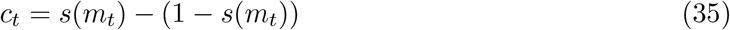

where *s*(*m*_*t*_) rescales the estimate to the unit range, representing the inferred probability of reward for fractal 2 according to

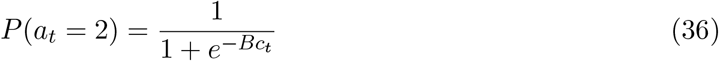

This model, when fitted to participant data, has 4 free parameters: the volatility update rate *λ*, the initial estimate of volatility *v*_0_, the noise parameter *ω*, and the inverse temperature parameter *B*. We estimated the free parameters using the L-BFGS-B constrained minimization algorithm from the SciPy library in *Python*, with values of VKF parameters constrained using priors informed by moment-matching. To ensure robust parameter estimation, we repeated the estimation 30 times with randomly initialized starting values and selected the parameters corresponding to the iteration that achieved the lowest negative log-likelihood.

#### False Discovery Rate Correction

To control for multiple comparisons, we applied the Benjamini-Hochberg false discovery rate (FDR) procedure (Benjamini & Hochberg, 1995). This method controls the expected proportion of false positives (Type I errors) among rejected hypotheses, offering a less conservative alternative to other multiple correction methods such as the Bonferroni correction, which controls the family-wise error rate (FWER). In the Benjamini-Hochberg procedure, the observed *p*-values are first sorted in ascending order:

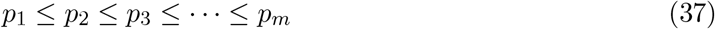

where *p*_*i*_ denotes the *i*-th smallest *p*-value out of the *m* total tests. Let *k* then be the largest index for which:

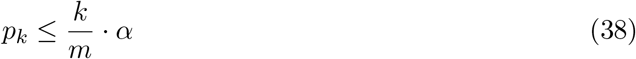

where *m* is the total number of tests, and *α* = 0.05 is the FDR threshold. All hypotheses corresponding to *p*-values *p*_1_, *p*_2_, …, *p*_*k*_ are subsequently rejected. The adjusted *p*-values are then computed using:

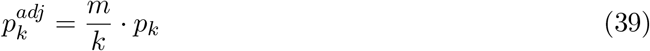

followed by a monotonicity correction to ensure that the adjusted *p*-values are non-decreasing across ranks:

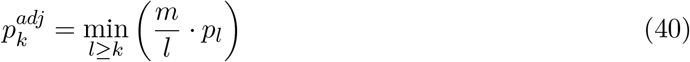

We used the fdrcorrectionfunction from the statsmodel.statsmodule in *Python* to implement FDR correction.

#### Software and Data Repository

The data and code for analysis are available at: https://github.com/Hashimsat/Internalizing_AnalysisCode.

The predator task and reversal learning tasks were developed in *Javascript* using the Phaser 3game development framework (https://phaser.io). Data analysis and computational modeling were conducted in *Python* (Python Software Foundation; https://www.python.org/) and *R* (R Core Team, 2021). We used the PyMC3 (Abril-Pla et al., 2023), pandas(McKinney, 2010), NumPy (Harris et al., 2020), SciPy(Virtanen et al., 2020), statsmodels(Seabold & Perktold, 2010), seaborn (Waskom, 2021), matplotlib(Hunter, 2007), tqdm(https://doi.org/10.5281/zenodo.1239851) and PIL (https://pillow.readthedocs.io/en/stable/index.html) libraries for *Python*-based analysis. Bayes factors were computed in *R* using the BayesFactor package (Morey & Rouder, 2024).

## Acknowledgments

We thank Sonia J. Bishop, Ulrike Lüken, Claire M. Gillan, Toby Wise, Ondrej Zika, and Ryszard Auksztulewicz for their helpful comments and for providing useful resources. We also thank Hauke Heekeren for his kind support over the years. M.H.S. was supported by the German Federal Ministry of Education and Research (BMBF) and the Max Planck School of Cognition, Leipzig, Germany. M.R.N. was supported by a Humboldt Research Fellowship for Experienced Researchers. R.M.C. was supported by the European Research Council (ERC) Consolidator grant (ERC-CoG-2024101123101). N.W.S. was supported by a Starting Grant from the European Union (ERC StG REPLAY-852669) and the Excellence Strategy of the Federal Government and the Länder. P.D. was supported by the Max Planck Society and the Humboldt Foundation. R.B. was supported by Deutsche Forschungsgemeinschaft (DFG, German Research Foundation), grant numbers BR 6959/2-1 and GL 984/3-1.

## Supplementary Materials

### Factor Analysis

We conducted a factor analysis using data from *N* = 748 participants. To verify the suitability of the questionnaire data for factor analysis, we employed the Kaiser-Meyer-Olkin (KMO) measure of sampling adequacy (Kaiser & Rice, 1974) and Bartlett’s test of sphericity (BST, Bartlett, 1951). The KMO assesses the proportion of variance that may be attributed to common variance among variables, with higher values indicating greater suitability for factor analysis. Our dataset achieved a total KMO value of 0.955, confirming its adequacy for factor analysis. This result was further supported by the BST results (*χ*^2^ = 50350.3, *p <* 0.001), demonstrating that the correlation matrix is not an identity matrix and is, therefore, appropriate for factor analysis.

We determined the number of factors by combining theoretical considerations, visual inspection of the scree plot, and parallel analysis. Parallel analysis compares the eigenvalues of the observed dataset with those derived from either a dataset generated by sampling from a Gaussian distribution or a resampled version of the original dataset with the same dimensions. This approach helps identify factors with eigenvalues that are higher than those expected from random noise, providing candidates for factor selection. However, parallel analysis using a randomly generated normal dataset can overestimate the number of factors, particularly in large samples, as the random eigenvalues tend to converge around 1 (Hayton et al., 2004). This can result in many eigenvalues from the observed data exceeding the threshold, suggesting an unrealistic number of factors. To mitigate this, we combined theoretical insights, the scree plot’s inflection point, and eigenvalues from the resampled dataset to refine our factor selection.

For our dataset, parallel analysis using the random normal dataset suggested 48 factors—an implausibly high number (Fig. S1). In contrast, parallel analysis using the resampled dataset identified a three-factor structure, which aligned with theoretical expectations and previous literature (Clark & Watson, 1991; Gagne et al., 2020; Simms et al., 2008). Thus, we adopted this three-factor model for further analysis.

**Figure S1.**
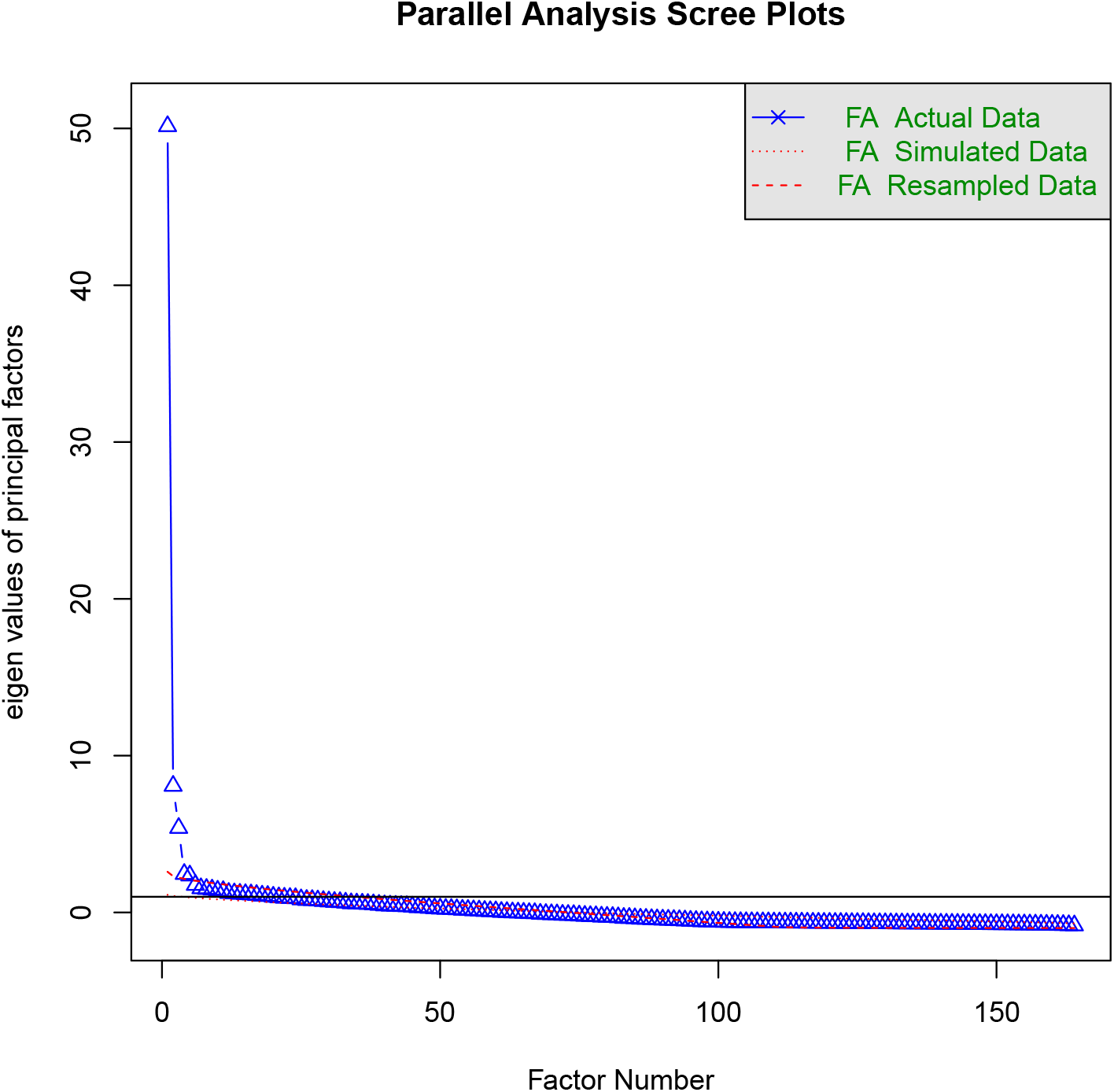
Scree plot of eigenvalues and parallel analysis results. The plot shows the eigenvalues of our questionnaire data, ordered in descending magnitude. The red dotted lines represent thresholds derived from parallel analyses, with a factor considered viable if its corresponding eigenvalue lies above the threshold. Using a randomly generated normal dataset, the parallel analysis yields eigenvalue thresholds close to 1 due to the large sample size, suggesting an unrealistic number of 48 factors. In contrast, parallel analysis with resampled data provides higher thresholds, identifying only three eigenvalues above the threshold. This supports a three-factor structure consistent with prior research (Clark & Watson, 1991; Gagne et al., 2020).

### Comparing factor structure with Gagne et al. (2020)

Gagne et al. (2020) performed factor analysis on a clinical dataset (*N* = 86) and validated the factor structure using an independent online sample (*N* = 199). We compared the factor loadings and scores from our analysis with those reported by Gagne et al. (2020), focusing on common questionnaire items. Our loadings were highly congruent with those from the clinical dataset for the general factor (cosine similarity = 0.97) and the depression-related factor (cosine similarity = 0.80) and showed moderate congruence for the anxiety-related factor (cosine similarity = 0.47). Comparisons with the independent online sample from Gagne et al. (2020) indicated strong congruence across all factors, with cosine similarities of 0.97 (general factor), 0.88 (depression-related factor), and 0.90 (anxiety-related factor).

As a control analysis, we applied the factor loadings from the clinical and online samples of Gagne et al. (2020) to our dataset to compute factor scores for all participants, and then compared these scores with our own factor scores (Fig. S2). The general-factor scores showed large correlations with those derived from both the clinical loadings (*r* = 0.71) and the online loadings (*r* = 0.93). Similarly, the depression-related factor scores exhibited large correlations with scores based on the clinical loadings (*r* = 0.75) and the online loadings (*r* = 0.89). However, for the anxiety-related factor, correlations were small for the clinical dataset (*r* = -0.07) but large for the online dataset (*r* = 0.9) from Gagne et al. (2020). These findings highlight the strong congruence between our factor structure and that of Gagne et al. (2020), particularly for the general and depression-related factors. The discrepancy in the anxiety-related factor, showing alignment with the online sample but not the clinical sample, suggests that clinical populations may exhibit distinct latent anxiety-related constructs compared to the general population.

**Figure S2.**
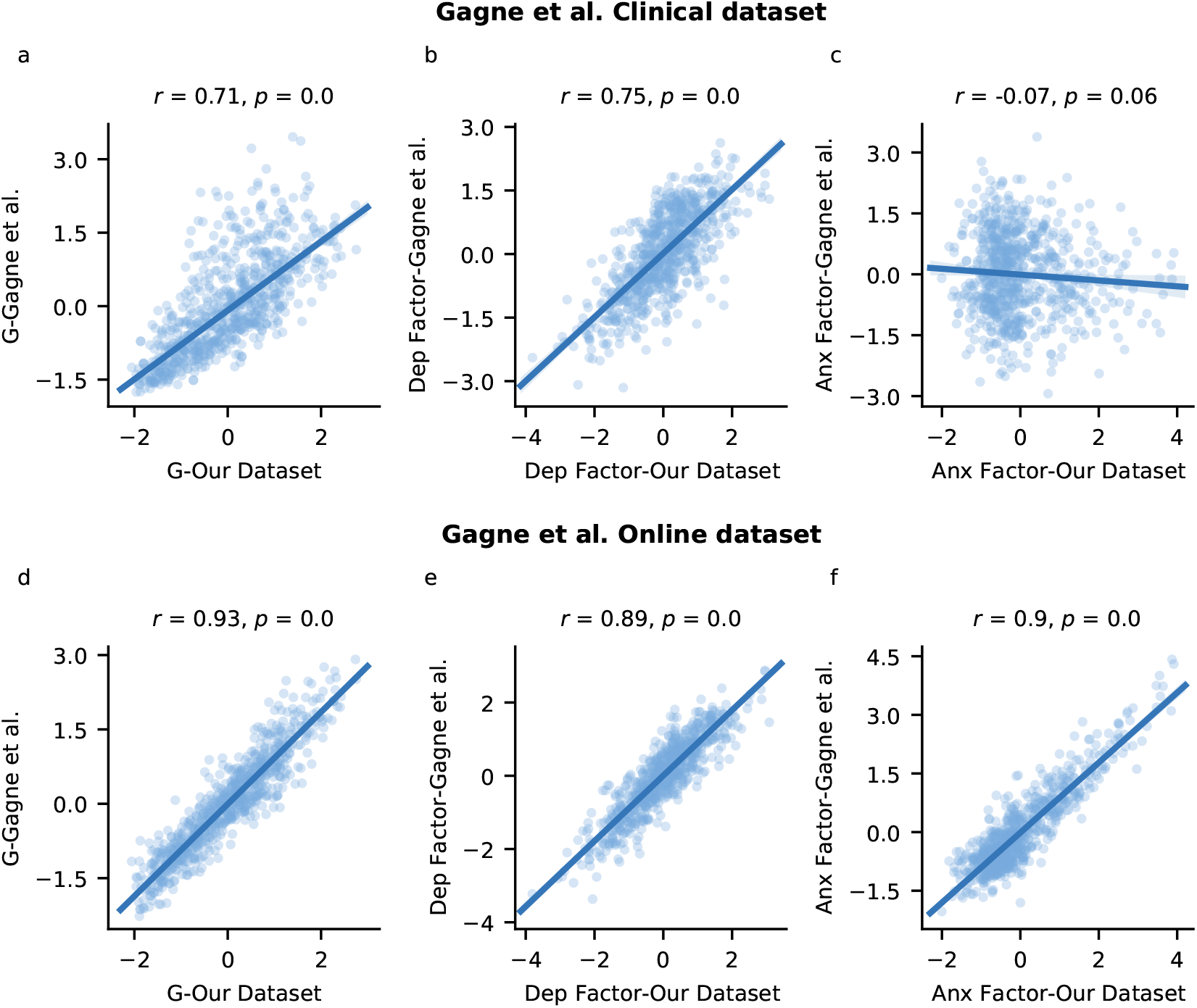
Comparison of factor scores from our analysis with those derived using loadings from Gagne et al. (2020). G represents the internalizing factor, Dep represents the depression-related factor, and Anx represents the anxiety-related factor. **a-c**| Correlation of our factor scores (x-axes) with scores derived using loadings from the clinical dataset (y-axes): **a**| General-factor scores exhibit large correlations. **b**| Depression-related scores also exhibit strong correlations. **c**| The anxiety-related factor shows a negligible association with the scores derived using the clinical dataset. **d-e**| Correlation of our factor scores (x-axes) with scores derived using loadings from the independent online sample (y-axes). **d**| General-factor scores reveal large correlations. **e**| Similarly, we observed large correlations for the depression-related factor. **f**| The anxiety-related scores show a large association with the scores derived using the online dataset.

**Figure S3.**
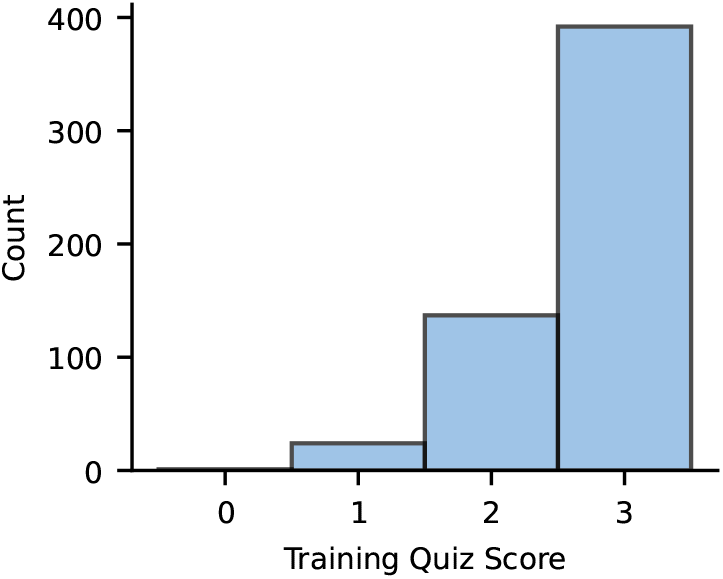
Participant performance in the training quiz for the predator-task version with different conditions (*N* = 554). Participants demonstrated a good understanding of the predator task, with 392 participants getting all three answers correct, 137 participants getting two answers correct, and 24 participants getting only 1 answer correct. Only 1 participant failed to answer any question correctly.

### Predator Task

#### Training Quiz

After completing the initial training phase of the predator task, participants were assessed on their understanding of the task through a brief quiz consisting of three multiple-choice questions. The questions were: (1) “When should you turn the fire on?” (2) “Where should you place the fire after turning it on?” and (3) “How do you get more points in the game?” Each question offered four answer options. After submitting their answers, participants received immediate feedback on their responses, along with a concise explanation of the correct answers to reinforce their understanding of the task.

#### Task Version with Different Variability and Hazard-Rate Conditions

In experiments 1-4, a total of *N* = 563 participants (demographic details in Table S1) completed a predator-task version with different variability and hazard-rate conditions. Nine participants who identified their gender as “non-binary” were excluded due to insufficient sample size for appropriately controlling for gender, resulting in a final sample of *N* = 554 participants. Before starting the task, participants received training and completed a training quiz to assess their understanding. Analysis of the quiz data showed that participants demonstrated a good understanding of the task: 392 participants answered all three questions correctly, 137 answered two questions correctly, 24 answered one question correctly, and only one participant failed to answer any question correctly (Fig. S3).

#### Descriptive Results

Next, we analyzed estimation errors across the different variability and hazard-rate conditions. Participants had higher estimation errors in the high-variability condition (median = 36.26, IQR 32.93 to 40.7) compared to the low-variability condition (median = 29.58, IQR 26.74 to 33.75, paired-sample *t*-test *t*_553_ = -24.27, *p* < 0.001, *BF*_01_ = 0). Similarly, participants had higher estimation errors in the high-hazard-rate condition (median = 35.72, IQR 32.69 to 40.32) compared to the low-hazard-rate condition (media = 30.36, IQR 27.23 to 34.21, *t*_553_ = -21.05, *p* < 0.001, *BF*_01_ = 0). These results suggest an effective implementation of the variability and hazard-rate manipulations in this task version. Importantly, the results were consistent when analyzing estimation errors across conditions separately for each of the four experiments (Fig. S4), where estimation errors were consistently lower in the low-variability and the low-hazard-rate condition compared to their respective counterparts.

Next, we examined whether internalizing-related differences in estimation errors and empirical learning rates emerged across the different variability and hazard-rate conditions (Fig. S5). We did not find any significant differences in estimation errors between the low- and high-internalizing groups in any condition (Fig. S5 a-d). Likewise, regression analyses of estimation errors against internalizing, while controlling for age and gender, did not reveal a significant association in any of these conditions (Fig. S5 e-h). Similarly, empirical learning rates were not observed to be significantly different between internalizing groups (Fig. S5 i-l) and were not found to be significantly associated with internalizing scores in any of the conditions (Fig. S5 m-p). These results suggest that while the different variability and hazard-rate conditions do have an influence on the overall performance across participants, they are not associated with any internalizing-related learning impairments.

**Table S1.** Basic demographic details of participants who completed the predator task version with different variability and hazard-rate conditions. STAI-Y1: Spielberger State-Trait Anxiety Inventory - state scale; STAI-Y2: Spielberger State-Trait Anxiety Inventory - trait scale; STICSA-T: State-Trait Inventory for Cognitive and Somatic Anxiety - trait scale, IUS-27: Intolerance of Uncertainty Scale; BDI: Beck’s Depression Inventory; PSWQ: Penn State Worry Questionnaire; MASQ: Mood and Anxiety Symptom Questionnaire; MASQ-Anh: Mood and Anxiety Symptom Questionnaire - anhedonia subscale; MASQ-AnxAr: Mood and Anxiety Symptom Questionnaire - anxious arousal subscale

| Measure | Value |
| --- | --- |
| Number of Participants (N) | 563.0 |
| Females (N) | 269.0 |
| Non-binary (N) | 9.0 |
| Age (mean ± SD) | 32.27 ± 6.85 |
| STAI-Y1 (mean ± SD) | 42.71 ± 13.02 |
| STAI-Y2 (mean ± SD) | 50.24 ± 12.98 |
| STICSA-T (mean ± SD) | 40.18 ± 13.42 |
| BDI (mean ± SD) | 19.44 ± 14.27 |
| PSWQ (mean ± SD) | 54.36 ± 14.53 |
| IUS-27 (mean ± SD) | 77.82 ± 23.33 |
| MASQ-Anh (mean ± SD) | 69.41 ± 18.31 |
| MASQ-AnxAr (mean ± SD) | 28.53 ± 10.67 |
| General Factor (mean ± SD) | 0.04 ± 1.0 |
| Anxiety-Specific Factor (F1) (mean ± SD) | -0.01 ± 0.98 |
| Depression-Specific Factor (F2) (mean ± SD) | 0.03 ± 1.0 |

**Figure S4.**
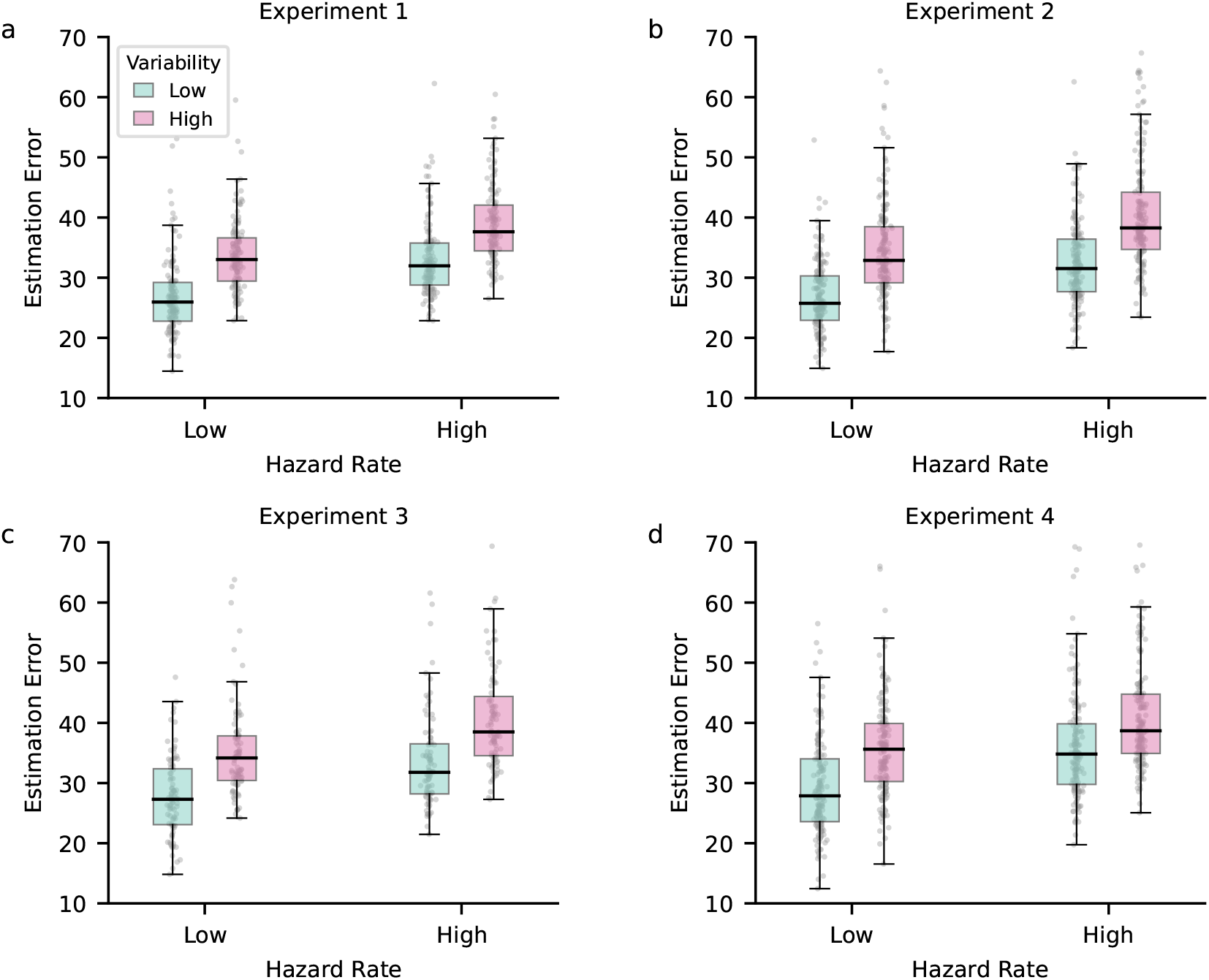
Estimation errors across variability and hazard-rate conditions in the predator task. Across all four experiments using the predator task with differing variability and hazard-rate conditions (**a-d**), participants exhibited lower estimation errors in low-variability conditions compared to high-variability conditions. Similarly, high-hazard-rate conditions consistently led to higher estimation errors compared to low-hazard-rate conditions across tasks.

**Figure S5.**
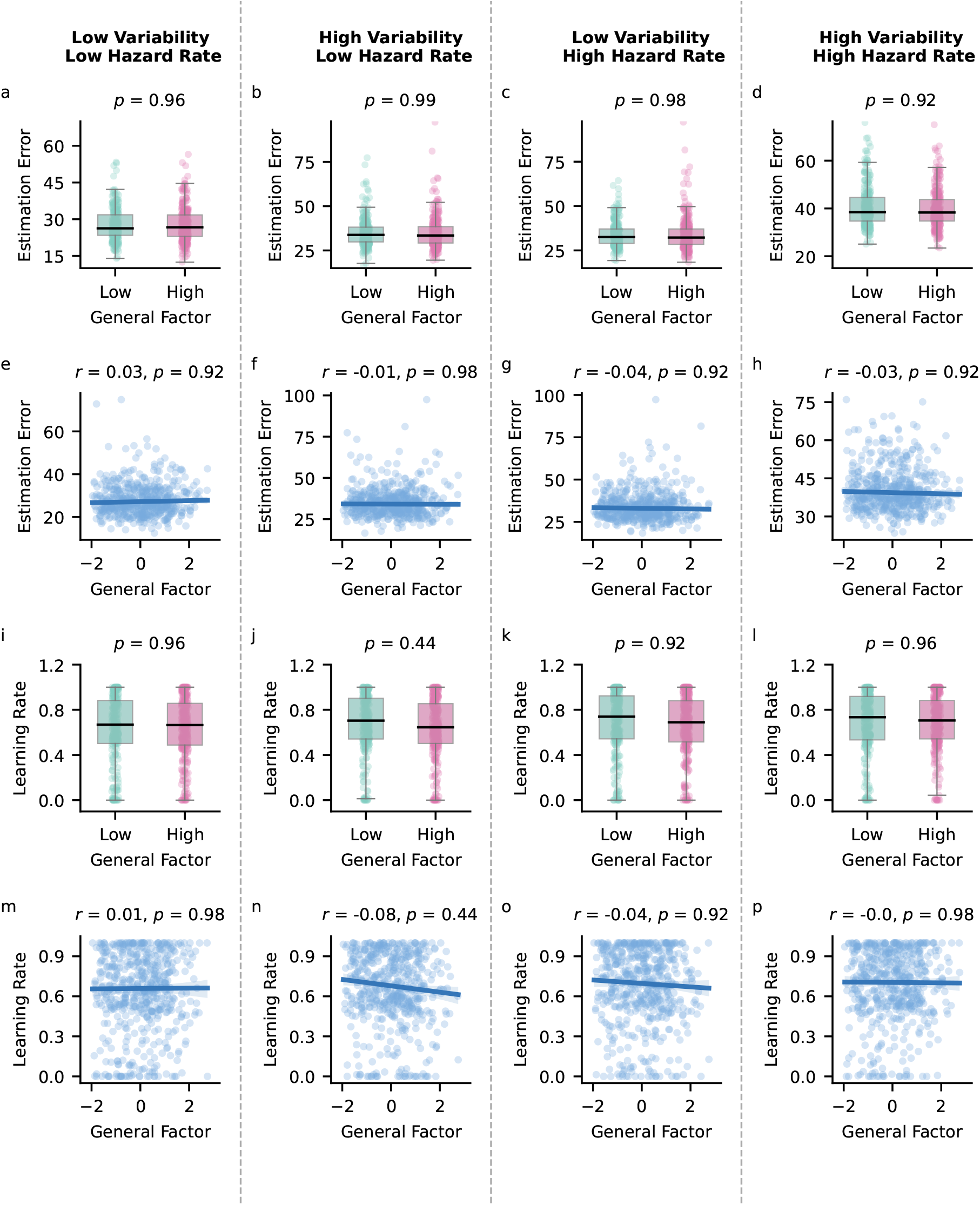
Effect of internalizing on estimation errors and empirical learning rates across variability and hazard-rate conditions in the predator task. **a-d**| Participants were categorized into low- and high-internalizing groups based on their general-factor scores. We did not find any significant differences in estimation errors between groups in any condition: low variability, low hazard rate (**a**); high variability, low hazard rate (**b**); low variability, high hazard rate (**c**); or high variability, high hazard rate (**d**). **e-h**| Regression analysis of estimation errors against internalizing did not reveal any significant associations across conditions. **i-l**| Similarly, empirical learning rates were not found to differ significantly between internalizing groups across all variability and hazard-rate conditions. **m-p**| Regression analyses also did not show any significant associations between internalizing and learning rates across conditions. All *p*-values were corrected for multiple comparisons using false discovery rate correction.

**Figure S6.**
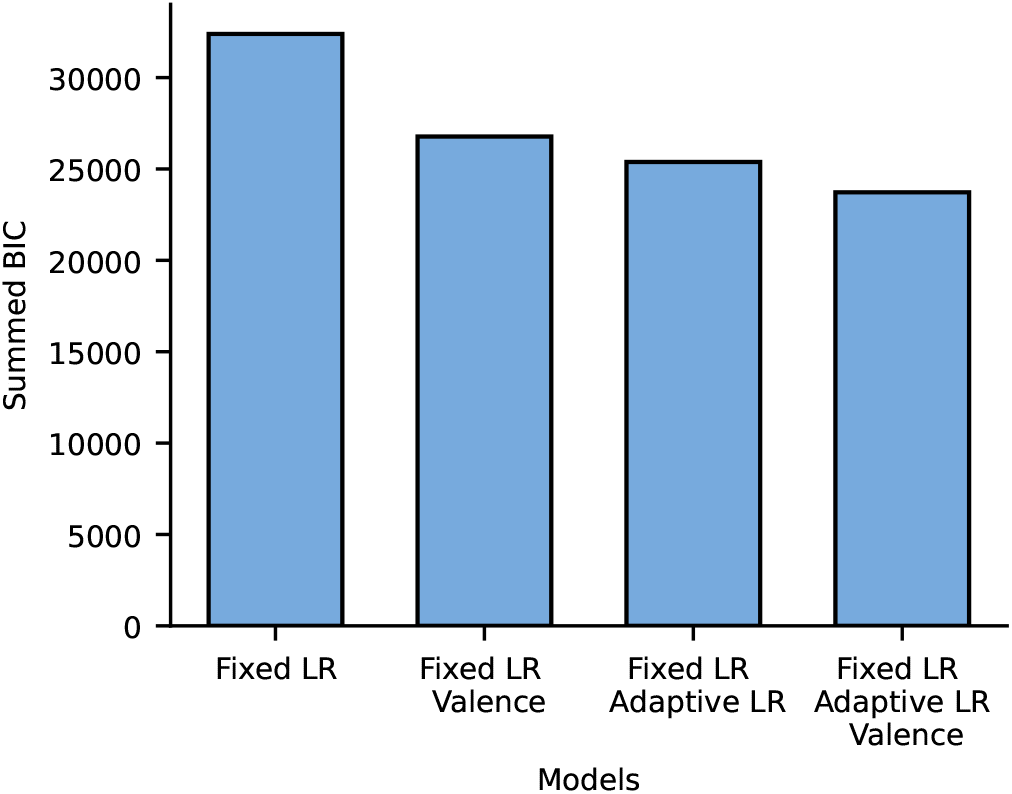
Model comparison for the linear regression using the reduced Bayesian model. We evaluated multiple regression models, including those with a single fixed learning rate, a fixed learning rate with valence, a fixed and adaptive learning rate, and a combination of all three. All models included parameters accounting for hazard-rate level, stochasticity, and their interaction. The best-fitting model was the full model with a fixed learning rate, an adaptive learning rate, and valence, which achieved the lowest summed Bayesian information criterion (BIC) score.

#### Model Comparison

We compared different regression models for the predator task by evaluating their summed Bayesian Information Criterion (BIC) scores (Fig. S6). The tested models included: (1) a model with only a fixed learning-rate parameter, (2) a model with a fixed learning rate and valence parameter, (3) a model incorporating both fixed and adaptive learning rates, and (4) a model with fixed learning rate, adaptive learning rate, and valence. All models accounted for hazard rate and stochasticity levels. The best-fitting model, which included a fixed learning rate, adaptive learning rate, and valence, achieved the lowest summed BIC score of 23722.34.

**Figure S7.**
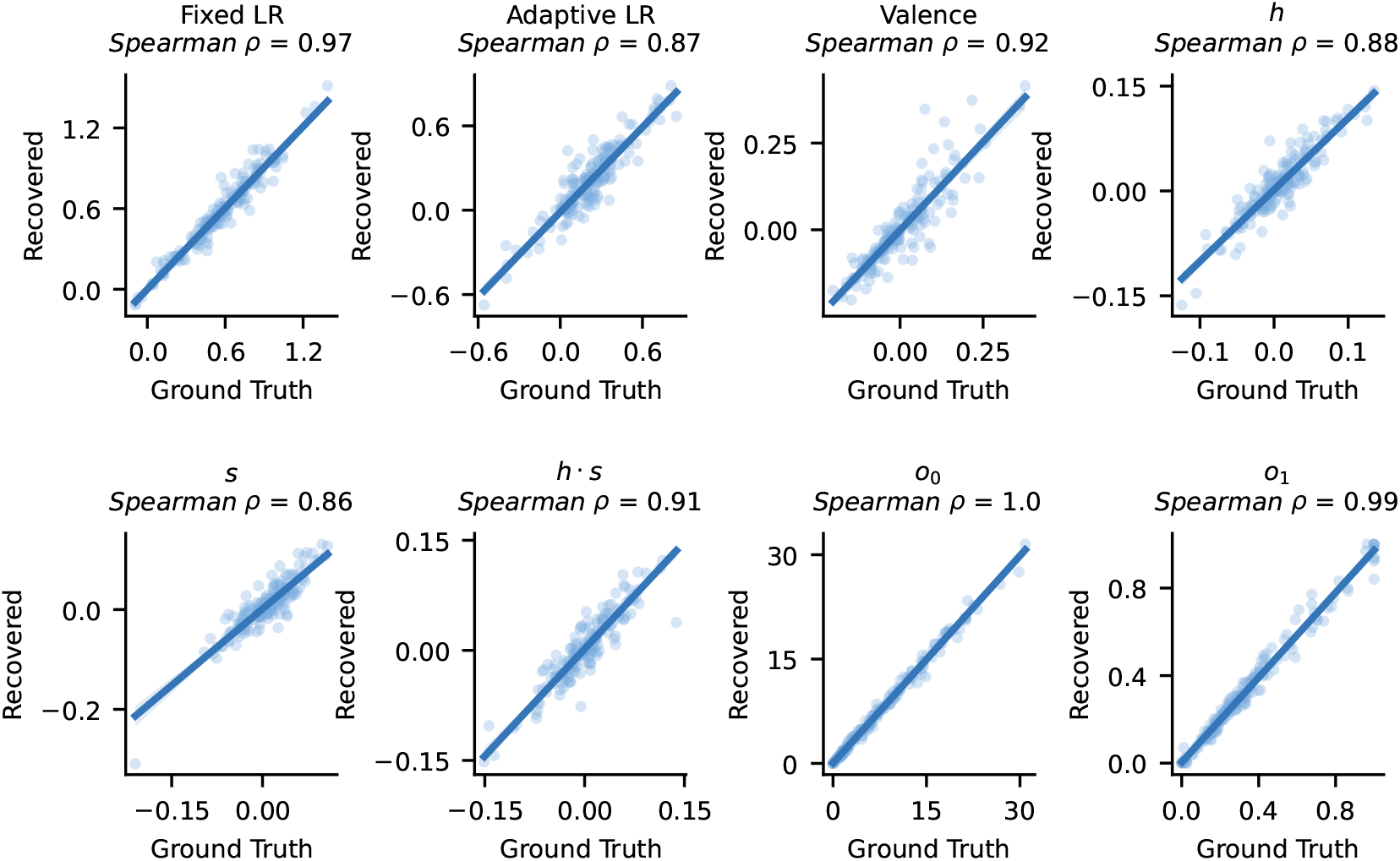
Parameter recovery analysis for the winning regression model. To evaluate parameter identifiability, we simulated five datasets using subject-specific parameter values from the winning model. The model was then applied to each dataset, and Spearman’s rank correlation was computed between the ground truth and recovered parameter values. The figure presents results from one example dataset, where each panel corresponds to a model parameter (x-axis: ground truth values, y-axis: recovered values). Across all five simulated datasets, the average Spearman correlation was *ρ* = 0.93, highlighting excellent parameter recoverability.

#### Parameter Recovery

We conducted a parameter recovery analysis to evaluate the identifiability of parameters in the winning regression model. Using subject-specific regression parameters from the winning model, we simulated trial-by-trial updates across five datasets. The model was then applied to each simulated dataset, and Spearman’s rank correlation was computed between the original parameter values (ground truth) and the estimated values (recovered parameters) for each dataset. The parameters exhibited excellent identifiability, with an average correlation across datasets and components of Spearman’s *ρ* = 0.93. An example of parameter recovery from one of the simulated datasets is shown in Fig. S7.

**Figure S8.**
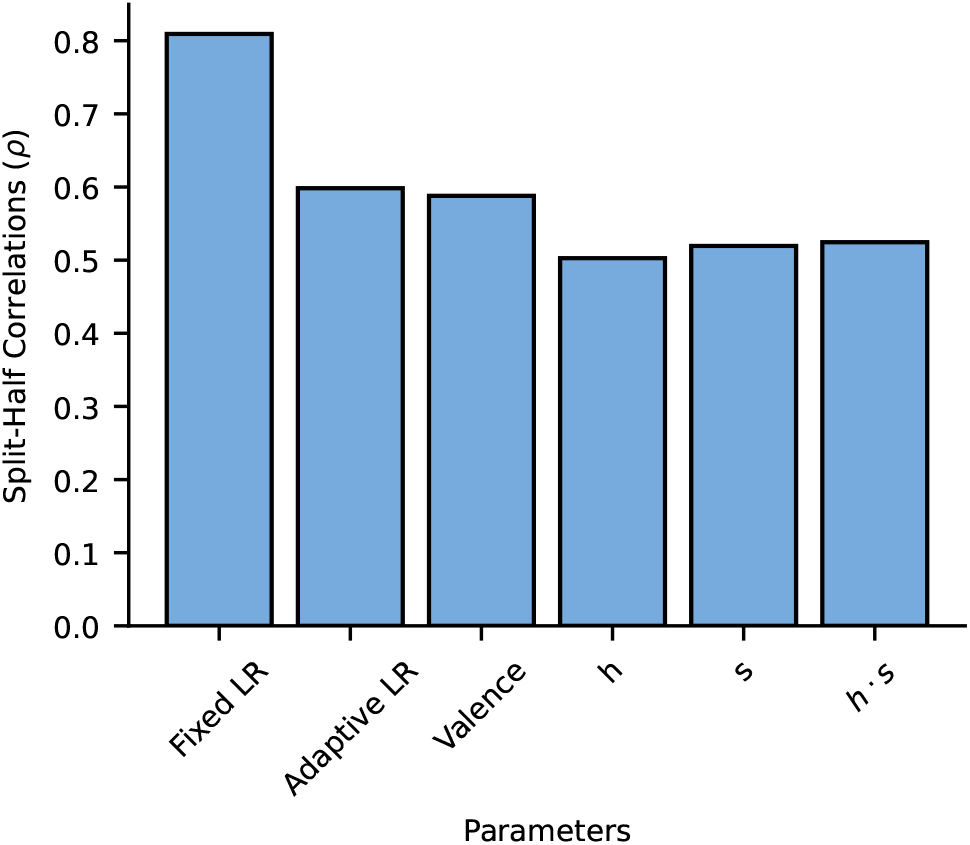
Split-half reliability of model parameters in the linear regression model. We assessed split-half reliability by dividing the data into odd and even trials, fitting the linear regression model to each subset, and calculating Spearman’s rank correlation *ρ* between the resulting model parameters. This yielded moderate-to-good correlations, indicating reliable model parameters, in particular for the fixed learning rate.

#### Split-Half Reliability

We used a linear regression model combined with the reduced Bayesian model to quantify the extent to which participants relied on fixed and adaptive learning rates. This model included parameters representing trial success (Valence) and parameters for variability and hazard-rate conditions. To evaluate the reliability of these model parameters, we conducted a split-half reliability analysis. The dataset was divided into odd and even trials, the regression model was applied separately to each subset, and Spearman’s rank correlation coefficients were calculated between the resulting parameter estimates (Fig. S8). We found moderate-to-good correlations for all model parameters (Fixed LR: Spearman *ρ* = 0.81, Adaptive LR: Spearman *ρ* = 0.6, Valence: Spearman *ρ* = 0.59, *h*: Spearman *ρ* = 0.5, *s*: Spearman *ρ* = 0.52, *s* · *h*: Spearman *ρ* = 0.52), indicating reliable parameter estimates.

#### Model Results

A robust linear regression of model parameters against factor scores did not reveal any significant association between factor scores and model parameters after applying FDR correction (Fig. S9, Table S2). The only significant effect that we observed was an association between gender and valence: females exhibited a higher learning rate following unsuccessful trials and a lower learning rate following successful trials compared to males (*r*(554.0) = -0.127, *p* = 0.022, *BF*_01_ = 0.066).

To investigate potential experiment-specific effects, we further examined the relationship between fixed and adaptive learning rates and general-factor scores across the four experiments (Fig. S10). Consistent with the initial analysis, we did not find significant associations between either the fixed or adaptive learning rates and internalizing psychopathology in any experiment.

As a final analysis, we examined the relation between questionnaire scores (including subscales) and fixed and adaptive learning rates extracted from the regression model, using FDR correction for multiple comparisons (Fig. S11). Fixed learning rates were not found to be significantly associated with any questionnaire or their associated sub-scales (Fig. S11 a-k). Similarly, adaptive learning rates were also not found to be significantly associated with any of the questionnaires (Fig. S11 l-v), except for the anxious-arousal subscale of the Mood and Anxiety Symptom Questionnaire (MASQ-AnxAr). Here, we observed a significant negative association, with higher MASQ-AnxAr scores linked to lower adaptive learning rates (*r*(554.0) = -0.14, *p* < 0.001). However, this finding should be interpreted with caution, as MASQ-AnxAr scores were positively skewed, with relatively few participants scoring high on this subscale (Fig. S11u).

**Table S2.** Estimates from the regression model predicting learning-related parameters from the factor scores derived from our factor analysis while controlling for age and gender. All *p*-values are corrected for multiple comparisons using false discovery rate correction. G = General-Factor Score (Internalizing), F1 = Anxiety-related factor, F2 = Depression-related factor.

| Target Variable | Predictor | Estimate |
| --- | --- | --- |
| Fixed LR<br>($\beta_1$) | G | 0.003 ($p = 0.959$, $BF_{01} = 20.833$) |
| | F1 | 0.037 ($p = 0.744$, $BF_{01} = 14.493$) |
| | F2 | -0.128 ($p = 0.051$, $BF_{01} = 0.254$) |
| | Age | -0.04 ($p = 0.744$, $BF_{01} = 13.699$) |
| | Gender | -0.041 ($p = 0.744$, $BF_{01} = 13.333$) |
| Adaptive LR<br>($\beta_2$) | G | -0.039 ($p = 0.744$, $BF_{01} = 13.333$) |
| | F1 | -0.101 ($p = 0.105$, $BF_{01} = 1.127$) |
| | F2 | 0.067 ($p = 0.47$, $BF_{01} = 5.682$) |
| | Age | 0.026 ($p = 0.783$, $BF_{01} = 17.241$) |
| | Gender | -0.049 ($p = 0.707$, $BF_{01} = 10.309$) |
| Valence<br>($\beta_3$) | G | 0.003 ($p = 0.959$, $BF_{01} = 20.833$) |
| | F1 | -0.026 ($p = 0.783$, $BF_{01} = 16.393$) |
| | F2 | -0.024 ($p = 0.783$, $BF_{01} = 16.949$) |
| | Age | -0.091 ($p = 0.105$, $BF_{01} = 1.074$) |
|  | Gender | **-0.127 ($p = 0.022$), $BF_{01} = 0.066$** |
| $h$<br>($\beta_4$) | G | 0.017 ($p = 0.744$, $BF_{01} = 14.493$) |
| | F1 | 0.005 ($p = 0.889$, $BF_{01} = 20$) |
| | F2 | -0.021 ($p = 0.744$, $BF_{01} = 12.048$) |
| | Age | -0.007 ($p = 0.874$, $BF_{01} = 19.608$) |
| | Gender | 0.037 ($p = 0.341$, $BF_{01} = 3.906$) |
| $s$<br>($\beta_5$) | G | -0.014 ($p = 0.783$, $BF_{01} = 17.857$) |
| | F1 | 0.001 ($p = 0.959$, $BF_{01} = 20.833$) |
| | F2 | -0.034 ($p = 0.629$, $BF_{01} = 7.812$) |
| | Age | -0.011 ($p = 0.815$, $BF_{01} = 18.519$) |
| | Gender | -0.012 ($p = 0.815$, $BF_{01} = 18.519$) |
| $h \cdot s$<br>($\beta_6$) | G | 0.029 ($p = 0.707$, $BF_{01} = 10.101$) |
| | F1 | 0.022 ($p = 0.744$, $BF_{01} = 13.889$) |
| | F2 | 0.019 ($p = 0.749$, $BF_{01} = 15.152$) |
| | Age | 0.018 ($p = 0.783$, $BF_{01} = 16.129$) |
| | Gender | 0.014 ($p = 0.783$, $BF_{01} = 17.544$) |

**Figure S9.**
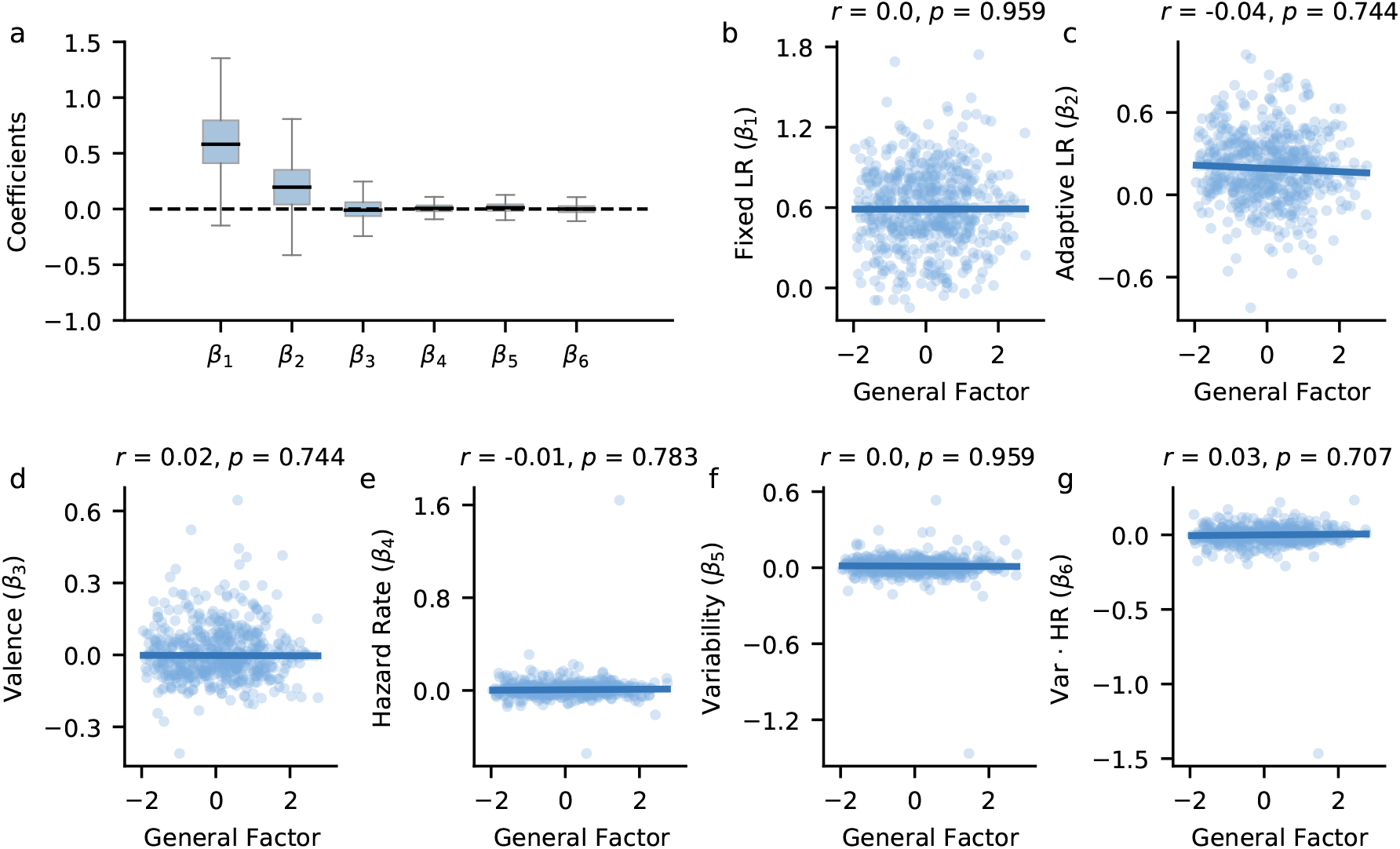
Model parameters from the linear regression using the reduced Bayesian model did not reveal any significant effects of internalizing on learning rates in the predator task. **a**| Boxplots of model parameters across participants indicate the use of both fixed and adaptive learning rates to facilitate learning. **b-g**| We did not find any significant associations between internalizing and the fixed learning rate (**b**), adaptive learning rate (**c**), valence (**d**), hazard-rate levels (**e**), variability levels (**f**), or the interaction between variability and hazard-rate levels (**g**). All *p*-values were FDR-corrected.

**Figure S10.**
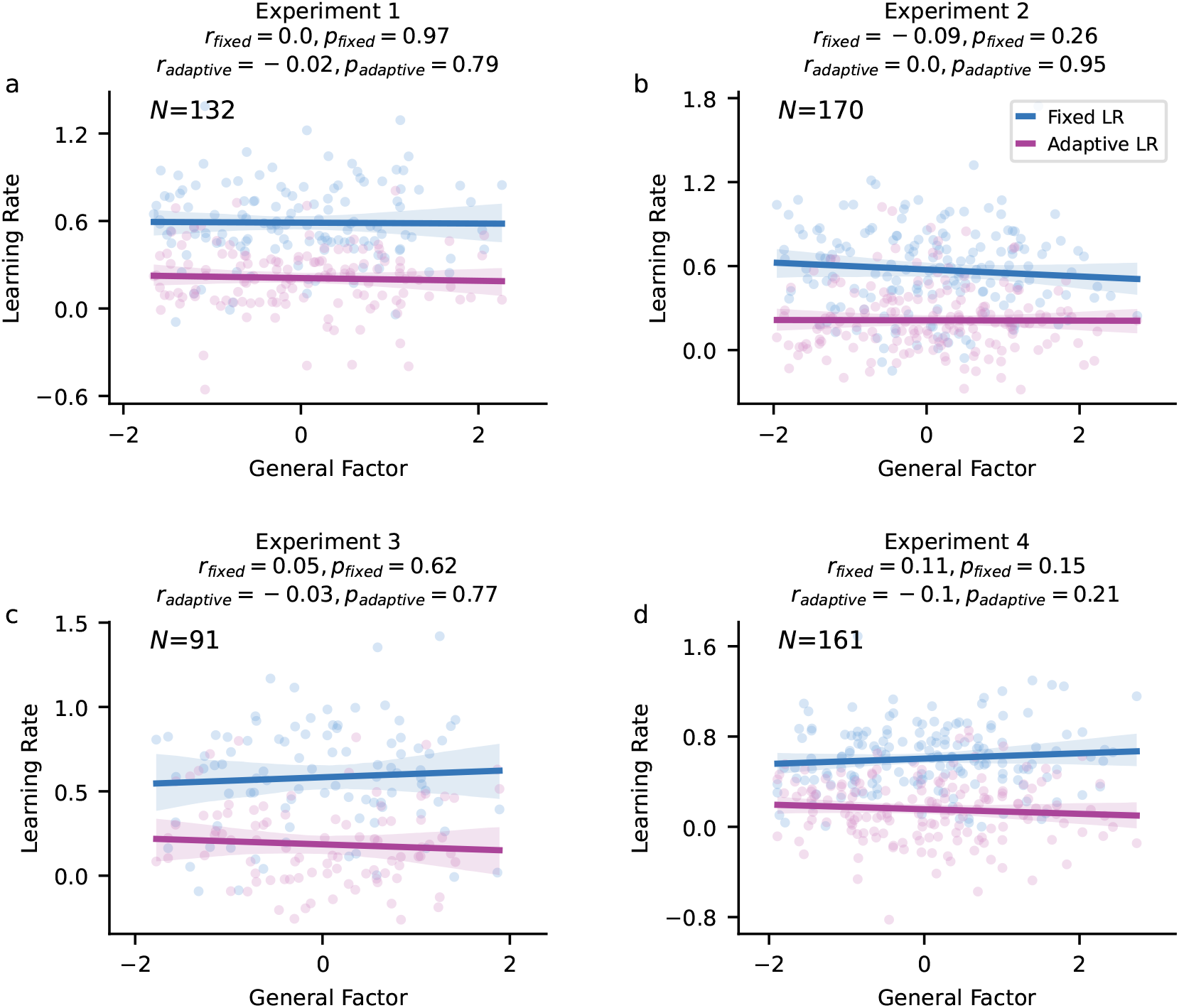
Regression of model parameters against general-factor scores across experiments. We did not find any significant associations between internalizing psychopathology and fixed or adaptive learning rates (LR) across all four experiments. **a**| In experiment 1, participants completed a predator-task version with different conditions. Neither fixed nor adaptive learning rates were found to be significantly associated with internalizing. **b**| In experiment 2, participants completed the predator task along with the probabilistic reversal learning task. Fixed and adaptive learning rates did not reveal any significant associations with internalizing in the predator task. **c**| In experiment 3, participants completed the predator task and a probabilistic reversal learning task with reward magnitudes. We did not find any significant associations between internalizing and fixed or adaptive LRs. **d**| In experiment 4, participants completed the predator task along with a probabilistic reversal learning task incorporating both reward and loss conditions. Fixed and adaptive LRs in the predator task were not found to be significantly associated with internalizing.

**Figure S11.**
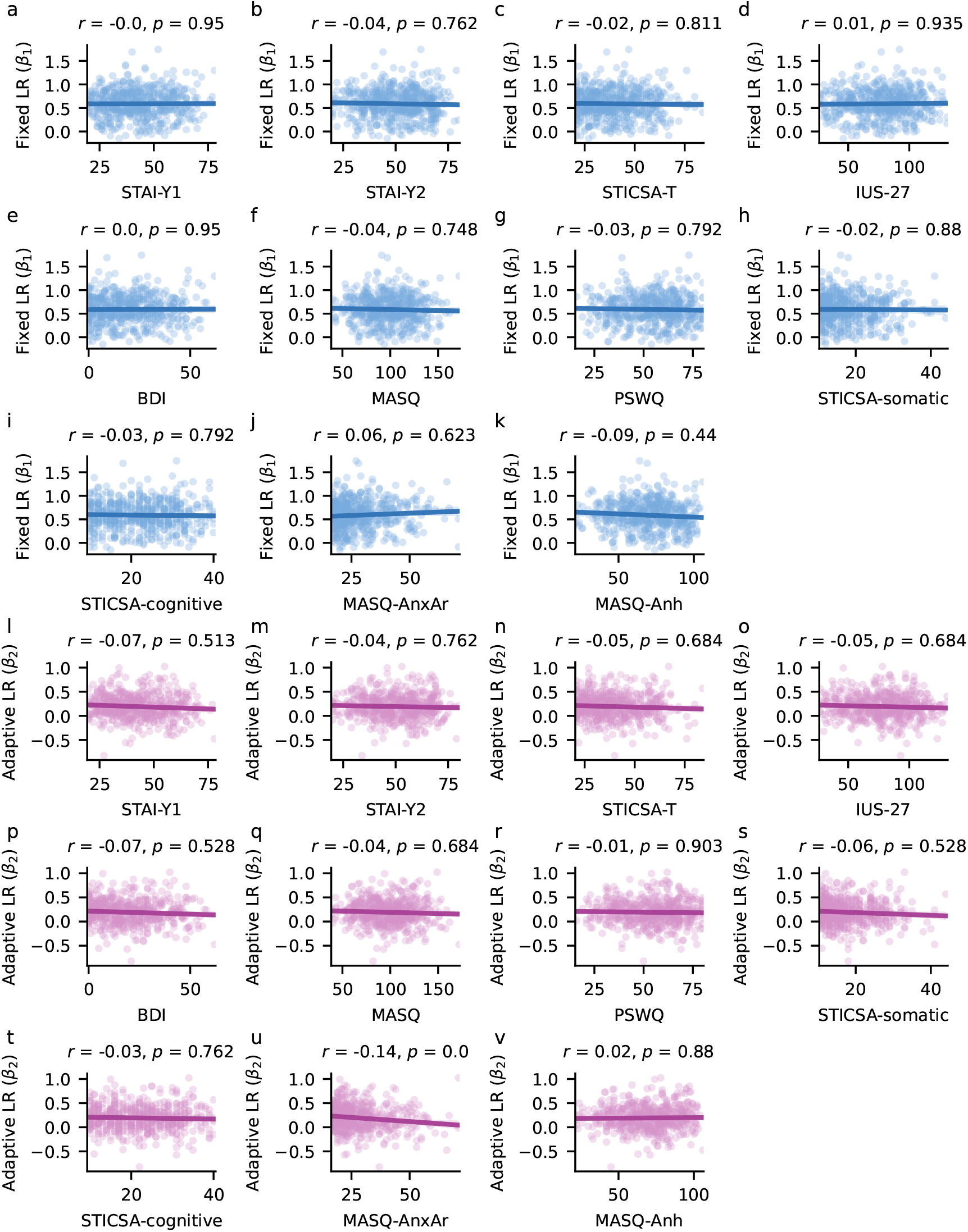
Regression of model parameters against questionnaire scores and associated subscales, controlling for age and gender. **a-k**| Association between fixed learning rate (LR) and questionnaire scores. No significant associations were observed between fixed LR and any questionnaire or subscale. **l-v**| Association between adaptive LR and questionnaire scores. The only significant association was a decrease in adaptive LR with increasing scores on the MASQ-AnxAr sub-scale. All *p*-values are corrected based on the false discovery rate. STAI-Y1: Spielberger State-Trait Anxiety Inventory State Scale; STAI-Y2: Spielberger State-Trait Anxiety Inventory Trait Scale; STICSA-T: State-Trait Inventory for Cognitive and Somatic Anxiety - Trait scale, IUS-27: Intolerance of Uncertainty Scale; BDI: Beck’s Depression Inventory; PSWQ: Penn State Worry Questionnaire; MASQ: Mood and Anxiety Symptom Questionnaire; STICSA-cognitive: State-Trait Inventory for Cognitive and Somatic Anxiety - Trait cognitive subscale; STICSA-somatic: State-Trait Inventory for Cognitive and Somatic Anxiety - Trait somatic subscale; MASQ-Anh: Mood and Anxiety Symptom Questionnaire - anhedonia subscale; MASQ-AnxAr: Mood and Anxiety Symptom Questionnaire - anxious arousal subscale

**Figure S12.**
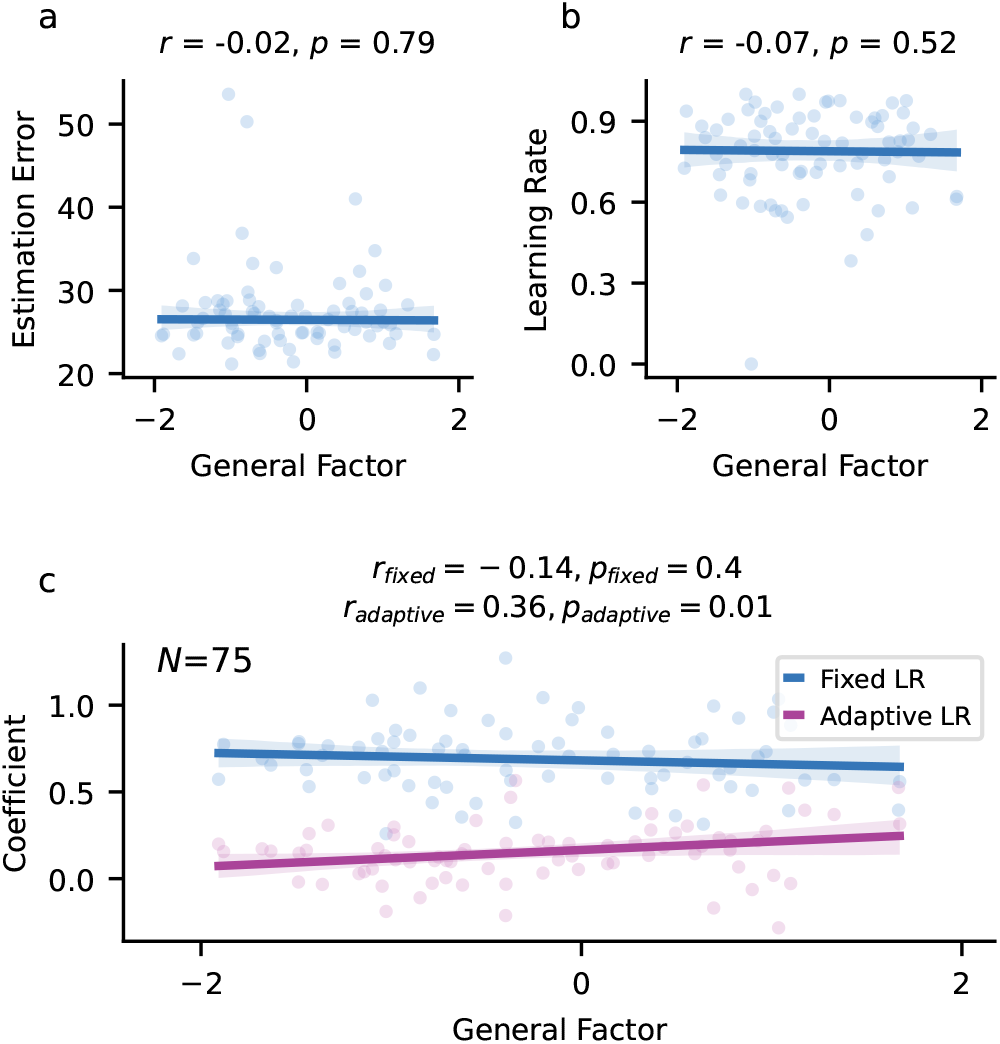
Results from the predator-task version with minimal training phase. Descriptive and model-based behavioral results were not found to be significantly associated with internalizing (general-factor scores). **a**| Estimation errors did not show significant associations with internalizing. **b**| Single-trial learning rates indicate participants exhibited higher overall learning rates, but these were not found to be significantly associated with internalizing. **c**| Regression of fixed learning rate with internalizing did not reveal any significant associations with internalizing. However, we observed a significant positive relationship between internalizing and adaptive learning rates, suggesting that individuals with higher internalizing symptoms exhibited more adaptive learning behavior. All *p*-values are corrected for multiple comparisons based on the false discovery rate.

#### Minimal-Training Version

To investigate the potential effects of task training on learning, *N* = 75.0 participants completed the predator task with minimal training under a low variability and hazard-rate condition in experiment 5. This design aimed to examine whether task training could mask learning impairments associated with internalizing. The analysis did not yield significant associations between internalizing and descriptive measures of learning (Fig. S12). Estimation errors (median = 26.22, IQR 24.72 to 28.3) and single-trial learning rates showed no significant association with internalizing, although participants exhibited higher overall single-trial learning rates (median = 0.81, IQR 0.7 to 0.9). Similarly, the test did not reveal a significant associations between the fixed learning rate and internalizing (*r*(75.0) = -0.14, *p* = 0.4, *BF*_01_ = 4.132). However, we observed a significant positive association between the adaptive learning rate and internalizing (*r*(75.0) = 0.36, *p* = 0.01, *BF*_01_ = 0.036), suggesting that individuals with higher internalizing symptoms adjusted their learning more flexibly in the minimal training version.

**Figure S13.**
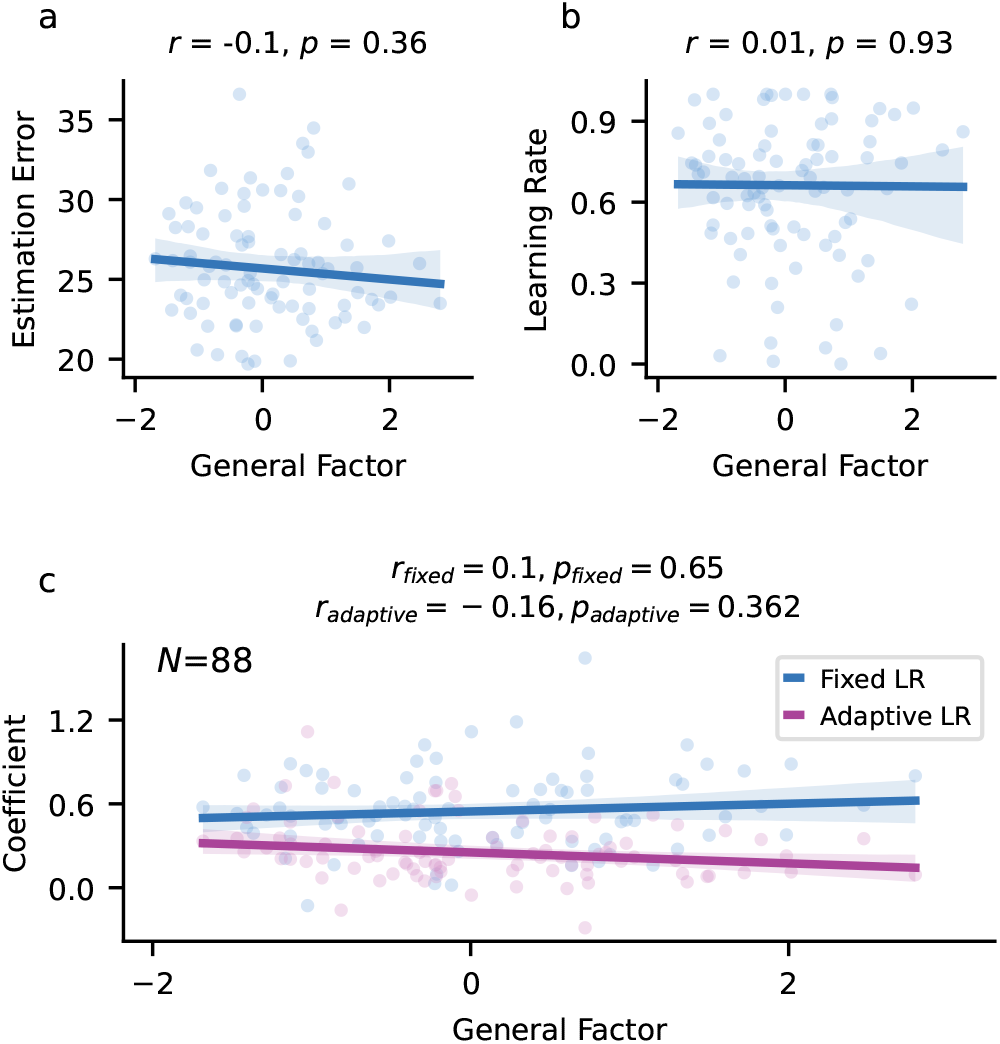
Results from the predator-task version with full training and a single variability and hazardrate condition. Descriptive and model-based parameters were not found to be significantly associated with internalizing (general-factor scores) in the predator-task version featuring only one variability and hazard rate condition (low variability, low hazard rate). **a**| Estimation errors in this task version were not found to be significantly associated with internalizing. **b**| Single-trial learning rates did not reveal significant associations between learning and internalizing. **c**| Regression of model parameters estimated using the reduced Bayesian model with internalizing did not reveal any significant associations with internalizing. All *p*-values of this regression are corrected using the false discovery rate.

#### Control Version

In experiment 6, *N* = 88.0 participants completed the predator task with full training under a low variability and hazard-rate condition. This design provided a high trial count for evaluating learning within a single condition and enabled comparisons with the minimal-training version of the task. Consistent with previous findings, we did not find significant associations between internalizing and either descriptive or model-based learning measures (Fig. S13).

#### In-Person Study

To assess the efficacy of screams as an aversive stimulus compared to electric shocks, experiments 7 and 8 incorporated versions of the predator task in which some blocks featured screams while others featured shocks. In experiment 7, we first observed a significant increase in skin conductance response (SCR) following outcome onset for failed trials compared to successful trials (horizontal gray line indicating significant SCR difference between the pink and green curves; Fig. S14a, *p* = 0.001). Next, we compared SCR in failed trials across shock and scream blocks. Both screams (green curve, *p <* 0.001) and shocks (pink curve, *p <* 0.001, Fig. S14b) elicited a significant SCR increase, confirming that screams function as effective aversive stimuli. Additionally, shocks produced a significantly greater SCR increase compared to screams (horizontal gray line, *p* = 0.023).

We replicated these patterns of results in experiment 8, which employed a different version of the predator task but retained the same stimulus structure, with separate blocks for screams and shocks. Again, SCR significantly increased after outcome onset for failed trials compared to successful trials (horizontal gray line, Fig. S14c, *p* = 0.001). Further analysis revealed that both screams (green curve, *p <* 0.001) and shocks (pink curve, *p <* 0.001, Fig. S14d) elicited significant SCR increases in failed trials, consistent with experiment 7. However, unlike in Experiment 7, we did not observe a significant difference in SCR between shocks and screams in this experiment (*p* = 0.44). These results validate the use of screams as an effective aversive stimulus, alleviating potential concerns that they might be insufficiently aversive to reveal learning impairments in internalizing individuals.

**Figure S14.**
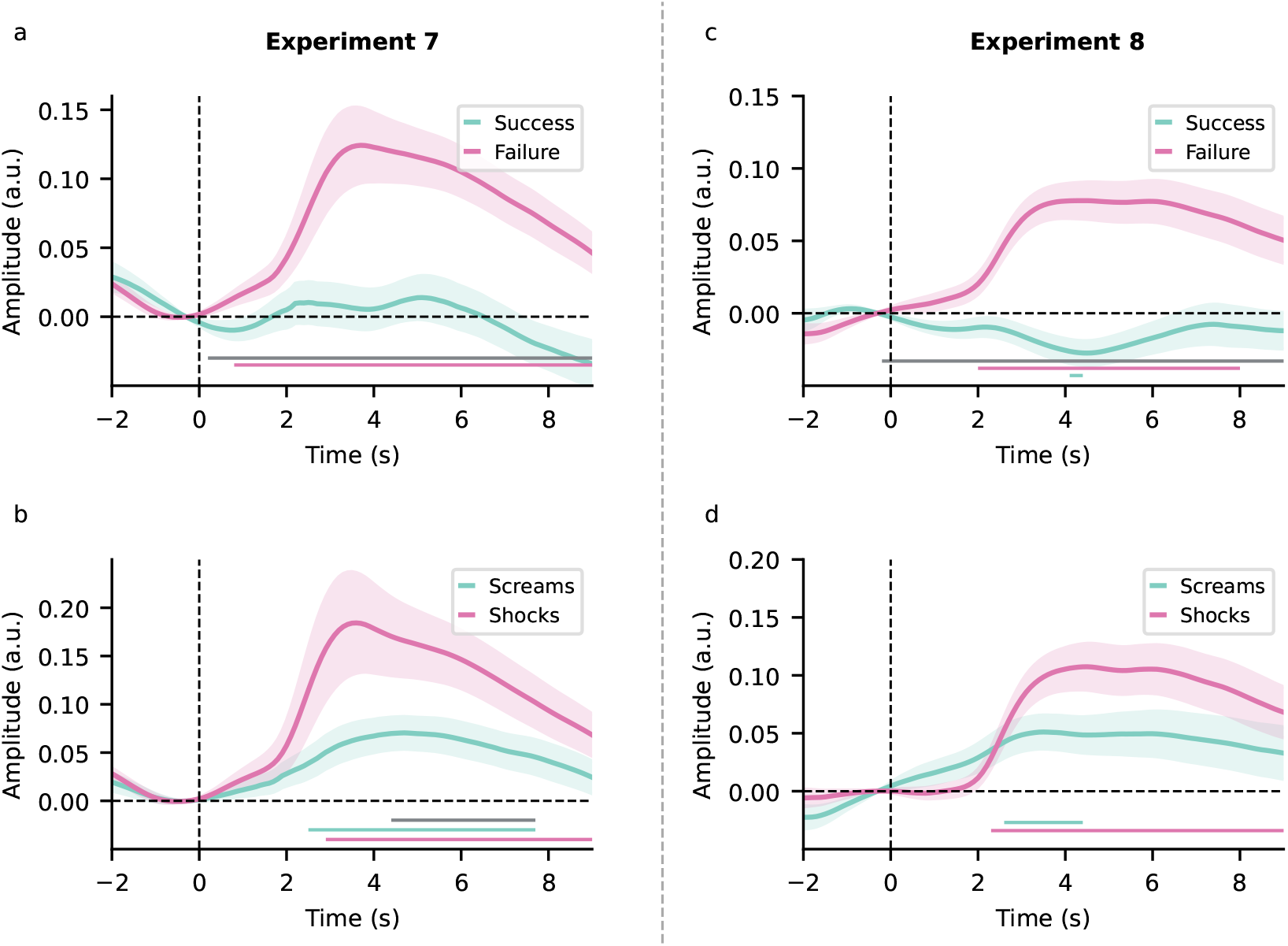
Results from the in-person studies with the predator task. **a**| In experiment 7, skin conductance response (SCR) significantly increased following outcome onset for failed trials (pink curve), whereas we did not observe a significant increase for successful trials (green curve). **b**| Further partitioning of failed trials into shock and scream phases revealed a significant SCR increase for both electric shocks (pink curve) and screams (green curve) after outcome onset. Additionally, SCR was significantly higher for shocks compared to screams in experiment 7 (horizontal gray line). **c**| Replicating the pattern observed in experiment 7, experiment 8 also showed a significant SCR increase for failed trials after outcome onset, while successful trials did not exhibit a significant increase. **d**| Similarly, in experiment 8, failed trials showed a significant SCR increase for both electric shocks (pink curve) and screams (green curve) after outcome onset. However, unlike in experiment 7, SCR for shocks was not significantly different from screams based on permutation testing. Horizontal lines in each plot indicate significant SCR regions identified through permutation testing: the pink line corresponds to the pink curve, the green line corresponds to the green curve, and the gray line represents the region of significant differences between conditions.

### Binary Reversal Learning Task

#### Task Version Without Reward Magnitudes

In experiment 2, *N* = 179 participants (demographic details in Table S3) completed a binary probabilistic reversal learning task. This task version did not include outcome magnitudes; participants were required to choose a fractal based on each fractal’s inferred underlying reward probabilities.

**Table S3.**
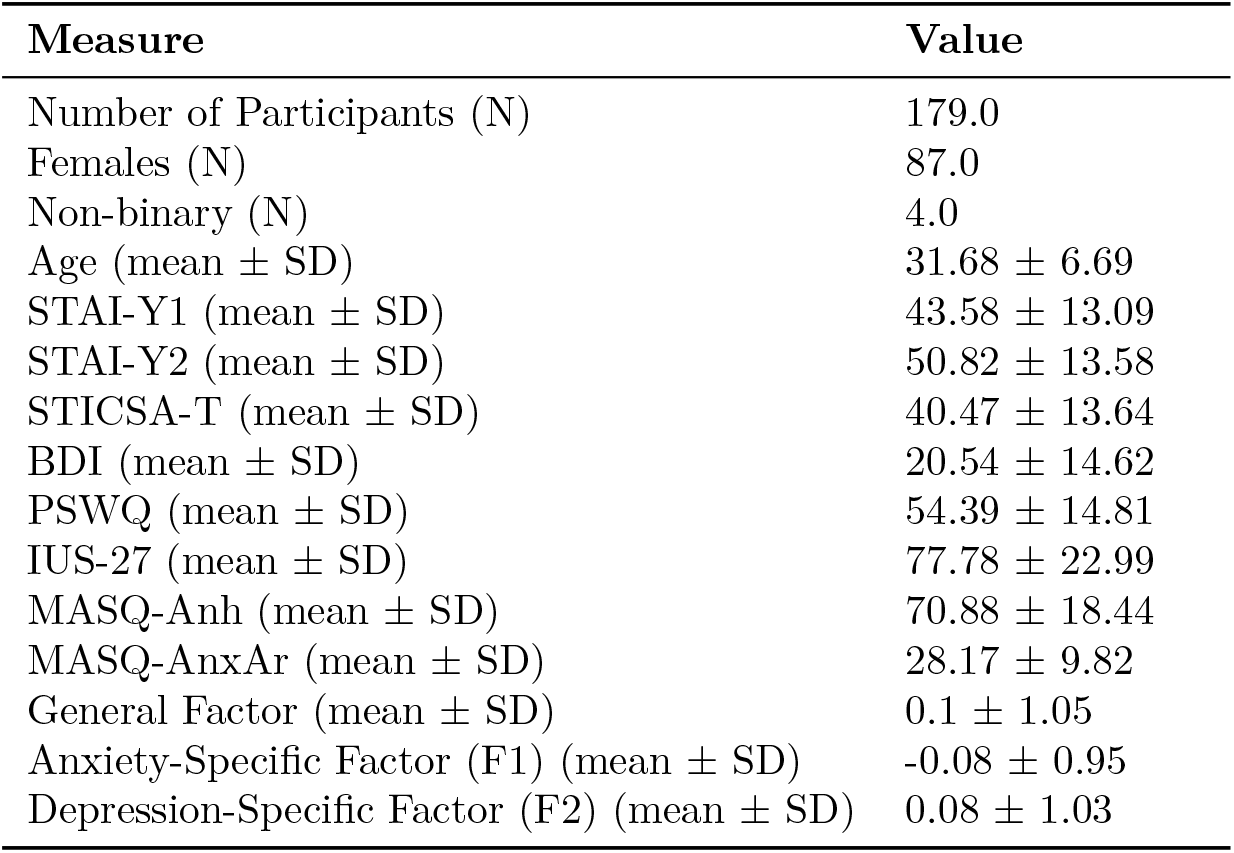
Basic demographic details of participants who completed the binary reversal learning task without reward magnitudes. STAI-Y1: Spielberger State-Trait Anxiety Inventory - state scale; STAI-Y2: Spielberger State-Trait Anxiety Inventory - trait scale; STICSA-T: State-Trait Inventory for Cognitive and Somatic Anxiety - trait scale, IUS-27: Intolerance of Uncertainty Scale; BDI: Beck’s Depression Inventory; PSWQ: Penn State Worry Questionnaire; MASQ: Mood and Anxiety Symptom Questionnaire; MASQ-Anh: Mood and Anxiety Symptom Questionnaire - anhedonia subscale; MASQ-AnxAr: Mood and Anxiety Symptom Questionnaire - anxious arousal subscale

#### Performance and Switch Rates

Task performance was quantified as the proportion of trials in which participants selected the highly rewarding fractal. Participants were categorized into low- and high-internalizing groups based on whether their scores fell below or above the mean general-factor score. We did not find significant differences in performance between these groups during either the stable phase (Welch’s *t*-test *t*_169.29_ = -1.72, *p* = 0.09, *BF*_01_ = 1.555) or the volatile phase (*t*_166.05_ = -1.6, *p*_*volatile*_ = 0.11, *BF*_01_ = 1.869; Fig. S15a). Similarly, regression analysis of performance against general-factor scores on a continuum did not reveal a significant association in either the stable (*r*(175) = 0.22, *p* = 0.06, *BF*_01_ = 0.381) or the volatile phase (*r*(175) = 0.17, *p* = 0.08, *BF*_01_ = 1.477; Fig. S15d). Although the *BF*_01_ values provided only anecdotal evidence for the null hypothesis, the observed trends, if anything, suggested a slight increase in performance with higher internalizing scores.

We next analyzed the percentage of switches following wins and losses, with switch percentage defined as the proportion of trials in which participants selected a different fractal than their previous choice. Participants had significantly higher switch rates after losses compared to wins. Specifically, in the stable phase, the median percentage of switches after wins was 12.04% compared to 47.69% after losses (paired *t*-test *t*_174_ = -20.18, *p <* 0.001, *BF*_01_ = 0). Similar results were found for the volatile phase, where the median percentage of switches after wins was 14.46% compared to 53.62% after losses (paired *t*-test *t*_174_ = -21.26, *p <* 0.001, *BF*_01_ = 0).

When participants were divided into low- and high-internalizing groups, no significant differences were observed in switch rates following wins in either the stable phase (Welch’s *t*-test *t*_169.41_ = 1.31, *p* = 0.19, *BF*_01_ = 2.755) or the volatile phase (*t*_163.01_ = 1.59, *p* = 0.11, *BF*_01_ = 1.898; Fig. S15b). Similarly, regression analyses of switch rates after wins, while controlling for age and gender, did not reveal a significant association in either the stable phase (*r*_*stable*_(175) = -0.16, *p*_*stable*_ = 0.08, *BF*_01_ = 1.536) or the volatile phase (*r*_*volatile*_(175) = -0.1, *p*_*volatile*_ = 0.27, *BF*_01_ = 4.975; Fig. S15e). Additionally, switch rates after losses were also not found to be significantly different between the low- and high-internalizing groups, in either the stable (*t*_173.0_ = -0.21, *p* = 0.84, *BF*_01_ = 5.988) or volatile phases (*t*_163.85_ = -0.04, *p* = 0.97, *BF*_01_ = 6.098; Fig. S15c). Corresponding regression analyses also did not yield any significant associations between internalizing and loss-switch rates in either the stable (*r*_*stable*_(175) = -0.06, *p*_*stable*_ = 0.54, *BF*_01_ = 9.009) or volatile phase (*r*_*volatile*_(175) = -0.03, *p*_*volatile*_ = 0.7, *BF*_01_ = 10.989; Fig. S15f). Overall, these findings suggest that internalizing is not associated with impaired adaptation in the reversal learning task.

**Figure S15.**
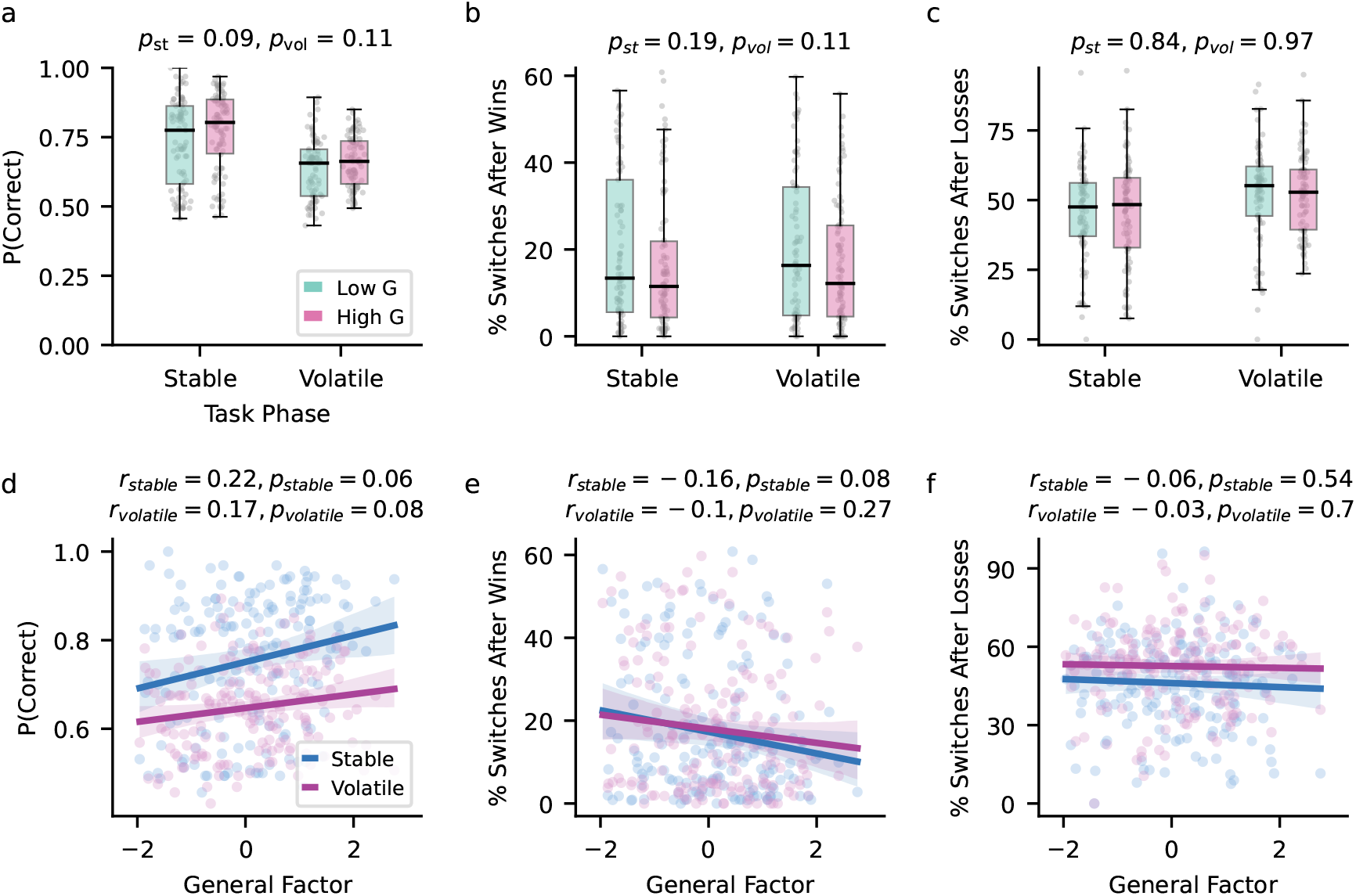
Performance and switch rates in the probabilistic reversal learning task. We analyzed performance and switch rates (proportion of trials in which participants chose a different fractal compared to their previous choice) by comparing participants with low and high general-factor scores. **a**| Performance, calculated as the proportion of trials in which participants chose the highly rewarding fractal, was not found to be significantly different between low- and high-internalizing groups in either the stable or volatile task phases. **b**| The percentage of switches following a win was not found to be significantly different between low- and high-internalizing groups. **c**| While participants switched more frequently after losses than after wins, switch rates after losses were not found to be significantly different between the groups. **d**| Performance was not found to be significantly associated with internalizing in either the stable or volatile phase. **e**,**f**| In neither stable nor volatile task phase did we find a significant association of internalizing with switch rates after wins (**e**) or switch rates after losses (**f**). All regression *p*-values are corrected based on the false discovery rate.

**Table S4.** Model comparison for the probabilistic reversal learning task. *α*: Learning-rate parameter, *β*: Inverse temperature, *δ*: Decay parameter. The subscript *ck* represents the choice-kernel parameter. Superscripts indicate parameter components: *sv*: Baseline component and a task phase-dependent component (stable vs. volatile). *gb*: Component encoding outcome valence (good vs. bad). *sv· gb*: Interaction between task phase and outcome valence. PSIS-LOO: Pareto-smoothed importance-sampling-based approximation of leave-one-out cross-validation. *SE*: standard error.

| Model Number | Parameters | # of Parameter Components | PSIS-LOO (SE) |
| --- | --- | --- | --- |
| Model 1 | $\alpha^{sv}, \beta^{sv}$ | 4.00 | 54159.47 (244.44 ) |
| Model 2 | $\alpha^{sv,gb}, \beta^{sv}$ | 5.00 | 52214.68 (248.21) |
| Model 3 | $\alpha^{sv,gb}, \beta^{sv,gb}$ | 5.00 | 52214.68 (248.21) |
| Model 4 | $\alpha^{sv,gb,sv \cdot gb}, \beta^{sv,gb}$ | 7.00 | 51769.51 (248.25) |
| Model 5 | $\alpha^{sv,gb,sv \cdot gb}, \beta^{sv,gb,sv \cdot gb}$ | 8.00 | 51766.30 (248.28) |
| **Model 6** | $\alpha^{sv,gb,sv \cdot gb}, \beta^{sv,gb,sv \cdot gb}, \alpha_{ck}, \beta_{ck}$ | 10.00 | **51489.81 (249.75)** |
| Model 7 | $\alpha^{sv,gb,sv \cdot gb}, \beta^{sv,gb,sv \cdot gb}, \delta^{sv}$ | 10.00 | 51799.15 (243.84) |
| Model 8 | $\alpha^{sv,gb,sv \cdot gb}, \beta^{sv,gb,sv \cdot gb}, \alpha_{ck}, \beta_{ck}, \delta^{sv}$ | 12.00 | 51512.25 (246.36) |

#### Model Comparison

We fitted multiple variations of the canonical Rescorla-Wagner model to the task data, with models differing primarily in how the learning rate and inverse temperature parameters were divided into components (see Model Space for details). Model 6 emerged as the best-fitting model, with a PSIS-LOO value of 51489.81 (standard error = 249.75; Table S4). For this model, all population-level and individual model parameters had a Gelman-Rubin statistic 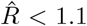,indicating consistency across chains.

**Figure S16.**
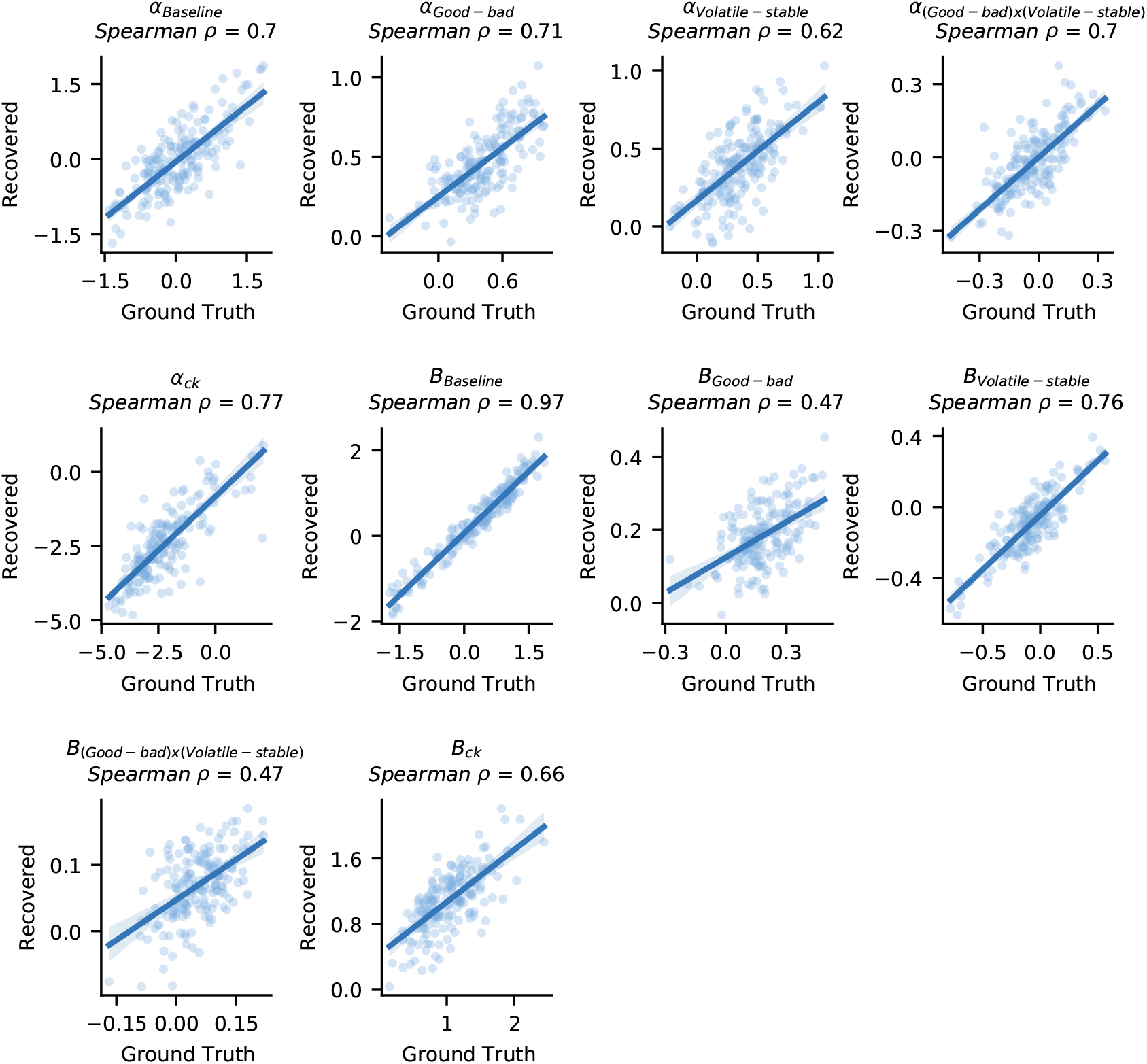
Parameter recovery analysis for the winning model (Model 6). To assess parameter identifiability, we simulated five datasets using subject-specific posterior means from the winning model. The model was then fitted to each dataset, and Spearman’s rank correlation was computed between the ground truth and recovered parameter values. The figure shows results for one example dataset, with each panel representing a model parameter (x-axis: ground truth parameter values, y-axis: recovered values). The average Spearman correlation between the ground truth and recovered parameters across all 5 simulated datasets was *ρ* = 0.68.

#### Parameter Recovery

We conducted a parameter recovery analysis to assess the identifiability of parameters in Model 6. We simulated choice data across five datasets using the subject-specific posterior means of all parameter components from the winning model. The model was then fitted to each simulated dataset, and Spearman’s rank correlation was computed between the original parameter values (ground truth) and the estimated values (recovered parameters) for each dataset. The parameters exhibited good identifiability, with an average correlation across datasets and components of Spearman’s *ρ* = 0.68. An example of parameter recovery from one of the simulated datasets is shown in Fig. S16.

To further assess parameter recoverability across a broader range of values, we generated an additional dataset for 500 subjects, in which the learning-rate components were sampled from a uniform distribution, *U* (− 2, 2). When the winning model was fitted to this dataset, recoverability remained strong, with a mean correlation between ground truth and recovered parameters of *ρ* = 0.76.

**Figure S17.**
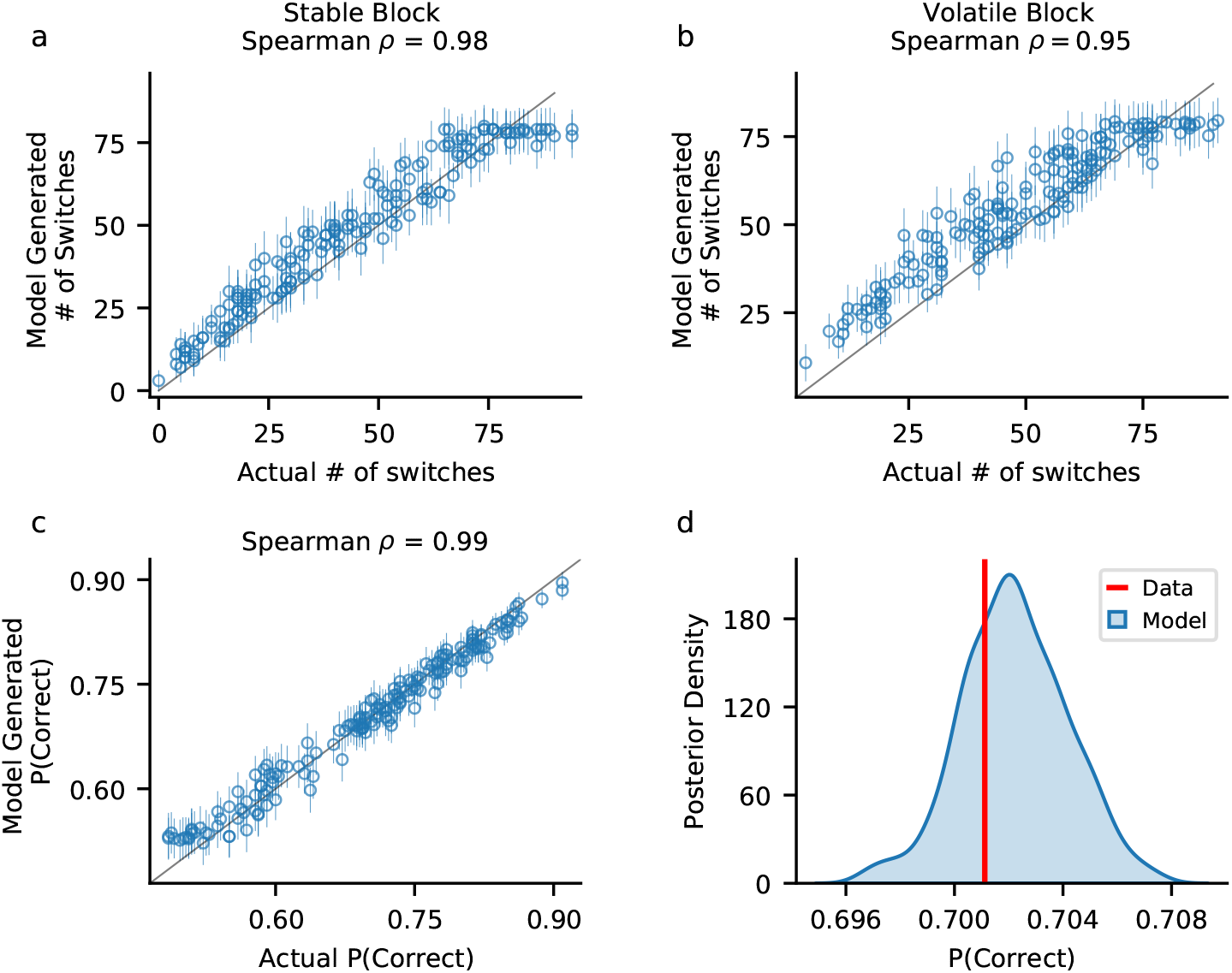
Posterior predictive checks for the winning model. To assess how well the winning model (Model 6) captured key qualitative features of choice behavior, we generated 500 simulated choice datasets per participant by drawing random samples from each participant’s joint posterior distribution (320 trials per dataset). **a-b**| Spearman correlations between the average number of switches across simulations and actual participant switches in the stable (**a**) and volatile (**b**) task phases. We observed strong correlations in both phases, indicating that the model successfully reproduces switching behavior. Circles represent the average number of switches, and error bars denote the standard deviation across simulations. **c**| Correlation between the proportion of trials in which the high-reward fractal was chosen in the simulated data and actual participants (P(Correct)). The strong correlation suggests that the model accurately captures choice accuracy. **d**| Distribution of the model’s predicted performance across trials and participants. The red line represents the actual average performance across participants. The close alignment between the predicted distribution and the actual value further supports the model’s ability to replicate key behavioral patterns.

#### Model Reproduction of Basic Features

To evaluate how well the model’s posterior predictions captured key qualitative aspects of participant choices, we analyzed both the number of switches and choice accuracy in model-generated choice data (Fig. S17). We simulated 500 choice datasets per participant by drawing 500 random samples from each participant’s joint posterior distribution, generating choices across 320 trials per dataset. We first examined switching behavior, defined as trials where participants selected a different fractal than in the previous trial. Comparing the number of switches in stable and volatile phases, we found strong Spearman correlations between the average number of switches across simulations and the actual number of switches made by participants, indicating that the model successfully reproduced switching behavior (*ρ* = 0.98 for the stable phase (Fig. S17a), and *ρ* = 0.95 for the volatile phase (Fig. S17b)). Next, we further analyzed the posterior predictions by comparing the proportion of trials where the high-reward fractal was chosen (P(Correct)) between the simulations and actual data (Zhang et al., 2020). Averaging across the 500 simulated datasets, we found a strong correlation with participant accuracy (*ρ* =0.99, Fig. S17c). Finally, to assess overall model performance, we computed the distribution of P(Correct) across the 500 simulations. This distribution was centered around the average actual P(Correct) of participants (Fig. S17d), further demonstrating that the model effectively captured key behavioral patterns.

**Figure S18.**
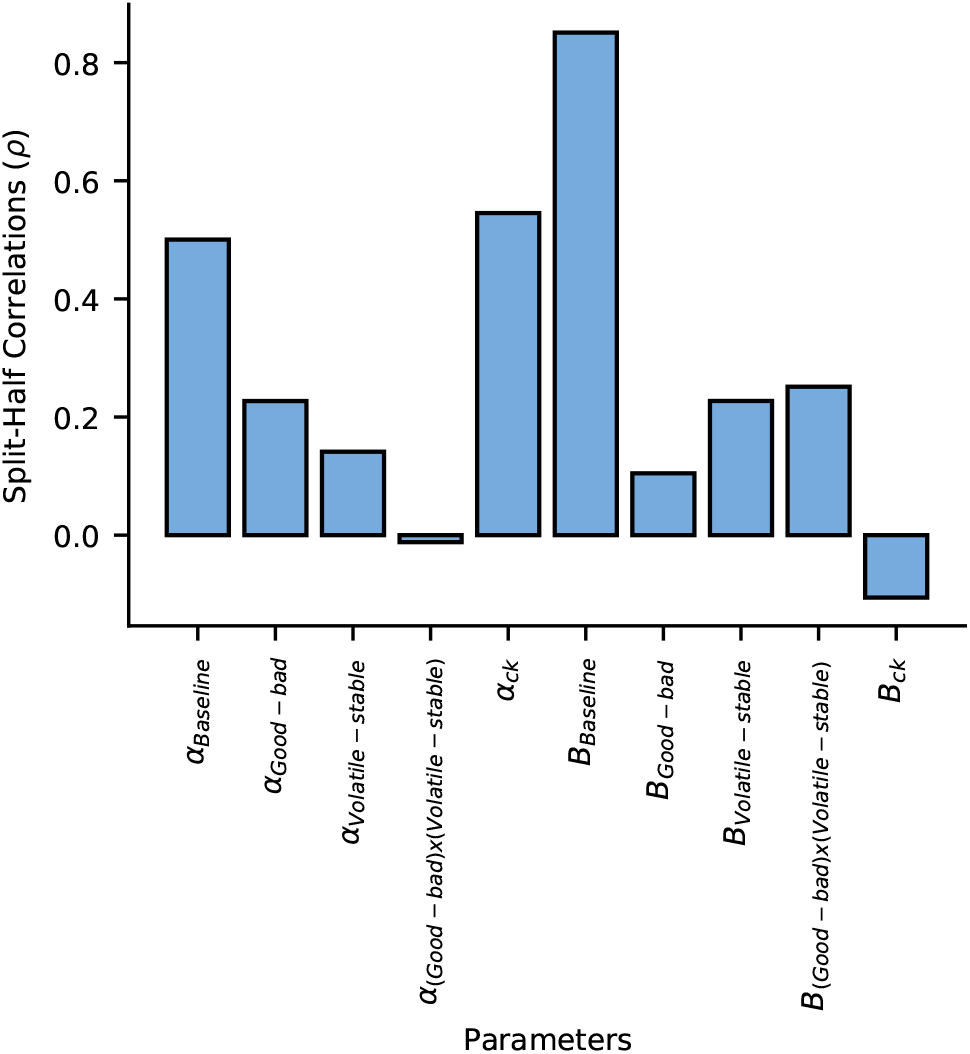
Split-half reliability of model parameters for the reversal learning task. To assess reliability, we split the data into two subsets: one containing the first block of each task phase and the other containing the second block. The winning hierarchical model (Model 6) was fit separately to each subset, and Spearman’s rank correlation *ρ* was computed for the resulting parameter estimates. The analysis revealed poor-to-moderate reliability for the learning-rate components.

#### Split-Half Reliability

The reversal learning task comprised two phases – volatile and stable – each consisting of two consecutive blocks. To assess the reliability of the model parameters, we split the dataset into two parts: one containing the first block of each phase and the other containing the second block. We then fit the winning model (Model 6) separately to both datasets and analyzed reliability by computing the Spearman rank correlation for each parameter (Fig. S18). The analysis revealed poor-to-moderate reliability for all learning-rate parameters (*α*_*Baseline*_ Spearman *ρ* = 0.5, *α*_*Good*−*bad*_ Spearman *ρ* = 0.23, *α*_*V olatile*−*stable*_ Spearman *ρ* = 0.14, *α*_(*Good* −*bad*)*x*(*V olatile*− *stable*)_ Spearman *ρ* = -0.01), with the choice-kernel learning rate showing moderate reliability (*α*_*ck*_ Spearman *ρ* = 0.55). For the inverse temperature parameter, only the baseline component exhibited good reliability (*B*_*Baseline*_ Spearman *ρ* = 0.85), while all other components showed poor reliability (*B*_*Good*−*bad*_ Spearman *ρ* = 0.1, *B*_*V olatile*−*stable*_ Spearman *ρ* = 0.23, *B*_(*Good*− *bad*)*x*(*V olatile*− *stable*)_ Spearman *ρ* = 0.25). These findings suggest generally low parameter reliability in the reversal learning task, aligning with estimates reported by Suddell et al. (2024).

**Figure S19.**
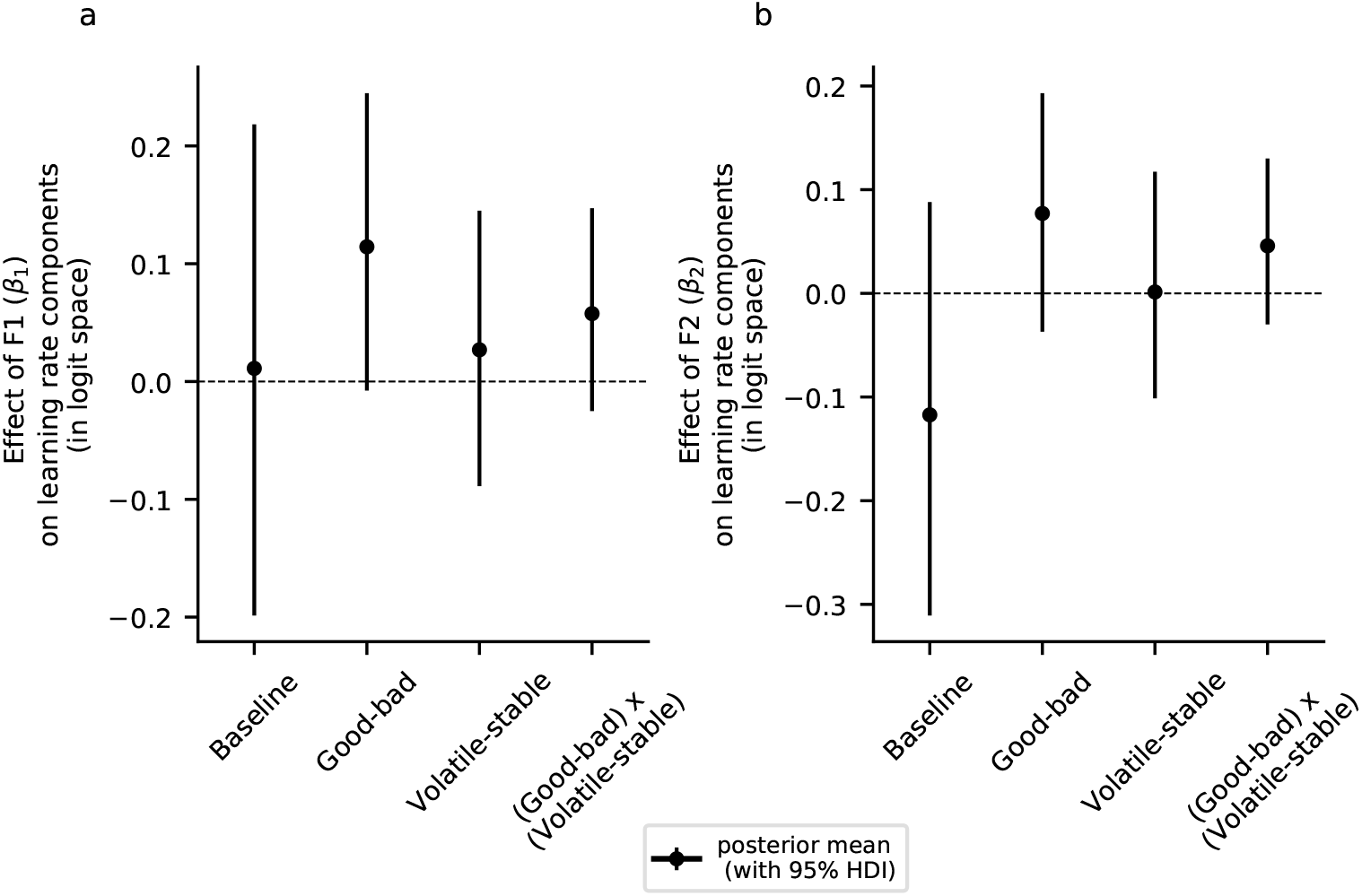
No significant effects of F1 and F2 on learning-rate components. **a**| The effect of the anxiety-related factor F1 on learning rates, represented by the *β*_1_ population-parameter posterior means and the corresponding 95% highest-density interval (HDI). We did not observe a significant effect of F1, as all HDIs included zero. **b**| The *β*_2_ population parameter, representing the depression-related factor F2, was also not found to have any significant effect on learning-rate components.

#### Model Results

We failed to find a significant effect of internalizing (general-factor scores) on any of the learning-rate components (Fig. 5). Similarly, analyses of the anxiety-related factor F1 and the depression-related factor F2 did not reveal significant associations with the learning-rate components (Fig. S19), and all 95% HDIs contained zero.

We next analyzed the components of the inverse temperature parameter and their associations with factor scores (Fig. S20). The parameter captures choice stochasticity, where higher values indicate more deterministic choices. At the group level, we observed significant effects of both outcome valence and task phase on the inverse temperature. Specifically, inverse temperature was higher after good outcomes more deterministic choices following a good outcome (*B*_*Good*− *bad*_ *µ*_0_ = 0.18, 95% HDI = [0.13, 0.23]). Task phase also significantly influenced inverse temperature, with lower values (more stochastic choices) in the volatile phase compared to the stable phase (*B*_*V olatile*− *stable*_ *µ*_0_ = -0.08, 95% HDI = [-0.14, -0.03]). Additionally, we found a significant interaction between outcome valence and task phase (*B*_(*Good*− *bad*)*x*(*V olatile* − *stable*)_; *µ*_0_=0.05, 95% HDI = [0.02, 0.09]; Fig. S20a).

**Figure S20.**
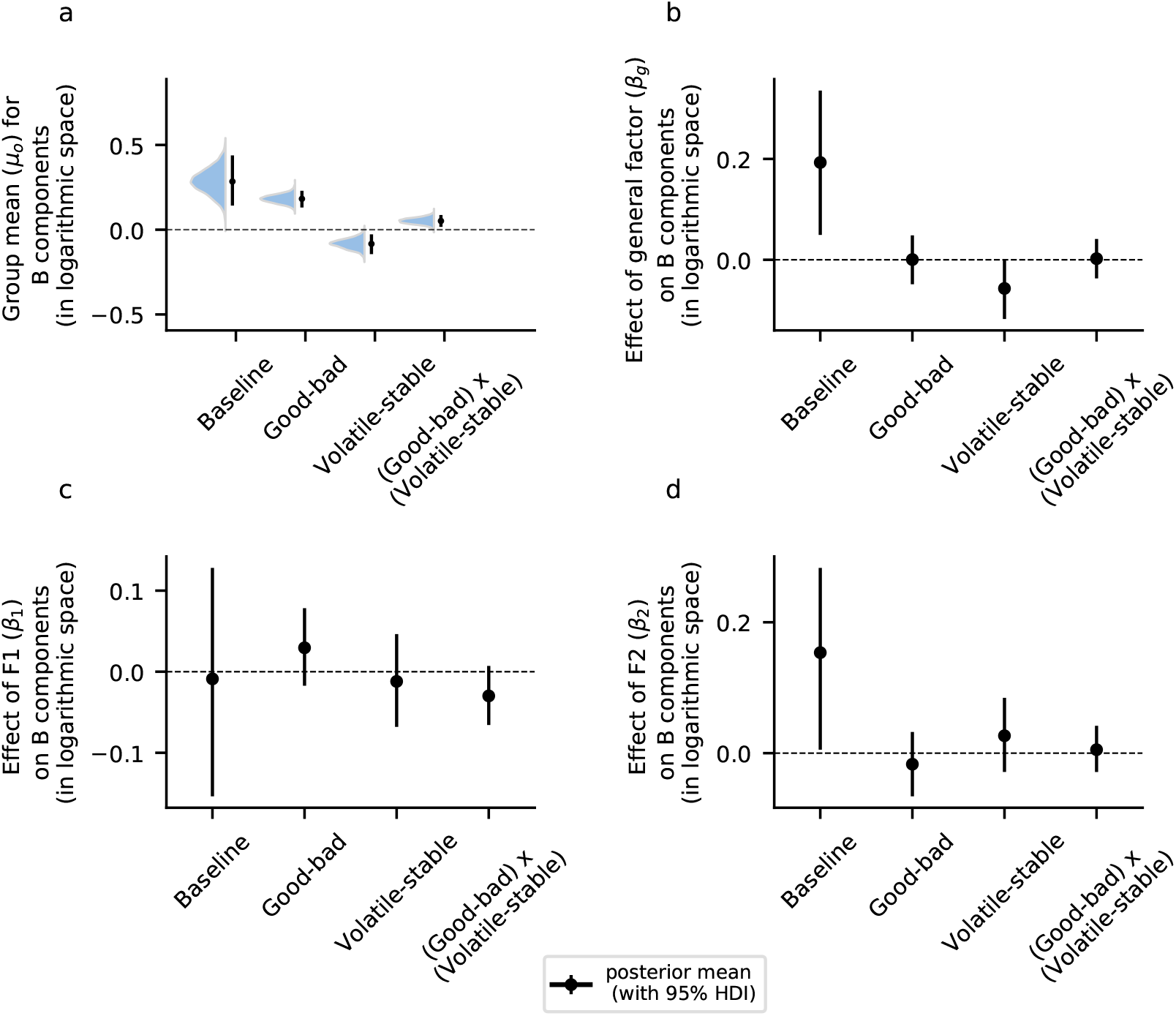
Posterior distributions and associations with factors for the inverse temperature parameter. The parameter represents choice stochasticity, with higher values indicating more deterministic choices. **a**| Posterior distributions for the mean parameter *µ*_0_ of the inverse temperature components, with error bars representing the mean and 95% highest-density intervals (HDI) for each distribution. Significant effects of outcome valence (good vs. bad), task phase (volatile vs. stable), and their interaction were observed across all participants. **b**| The influence of internalizing (general-factor scores) on choice-stochasticity components (inverse temperature), shown by posterior means and 95% HDIs. General-factor scores significantly modulated the baseline inverse temperature, leading to more deterministic choices (*B*_*Baseline*_ *β*_*g*_ = 0.19, 95% HDI = [0.05, 0.34]). Additionally, general-factor scores influenced the difference in choice stochasticity between stable and volatile task phases, leading to more stochastic choices in volatile compared to stable phase (*B*_*V olatile*− *stable*_ *β*_*g*_ = -0.06, 95% HDI = [-0.12, 0.0]). **c**| The anxiety-related factor F1, represented by the population parameter *β*_1_, showed no significant effects on any inverse temperature components. **d**| The depression-related factor F2, represented by the population parameter *β*_2_, was associated with a significant increase in baseline inverse temperature, suggesting reduced choice stochasticity for participants with higher F2 scores

**Figure S21.**
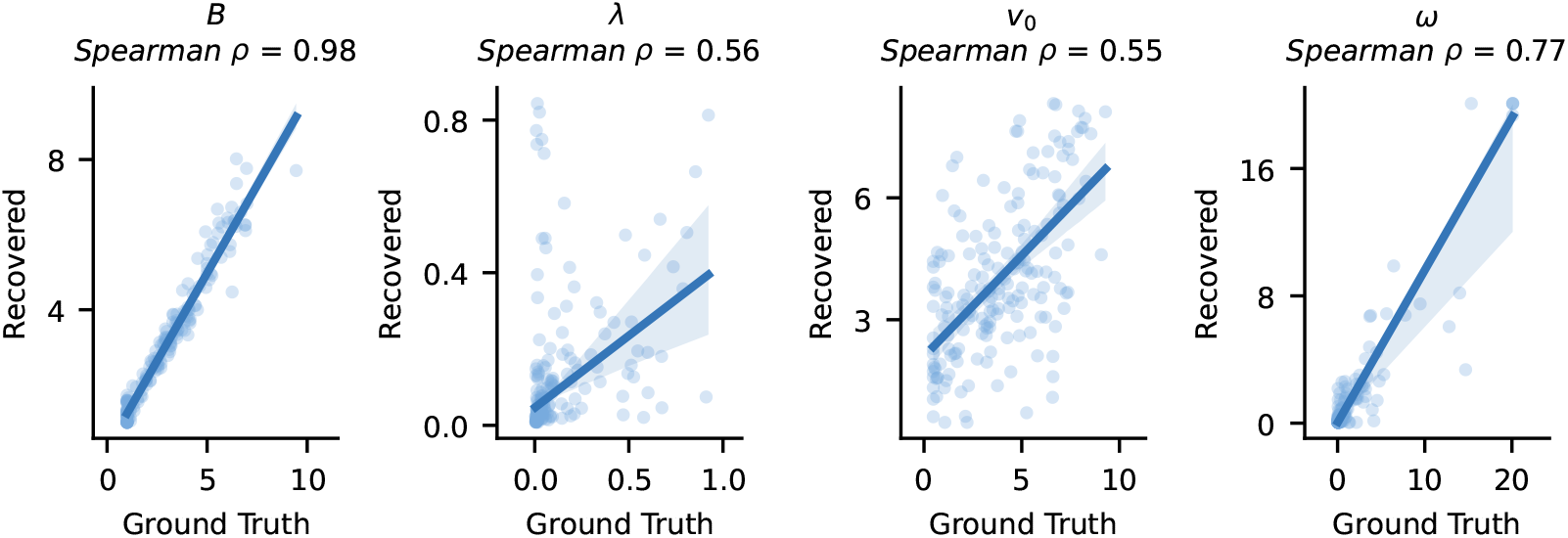
Parameter recovery analysis for the volatile Kalman filter (VKF) model. To assess parameter identifiability, we simulated five datasets using subject-specific parameter values from the VKF. The model was then fitted to each dataset, and Spearman’s rank correlation was computed between the ground truth and recovered parameter values. The figure shows results for one example dataset, with each panel representing a model parameter (x-axis: ground truth parameter values, y-axis: recovered values). The average Spearman correlation between the ground truth and recovered parameters across all 5 simulated datasets was *ρ* = 0.70.

Internalizing psychopathology also modulated the components of the inverse temperature parameter (Fig. S20b). General-factor scores were associated with an increase in baseline inverse temperature, indicating reduced choice stochasticity (*B*_*Baseline*_ *β*_*g*_ = 0.19, 95% HDI = [0.05, 0.34]). Additionally, general-factor scores marginally modulated the difference in choice stochasticity between the stable and volatile task phases, with higher scores linked to more stochastic choices in the volatile phase relative to the stable phase (*B*_*V olatile* − *stable*_ *β*_*g*_ = -0.06, 95% HDI = [-0.12, 0.0]). When examining the anxiety-related factor F1, we did not find significant effects for any inverse temperature components, as all 95% HDIs contained zero (Fig. S20c). In contrast, the depression-related factor F2 was significantly associated with increased baseline inverse temperature, suggesting reduced choice stochasticity for participants with higher F2 scores (*B*_*Baseline*_ *β*_2_ = 0.15, 95% HDI = [0.01, 0.28]; Fig. S20d).

#### Binary Volatile Kalman Filter

As an additional analysis, we fitted a volatile Kalman filter (VKF) to participant behavior in the reversal learning task. The model included 4 free parameters: the volatility update rate *λ*, the initial estimate of volatility *v*_0_, the noise parameter *ω*, and the inverse temperature parameter *B*. The model parameters exhibited moderate-to-good identifiability, with an average correlation across datasets and components of Spearman’s *ρ* = 0.70. An example of parameter recovery from one of the simulated datasets is shown in Fig. S21.

We did not find any significant association of model parameters with internalizing (Fig. S22). Specifically, *λ* was not found to differ significantly between the low- and high-internalizing groups (Welch’s *t*-test *t*_167.79_ = -0.31, *p* = 0.76, *BF*_01_ = 5.814; Fig. S22a) and was not found to be significantly associated with internalizing in a regression analysis (*r* = 0.01, *p* = 0.79, *BF*_01_ = 11.494; Fig. S22b). Similarly, we did not find a significant group difference in *v*_0_ (*t*_168.52_ = 0.82, *p* = 0.41, *BF*_01_ = 4.464; Fig. S22c), with no significant association observed with internalizing in a regression analysis (*r* = 0.05, *p* = 0.54, *BF*_01_ =9.901; Fig. S22d). Likewise, *ω* was not found to be significantly different between the two internalizing groups (*t*_172.04_ = -1.08, *p* = 0.28, *BF*_01_ = 3.546; Fig. S22e), where the regression analysis did not reveal a significant association with internalizing (*r* = 0.01, *p* = 0.29, *BF*_01_ = 6.897; Fig. S22f). The same was true for the inverse temperature parameter *B*, where we did not find a significant group difference (*t*_169.47_ = -0.95, *p* = 0.34, *BF*_01_ = 4.016; Fig. S22g), and no significant association with internalizing in a regression analysis (*r* = 0.13, *p* = 0.08, *BF*_01_ = 2.571; Fig. S22h). These results align with findings from the hierarchical Rescorla-Wagner models, further supporting the hypothesis that internalizing does not impact learning under volatility.

**Figure S22.**
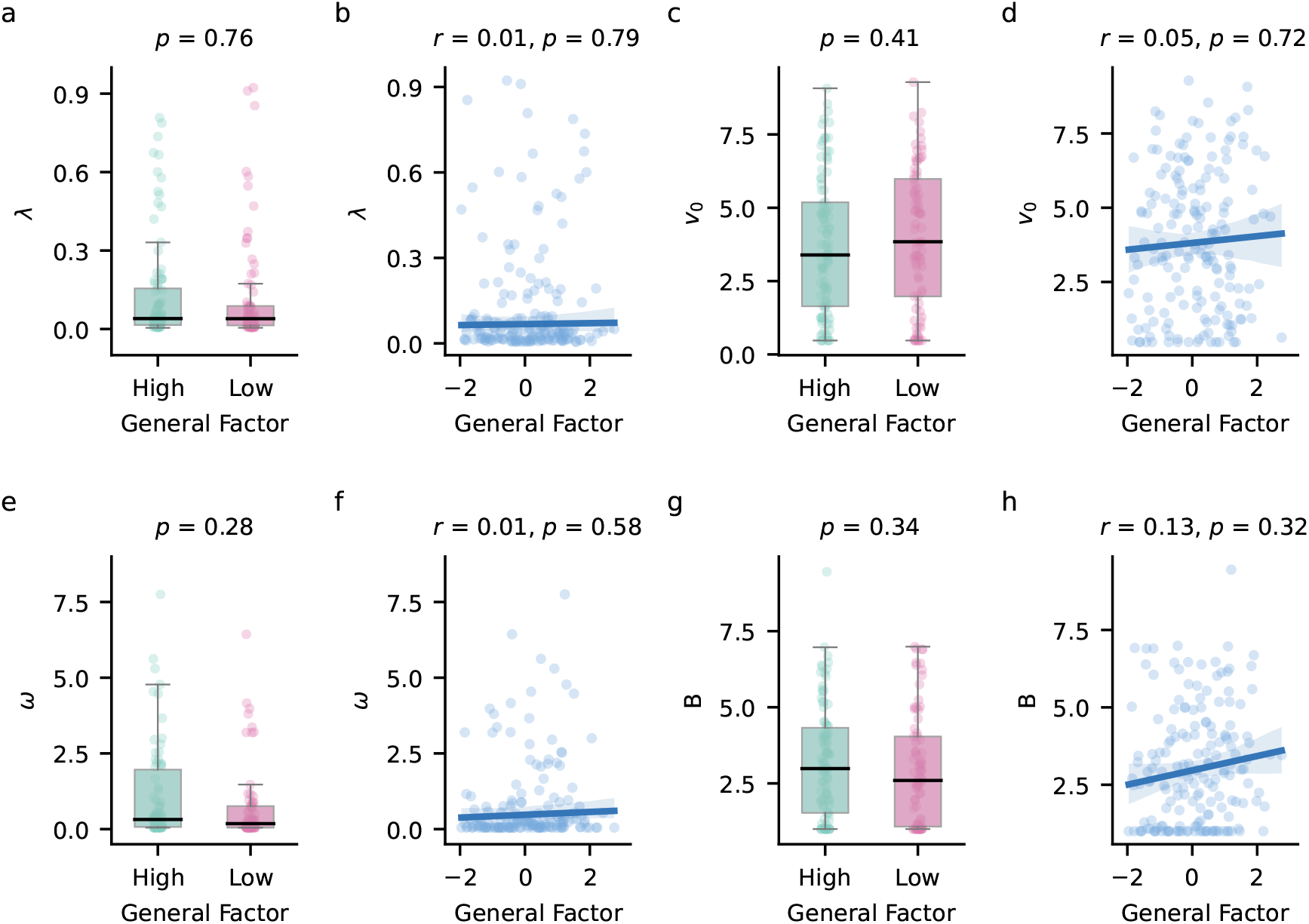
Results from a volatile Kalman filter (VKF) fitted to participant data and analyzed against internalizing. The VKF model fitted to the participant data consisted of 4 free parameters: the volatility update rate *λ*, the initial estimate of volatility *v*_0_, the noise parameter *ω* and the inverse temperature parameter *B*. **a**| The volatility update parameter *λ*, which determines the extent of change in the inferred volatility, was not found to differ significantly between the low- and high-internalizing groups. **b**| Regression analyses did not reveal any significant associations between *λ* and internalizing. **c**| The initial volatility estimate *v*_0_ was not found to differ significantly between the low- and high-internalizing groups. **d**| Regression analyses did not reveal significant associations between *v*_0_ and internalizing. **e**| The noise parameter *ω*, which indicates the scale of volatility throughout the task, was not observed to be significantly different between the low- and high-internalizing groups. **f**| We did not find a significant association of *ω* with internalizing in a regression analysis. **g**| The inverse temperature parameter *B* was not found to be significantly different between the low- and high-internalizing groups in either task phase. **h**| Regression analyses did not reveal any significant association between *B* and internalizing. All regression *p*-values are corrected based on the false discovery rate.

#### Task Version With Reward Magnitudes

In experiment 3, *N* = 94 participants (demographic details in Table S5) completed a version of the binary probabilistic reversal learning task, where the fractals were also associated with varying reward magnitudes. In this version, participants earned points equal to the reward magnitude associated with the chosen fractal on each trial, provided the chosen fractal was the winning fractal for that trial (details in Task Variants).

**Table S5.**
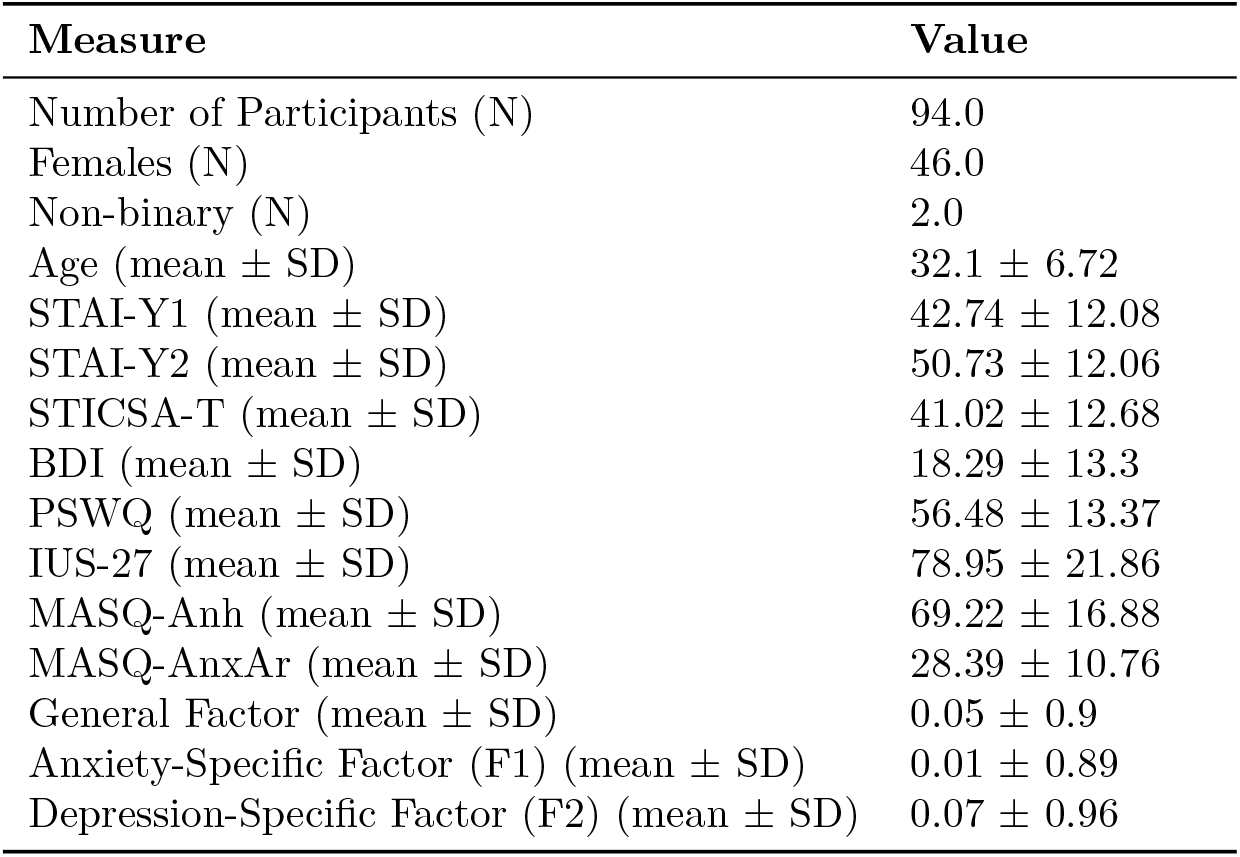
Basic demographic details of participants who completed the binary reversal learning task with varying reward magnitudes. STAI-Y1: Spielberger State-Trait Anxiety Inventory - state scale; STAI-Y2: Spielberger State-Trait Anxiety Inventory - trait scale; STICSA-T: State-Trait Inventory for Cognitive and Somatic Anxiety - trait scale, IUS-27: Intolerance of Uncertainty Scale; BDI: Beck’s Depression Inventory; PSWQ: Penn State Worry Questionnaire; MASQ: Mood and Anxiety Symptom Questionnaire; MASQ-Anh: Mood and Anxiety Symptom Questionnaire - anhedonia subscale; MASQ-AnxAr: Mood and Anxiety Symptom Questionnaire - anxious arousal subscale

#### Descriptive Results

To investigate whether internalizing influenced how participants incorporated reward magnitudes into their decision-making, we first analyzed the total scores achieved in each task phase and examined their associations with internalizing (Fig. S23a and S23b). Since each task phase consisted of two blocks, the total score was calculated as the average of the maximum scores achieved in these blocks. We did not find any significant differences in the total scores when participants were categorized into low- and high-internalizing groups based on their general-factor scores, for either the stable (Welch’s *t*-test *t*_89.94_ = 0.39, *p* = 0.7, *BF*_01_ = 4.274) or volatile task phase (*t*_87.48_ = -1.11, *p* = 0.27, *BF*_01_ = 2.66; Fig. S23a). Regression analyses further confirmed these findings, where we did not find significant associations between total scores and general-factor scores in either the stable phase (*r*(92) = -0.06, *p* = 0.66, *BF*_01_ = 7.519) or the volatile phase (*r*(92) = 0.1, *p* = 0.66, *BF*_01_ = 5.814; Fig. S23b). Additionally, we examined task performance within each phase, and did not find any significant differences between the low- and high-internalizing groups in either task phase (stable phase: *t*_88.22_ = 0.26, *p* = 0.79, *BF*_01_ = 4.444, volatile phase: *t*_86.85_ = -0.89, *p* = 0.38, *BF*_01_ = 3.226; Fig. S23c). Regression analyses also did not reveal significant associations between performance and internalizing scores on a continuum for either the stable (*r*(92) = -0.07, *p* = 0.66, *BF*_01_ = 7.246) or volatile phase (*r*(92) = 0.1, *p* = 0.66, *BF*_01_ = 6.061; Fig. S23d).

Analysis of the percentage of switches following wins and losses did not reveal any significant associations with internalizing. Overall, participants exhibited a higher percentage of switches after losses compared to wins. In the stable phase, the median percentage of switches after wins was 23.3% compared to 46.66% after losses (paired *t*-test *t*_91_ = -9.73, *p* < 0.001, *BF*_01_ = 0). Similar results were found for the volatile phase, where the median percentage of switches after wins was 26.46% compared to 50.0% after losses (*t*_91_ = -10.15, *p* < 0.001, *BF*_01_ = 0).

**Figure S23.**
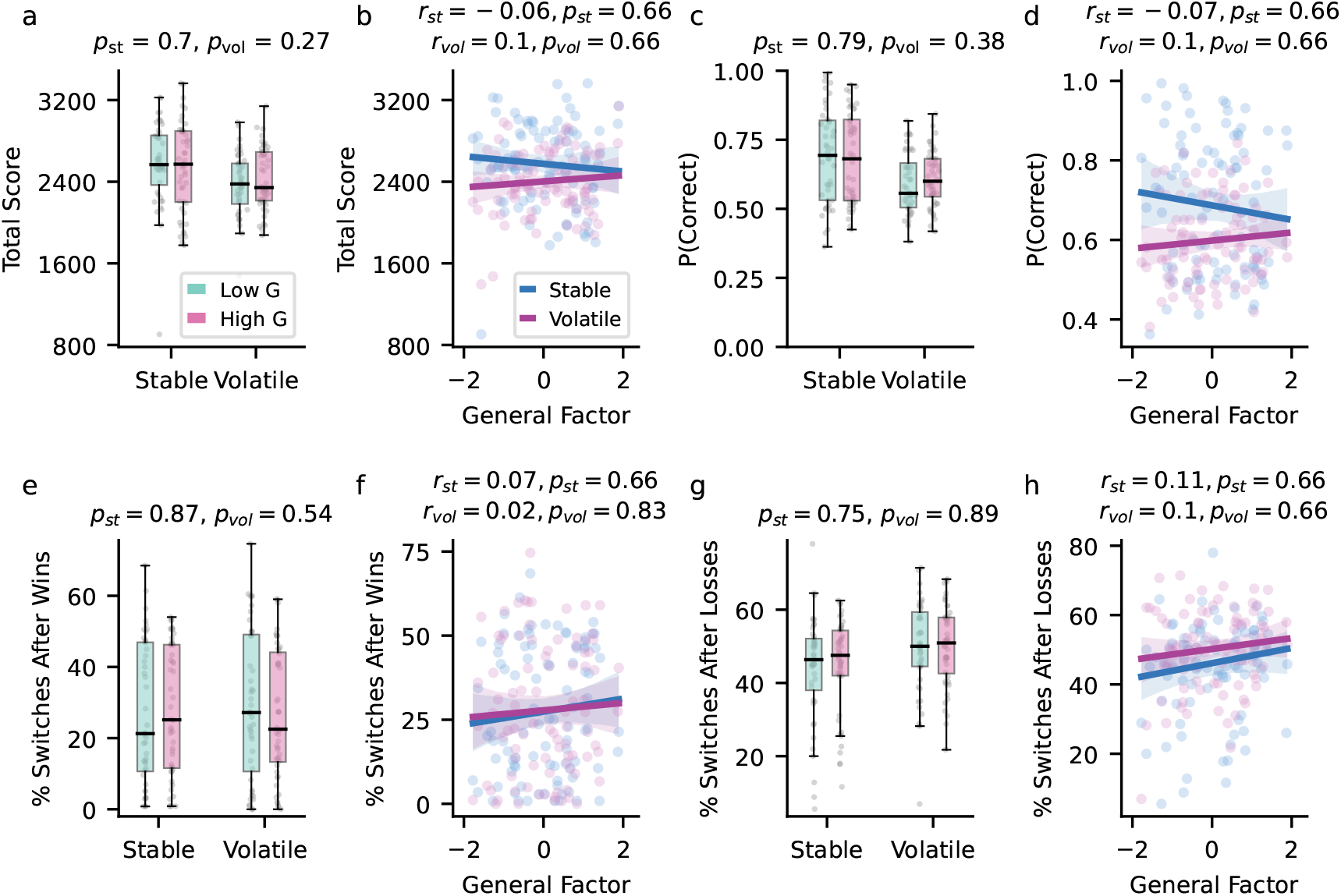
Total score, performance, and percentage of switches in the binary reversal learning task with reward magnitudes. *r*_*st*_ and *p*_*st*_ represent statistics for the stable phase, whereas *r*_*vol*_ and *p*_*vol*_ represent statistics for the volatile task phase. **a**| Total scores, calculated as the average of the maximum scores achieved in each block of a task phase, were not found to be significantly different between the low- and high-internalizing groups across either task phase. **b**| Regression analyses did not reveal any significant associations between total scores and internalizing in either the stable or volatile phase. **c**| Performance, measured as the proportion of trials in which participants selected the rewarding fractal, did not show any significant differences between the low- and high-internalizing groups in either task phase. **d**| Regression analyses did not reveal significant associations between performance and internalizing for either task phase. **e**| The percentage of switches after hits was not found to be significantly different between the low- and high-internalizing groups in either the stable or volatile phase. **f**| Regression analyses did not show significant associations between the percentage of switches after hits and internalizing for either phase. **g**| The percentage of switches after losses was also not found to be significantly different between the low- and high-internalizing groups in either task phase. **h**| Regression analyses did not reveal any significant associations between the percentage of switches after losses and internalizing in either task phase. All regression *p*-values are corrected based on the false discovery rate.

When we categorized participants into low- and high-internalizing groups, we failed to find significant differences in the percentage of switches following wins in either the stable (*t*_87.01_ = -0.17, *p* = 0.87, *BF*_01_ = 4.505) or the volatile phase (*t*_85.25_ = 0.62, *p* = 0.54, *BF*_01_ = 3.861; Fig. S23e). Further regression analyses controlling for age and gender also did not reveal any significant association between internalizing scores and the percentage of switches after wins in either phase (*r*_*stable*_(92) = 0.07, *p*_*stable*_ = 0.66, *BF* 01_*stable*_ = 7.299, *r*_*volatile*_(92) = 0.02, *p*_*volatile*_ = 0.83, *BF* 01_*volatile*_ = 8.475; Fig. S23f). Similarly, we did not find a significant difference in the percentage of switches after losses between the low- and high-internalizing groups in either the stable (*t*_85.73_ = -0.32, *p* = 0.75, *BF*_01_ = 4.367) or the volatile phase (*t*_85.16_ = -0.14, *p* = 0.89, *BF*_01_ = 4.525; Fig. S23g). Additionally, we did not observe any significant association between the percentage of switches after losses and internalizing, for either the stable (*r*_*stable*_(92) = 0.11, *p*_*stable*_ = 0.66, *BF*_01_ = 4.484) or volatile phase, controlling for age and gender (*r*_*volatile*_(92) = 0.1, *p*_*volatile*_ = 0.66, *BF*_01_ = 5.495; Fig. S23h).

**Table S6.** Model comparison for the probabilistic reversal learning task with reward magnitudes. *α*: Learning-rate parameter, *β*: Inverse temperature parameter, *γ*: Risk parameter, *λ*: Mixture parameter, *r*: Magnitude-scaling parameter, *δ*: Decay parameter, *ϵ*: Lapse parameter. The subscript *ck* represents the choice-kernel parameter. Superscripts indicate parameter components: *sv*: Baseline component and a task-phase-dependent component (stable vs. volatile). *gb*: Component encoding outcome valence (good vs. bad). *sv · gb*: Interaction between task phase and outcome valence. PSIS-LOO: Pareto-smoothed importance-sampling-based approximation of leave-one-out cross-validation. *SE*: standard error.

| Model Number | Parameters | # of Parameter Components | PSIS-LOO (SE) |
| --- | --- | --- | --- |
| Model 1 | $\alpha^{sv}, \gamma^{sv}, \beta^{sv}$ | 6.0 | 29320.81 (167.71 ) |
| Model 2 | $\alpha^{sv}, \lambda^{sv}, \beta^{sv}$ | 6.0 | 29072.52 (174.82 ) |
| Model 3 | $\alpha^{sv,gb,sv \cdot gb}, \lambda^{sv}, \beta^{sv}$ | 8.0 | 28263.58 (176.86) |
| Model 4 | $\alpha^{sv}, \lambda^{sv,gb,sv \cdot gb}, \beta^{sv,gb,sv \cdot gb}$ | 10.0 | 28644.95 (173.43) |
| Model 5 | $\alpha^{sv,gb,sv \cdot gb}, \lambda^{sv,gb,sv \cdot gb}, \beta^{sv,gb,sv \cdot gb}$ | 12.0 | 28009.58 (175.65) |
| Model 6 | $\alpha^{sv,gb,sv \cdot gb}, \lambda^{sv,gb,sv \cdot gb}, \beta^{sv,gb,sv \cdot gb}, r$ | 13.0 | 27648.36 (175.06) |
| Model 7 | $\alpha^{sv,gb,sv \cdot gb}, \lambda^{sv,gb,sv \cdot gb}, \beta^{sv,gb,sv \cdot gb}, r, \epsilon$ | 14.0 | 27649.14 (174.81) |
| Model 8 | $\alpha^{sv,gb,sv \cdot gb}, \lambda^{sv,gb,sv \cdot gb}, \beta^{sv,gb,sv \cdot gb}, r, \delta^{sv}$ | 15.0 | 27580.33 (173.28) |
| **Model 9** | $\alpha^{sv,gb,sv \cdot gb}, \alpha_{ck}, \lambda^{sv,gb,sv \cdot gb}, \beta^{sv,gb,sv \cdot gb}, \beta_{ck}, r$ | 15.0 | **27301.15 (176.35)** |
| Model 10 | $\alpha^{sv,gb,sv \cdot gb}, \alpha_{ck}, \lambda^{sv,gb,sv \cdot gb}, \beta^{sv,gb,sv \cdot gb}, \beta_{ck}, r, \delta^{sv}$ | 17.0 | 27364.28 (175.85) |

#### Model Comparison

We augmented the models fitted to the previous version of the binary reversal learning task by incorporating the effect of reward magnitudes into the models. Model fit was evaluated using PSIS-LOO (Table S6). The winning model was Model 9, consisting of 15.0 parameters and having a PSIS-LOO value of 27301.15 (SE = 176.35).

#### Parameter Recovery

We followed the parameter recovery approach described above (Parameter Recovery). Recovery results indicated good parameter recovery, with an average correlation for the learning-rate parameters across datasets and components of *ρ* = 0.66, and an overall average correlation across all components of *ρ* = 0.71. An example of parameter recovery from one of the simulated datasets is shown in Fig. S24.

**Figure S24.**
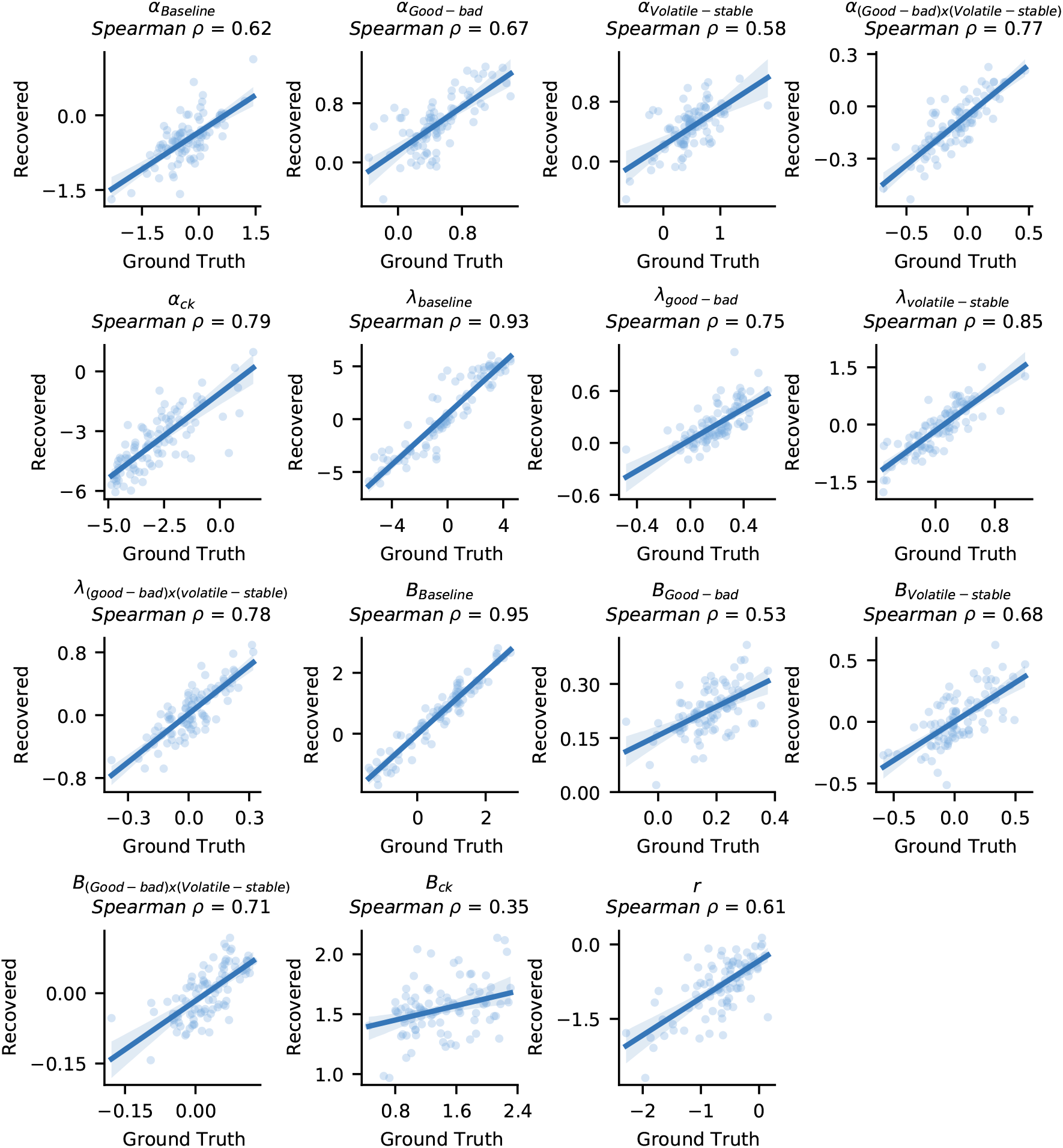
Parameter recovery analysis for the winning model (Model 9). The average Spearman correlation between the ground truth and recovered parameters across all 5 simulated datasets was *ρ* = 0.71.

**Figure S25.**
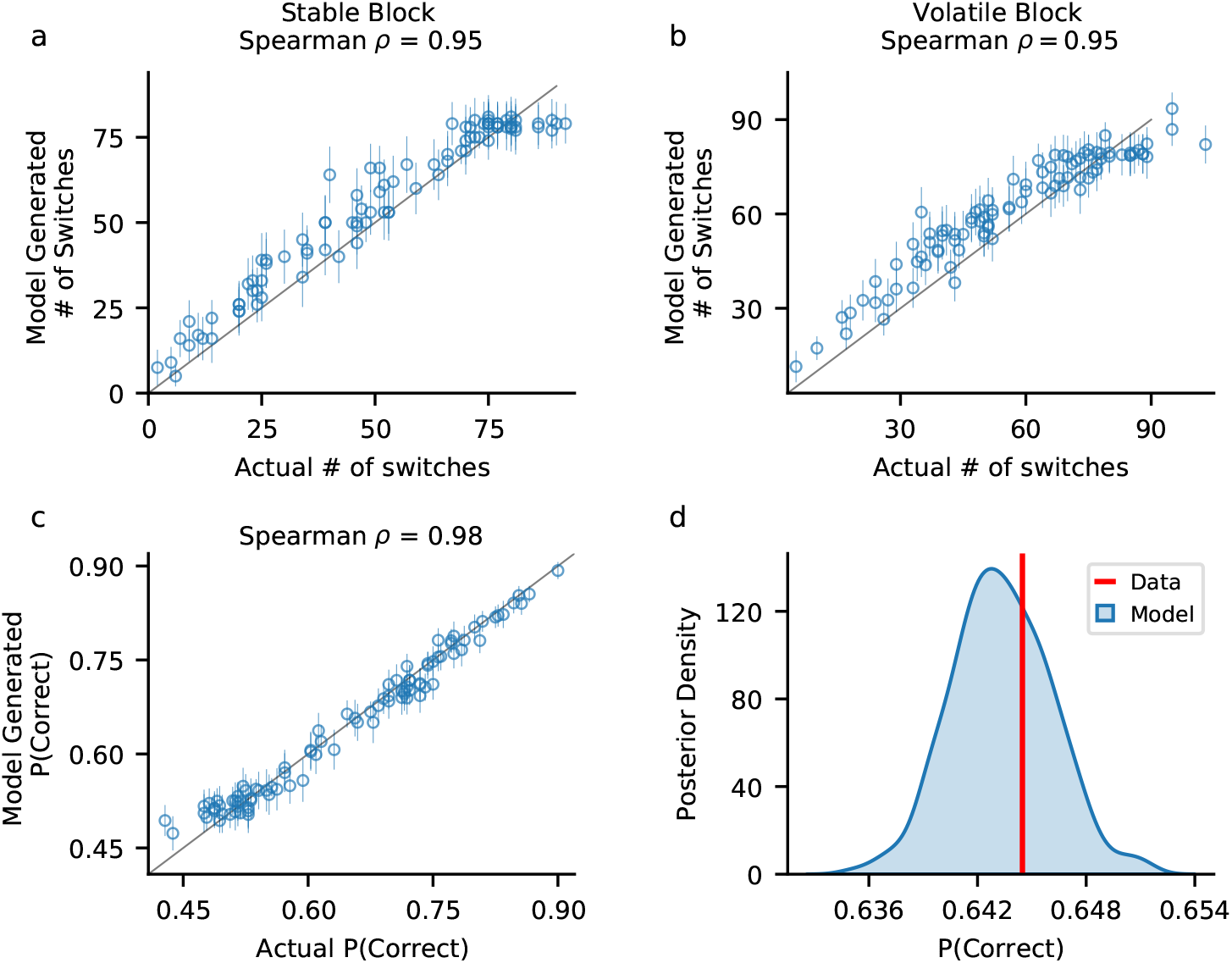
Posterior predictive checks for the winning model. To assess how well the winning model (Model 9) captured key qualitative features of choice behavior, we generated 500 simulated choice datasets per participant by drawing random samples from each participant’s joint posterior distribution (320 trials per dataset). **a-b**| Spearman correlations between the average number of switches across simulations and actual participant switches in the stable (**a**) and volatile (**b**) task phases were strong, indicating that the model successfully reproduces switching behavior. Circles represent the average number of switches, and error bars denote the standard deviation across simulations. **c**| We found a high correlation between the proportion of trials in which the high-reward fractal was chosen (P(Correct)) in the simulated data and actual participant performance. This suggests that the model captured choice accuracy accurately. **d**| Distribution of the model’s predicted performance across trials and participants. The red line represents the actual average P(Correct) across participants. The close alignment between the predicted distribution and the actual value further supports the model’s ability to replicate key behavioral patterns.

#### Model Reproduction of Basic Features

We used the procedure described in SM Model Reproduction of Basic Features to assess how well the winning model’s posterior predictions captured key qualitative aspects of participant behavior, focusing on switching patterns and choice accuracy in model-generated data (Fig. S25). The number of switches in the simulated datasets closely matched the actual participant data, showing strong Spearman correlations in both the stable (*ρ* = 0.95, Fig. S25a) and volatile (*ρ* = 0.95, Fig. S25b) task phases. Similarly, the model successfully reproduced choice accuracy, with a high correlation between simulated and actual P(Correct) (Spearman *ρ* = 0.98, Fig. S25c). Furthermore, the distribution of mean P(Correct) in the simulated data closely aligned with the actual average P(Correct) across participants (Fig. S25d). These results indicate that the winning model was able to effectively capture key behavioral patterns observed in participant choices.

**Figure S26.**
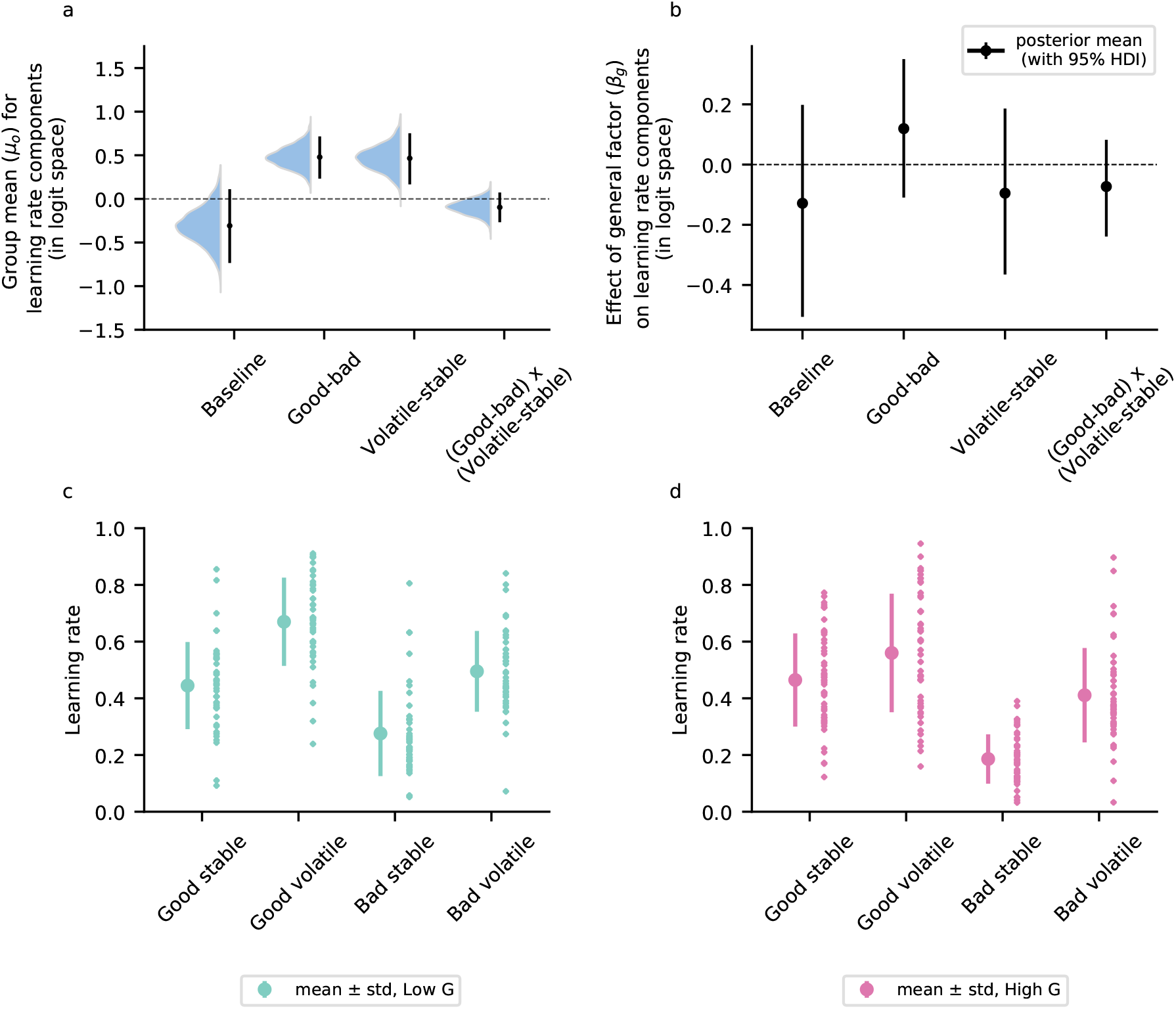
Mean learning-rate components and the effect of internalizing on learning-rate components for the reversal learning task with reward magnitudes. **a**| Mean learning-rate components across all participants. Learning rates were significantly higher after positive outcomes compared to negative outcomes and during the volatile phase compared to the stable phase. **b**| We did not find an effect of internalizing, represented by the population-level parameter *β*_*g*_, on learning-rate components, with the 95% highest-density intervals (HDIs) of all components containing zero. **c**,**d**| Participants were divided into low- and high-internalizing groups based on their general-factor scores (below and above the mean). We extracted their mean learning rates for the interaction between valence and task phase, visualizing the learning rate for the two groups.

#### Model Results

Across participants, we observed a significant effect of outcome valence (good vs. bad) and task phase (volatile vs. stable) on learning rates (Fig. S26a). Specifically, learning rates were higher after good outcomes compared to bad outcomes (*α*_*Good* − *bad*_ *µ*_0_ = 0.48, 95% HDI = [0.23, 0.72]). Similarly, participants had a higher learning rate in the volatile phase compared to the stable phase (*α*_*V olatile* − *stable*_ *µ*_0_ = 0.47, 95% HDI = [0.17, 0.75]). However, we did not find a significant interaction between outcome valence and task phase (*α*_(*Good*− *bad*)*x*(*V olatile*− *stable*)_ *µ*_0_ = -0.1, 95% HDI = [-0.27, 0.07]).

We did not find a significant effect of internalizing on any of the learning-rate components (Fig. S26b), with all the 95% HDI’s containing zero (*α*_*Baseline*_ *β*_*g*_ = -0.13, 95% HDI = [-0.51, 0.2]; *α*_*Good*−*bad*_ *β*_*g*_ = 0.12, 95% HDI = [-0.11, 0.35]; *α*_*V olatile* − *stable*_ *β*_*g*_ = -0.1, 95% HDI = [-0.37, 0.19]; *α*_(*Good*− *bad*)*x*(*V olatile* − *stable*)_ *β*_*g*_ = -0.07, 95% HDI = [-0.24, 0.08]). Furthermore, dividing participants into low- and high-internalizing groups based on their general-factor scores and visualizing their learning rates for the different valence and task-phase conditions, did not reveal qualitative differences between the groups, that is, learning from good and bad outcomes in either stable or volatile phase appeared comparable across participants (Fig. S26c and S26d). Similarly, analyses of the anxiety-related factor F1 and the depression-related factor F2 did not revealed a significant associations with any learning-rate component (all 95% HDIs contained zero; Fig. S27a and S27b).

**Figure S27.**
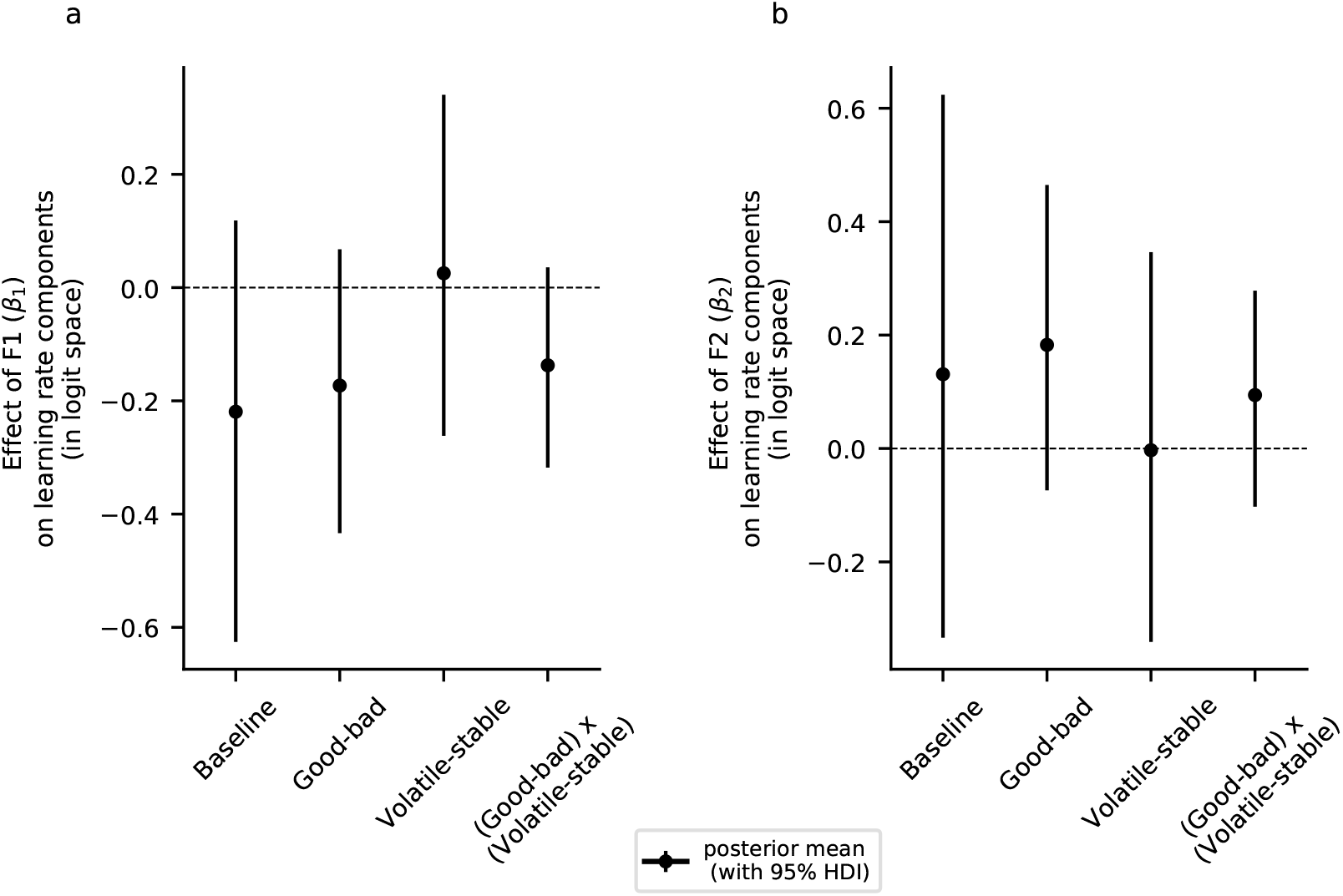
Effects of anxiety-related factor (F1) and depression-related factor (F2) on learning rates in the reversal learning task with reward magnitudes. **a**,**b**| We did not find an effect of either the anxiety-related factor (F1; represented by the *β*_1_ population-level parameter) (**a**) or the depression-related factor (F2; represented by the *β*_2_ population-level parameter) (**b**) on any learning-rate component, with all corresponding 95% highest-density intervals (HDIs) containing zero.

Next, we analyzed the inverse temperature parameter and its association with internalizing, anxiety, and depression (Fig. S28). Across participants, we found a significant effect of outcome valence (good vs. bad outcome) on the inverse temperature, and participants showed higher parameter values after good outcomes compared to bad outcomes (*B*_*Good*− *bad*_ *µ*_0_ = 0.17, 95% HDI = [0.12, 0.24]). This finding suggests that participants made less stochastic choices after rewards than non-rewards. However, we did not find a significant effect of task phase on the inverse temperature, suggesting that choice stochasticity may be similar in the volatile and the stable phase (*B*_*V olatile* − *stable*_ *µ*_0_ = 0.03, 95% HDI = [-0.05, 0.12]). Similarly, we did not find a significant interaction of outcome valence and task phase on the inverse temperature (*B*_(*Good* − *bad*)*x*(*V olatile* − *stable*)_ *µ*_0_ = 0.02, 95% HDI = [-0.02, 0.07], Fig. S28a). Additionally, we did not find a significant effect of internalizing (Fig. S28b), the anxiety-related factor (Fig. S28c), or the depression-related factor (Fig. S28d) on the inverse temperature parameter. These results suggest that the previously observed effects of internalizing and depression-related factors on choice stochasticity in the task without reward magnitudes might disappear when reward magnitudes are included. This highlights the specificity of these effects to particular task contexts and raises questions about their generalizability.

**Figure S28.**
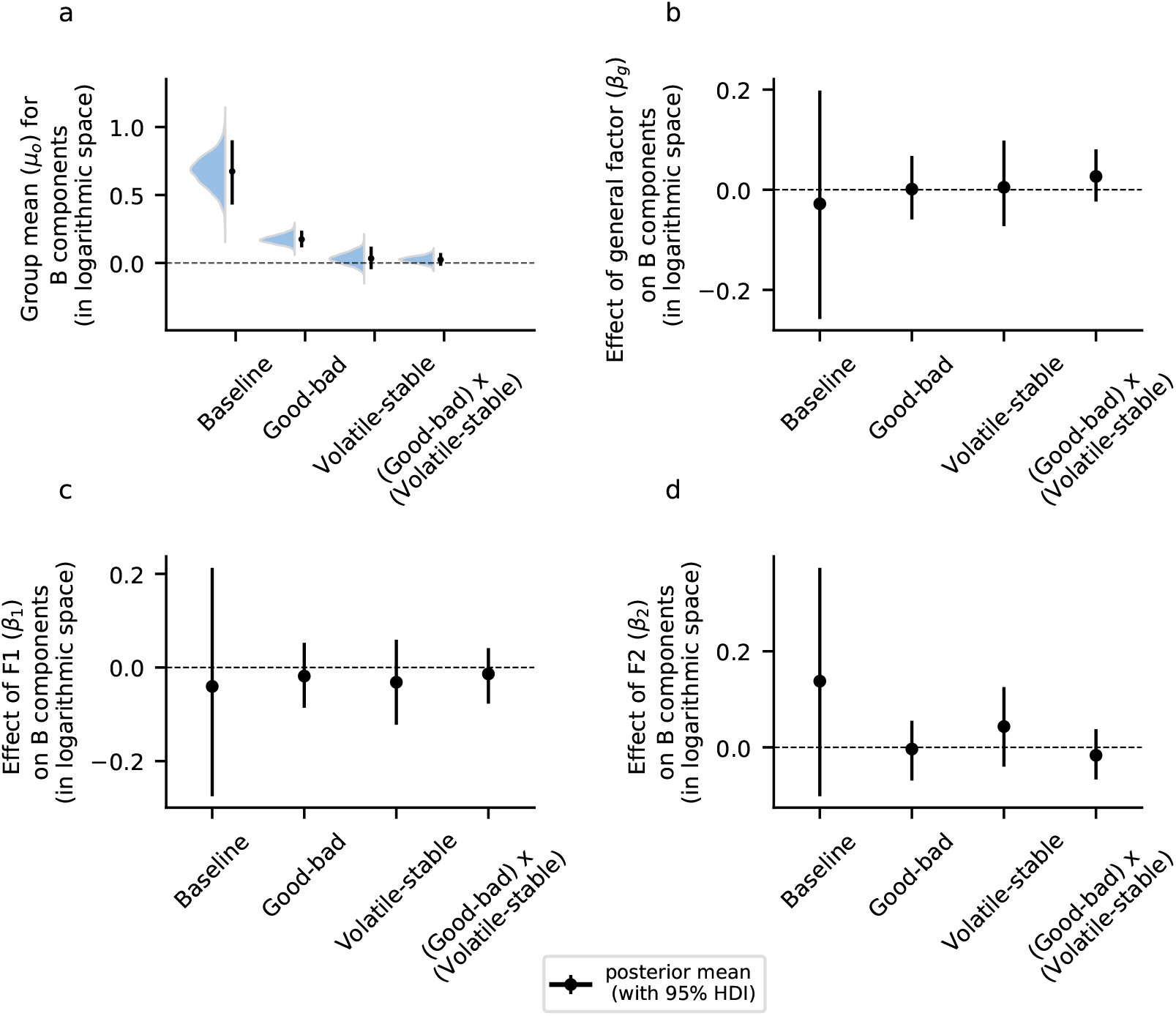
Mean inverse temperature and effect of internalizing, the anxiety-related factor F1, and the depression-related factor F2 on the inverse temperature parameter in the reversal learning task with reward magnitudes. **a**| Group mean (*µ*_0_, population-level parameter) of inverse temperature *B* across participants. A significant effect of outcome valence was observed, with inverse temperature increasing after good outcomes compared to bad outcomes. **b**| Effect of internalizing (*β*_*g*_, population-level parameter) on inverse temperature. We did not find a significant effect of internalizing on any of the inverse temperature components, with all corresponding 95% highest-density intervals (HDIs) containing zero. **c**| We did not observe an effect of the anxiety-related factor (F1, represented by the *β*_1_ population-level parameter) on any inverse temperature component, with all corresponding 95% HDIs containing zero. **d**| Similarly, we did not observe an effect of the depression-related factor (F2, represented by the *β*_2_ population-level parameter) on any inverse temperature component, with all corresponding 95% HDIs containing zero.

#### Task Version with Outcome Magnitudes – Reward and Loss Domain

In experiment 4, *N* = 161 participants (demographic details in Table S7) completed a binary probabilistic reversal learning task with reward magnitudes, where the task was additionally divided into two distinct domains: a reward domain and a loss domain. In the reward domain, each fractal was associated with a specific reward magnitude, and participants received either the displayed reward or no reward depending on whether their chosen fractal was the correct one in that trial. Conversely, in the loss domain, each fractal was associated with a loss magnitude. Participants began with an initial tally of 10,000 points and lost the corresponding magnitude if they selected an incorrect fractal, whereas selecting the correct fractal prevented any point deduction. This task, consisting of the same reward probability and reward magnitude schedules as Gagne et al. (2020), aimed to replicate their results and to investigate whether the nature of the domain (reward versus loss) influences learning processes and the interplay of internalizing in these learning processes.

**Table S7.** Basic demographic details of participants who completed the reversal learning task with outcome magnitudes, consisting of reward and loss domains. STAI-Y1: Spielberger State-Trait Anxiety Inventory - state scale; STAI-Y2: Spielberger State-Trait Anxiety Inventory - trait scale; STICSA-T: State-Trait Inventory for Cognitive and Somatic Anxiety - trait scale, IUS-27: Intolerance of Uncertainty Scale; BDI: Beck’s Depression Inventory; PSWQ: Penn State Worry Questionnaire; MASQ: Mood and Anxiety Symptom Questionnaire; MASQ-Anh: Mood and Anxiety Symptom Questionnaire - anhedonia subscale; MASQ-AnxAr: Mood and Anxiety Symptom Questionnaire - anxious arousal subscale

| Measure | Value |
| --- | --- |
| Number of Participants (N) | 161.0 |
| Females (N) | 80.0 |
| Non-binary (N) | 0.0 |
| Age (mean ± SD) | 31.87 ± 6.89 |
| STAI-Y1 (mean ± SD) | 43.12 ± 14.03 |
| STAI-Y2 (mean ± SD) | 50.23 ± 13.45 |
| STICSA-T (mean ± SD) | 40.98 ± 13.75 |
| BDI (mean ± SD) | 19.62 ± 14.19 |
| PSWQ (mean ± SD) | 54.42 ± 15.16 |
| IUS-27 (mean ± SD) | 81.42 ± 23.96 |
| MASQ-Anh (mean ± SD) | 68.41 ± 18.73 |
| MASQ-AnxAr (mean ± SD) | 29.89 ± 10.94 |
| General Factor (mean ± SD) | 0.07 ± 1.06 |
| Anxiety-Specific Factor (F1) (mean ± SD) | 0.14 ± 1.07 |
| Depression-Specific Factor (F2) (mean ± SD) | -0.14 ± 0.99 |

#### Descriptive Results

We analyzed total scores, performance, and percentage of switches separately for the reward and loss domains of the task (Figs. S29 and S30). For the reward domain, we did not find a significant difference in the total score between low- and high-internalizing groups in either the stable (Welch’s *t*-test *t*_156.44_ = -0.12, *p* = 0.91, *BF*_01_ = 5.848) or volatile task phase (*t*_157.84_ = -1.2, *p* = 0.74, *BF*_01_ = 3.03; Fig. S29a).

A regression analysis also did not reveal significant associations between internalizing and total scores in the stable phase (*r*(161) = 0.04, *p* = 0.86, *BF*_01_ = 10.204) or the volatile phase (*r*(161) = 0.11, *p* = 0.74, *BF*_01_ = 4.926; Fig. S29b). Similarly, performance, measured as the proportion of trials in which participants chose the high rewarding fractal, did not show any significant differences between the low- and high-internalizing groups for either task phase (stable phase: *t*_158.84_ = -0.47, *p* = 0.86, *BF*_01_ = 5.319, volatile phase: *t*_158.42_ = -0.32, *p* = 0.86, *BF*_01_ = 5.618; Fig. S29c). These results were further supported by the regression analysis, where we did not find any significant associations between internalizing and performance in either the stable phase (*r*(161) = 0.07, *p* = 0.86, *BF*_01_ = 7.407) or the volatile phase (*r*(161) = -0.03, *p* = 0.86, *BF*_01_ = 10.309; Fig. S29d). Next, we analyzed the percentage of switches after rewards (successful trials) or no rewards (failed trials). We did not find a significant difference in the percentage of switches following rewards between the low- and high-internalizing groups, for either the stable or volatile task phase (stable phase: *t*_158.91_ = 0.89, *p* = 0.86, *BF*_01_ = 4.082, volatile phase: *t*_156.57_ = 0.39, *p* = 0.86, *BF*_01_ = 5.464; Fig. S29e). Similarly, regression analysis did not reveal any significant association between internalizing and switch percentage after rewards, for either the stable (*r*(161) = -0.11, *p* = 0.74, *BF*_01_ = 5.376) or the volatile phase (*r*(161) = -0.03, *p* = 0.86, *BF*_01_ = 10.638; Fig. S29f). We also did not find a significant group difference in the percentage of switches following no reward, in either the stable and volatile phases (stable phase: *t*_151.57_ = -0.13, *p* = 0.91, *BF*_01_ = 5.848, volatile phase: *t*_158.23_ = -2.08, *p* = 0.32, *BF*_01_ = 0.81; Fig. S29g). However, regression analyses revealed a significant association between internalizing and switches following no rewards in the volatile phase (*r*(161) = 0.29, *p* < 0.001, *BF*_01_ = 0.004; Fig. S29h), indicating that participants with higher internalizing exhibited increased switching behavior after failed outcomes in this phase. This finding contrasts with the results from experiment 3, which also employed a reversal learning task involving reward magnitudes. In that experiment, no significant increase in switch percentages following failed trials was observed with internalizing. This discrepancy underscores the challenges of achieving consistent outcomes across similar tasks.

**Figure S29.**
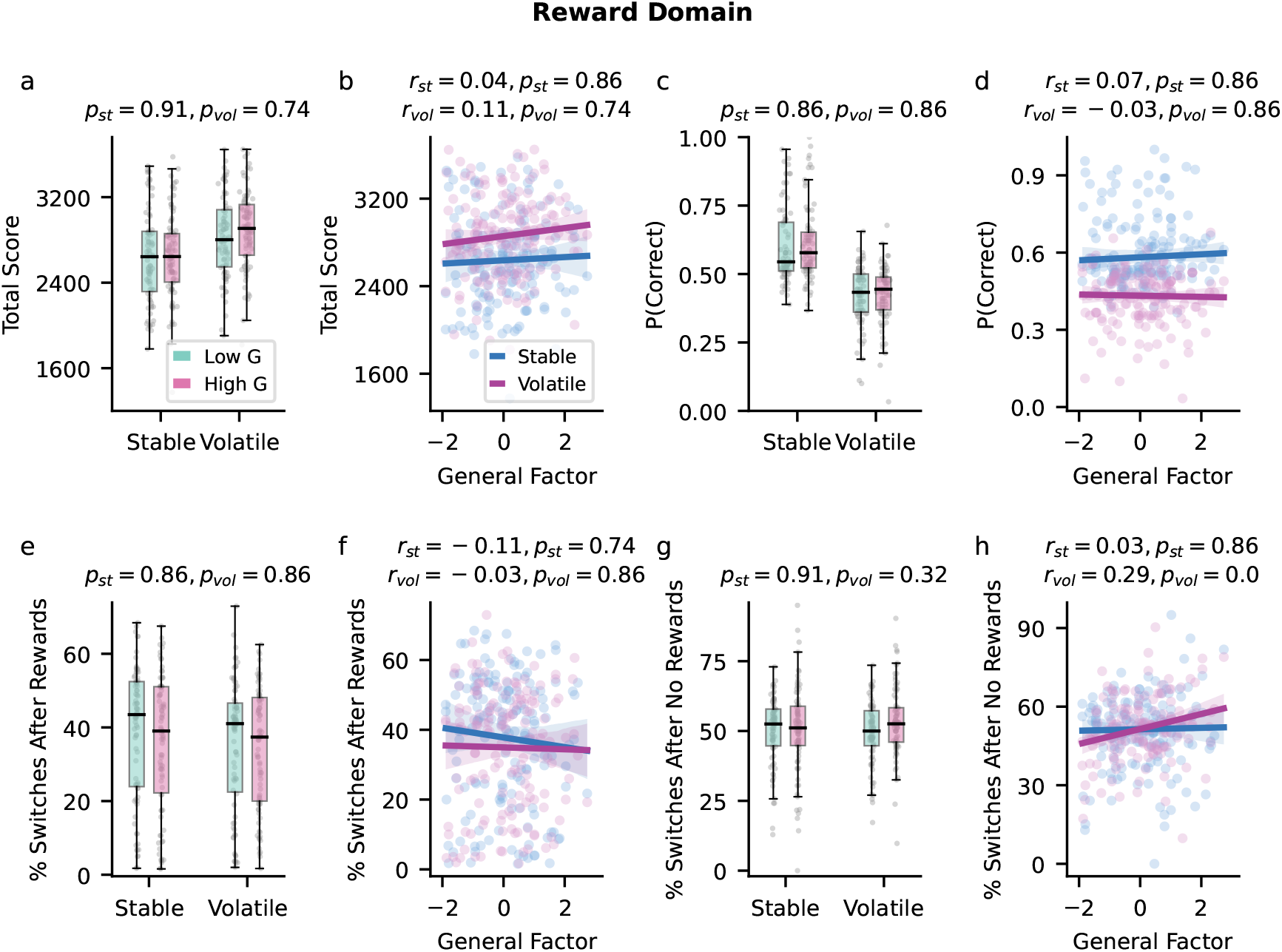
Total score, performance, and percentage of switches in the reward domain of the binary reversal learning task with reward and loss domain. Here, *r*_*st*_ and *p*_*st*_ denote the stable phase statistics, while *r*_*vol*_ and *p*_*vol*_ pertain to the volatile phase. **a**| We did not observe a significant difference in total scores between the low- and high-internalizing groups across either task phase. **b**| Regression analyses did not reveal any significant associations between total scores and internalizing in either the stable or volatile phase. **c**| Performance, measured as the proportion of trials in which participants selected the rewarding fractal, did not show any significant differences between the low- and high-internalizing groups in either task phase. **d**| Regression analyses found no significant associations between performance and internalizing for either task phase. **e**| We did not find any significant differences in the percentage of switches following rewards (successful trials) between the low- and high-internalizing groups in either the stable or volatile phase. **f**| Regression analyses did not reveal any significant associations between the percentage of switches after rewards and internalizing for either phase. **g**| The percentage of switches after no rewards (failed trials) was also not found to be significantly different between the low- and high-internalizing groups in either task phase. **h**| Regression analyses revealed a significant association between internalizing and the percentage of switches after no rewards in the volatile task phase. All *p*-values are corrected for multiple comparisons using false discovery rate correction.

**Figure S30.**
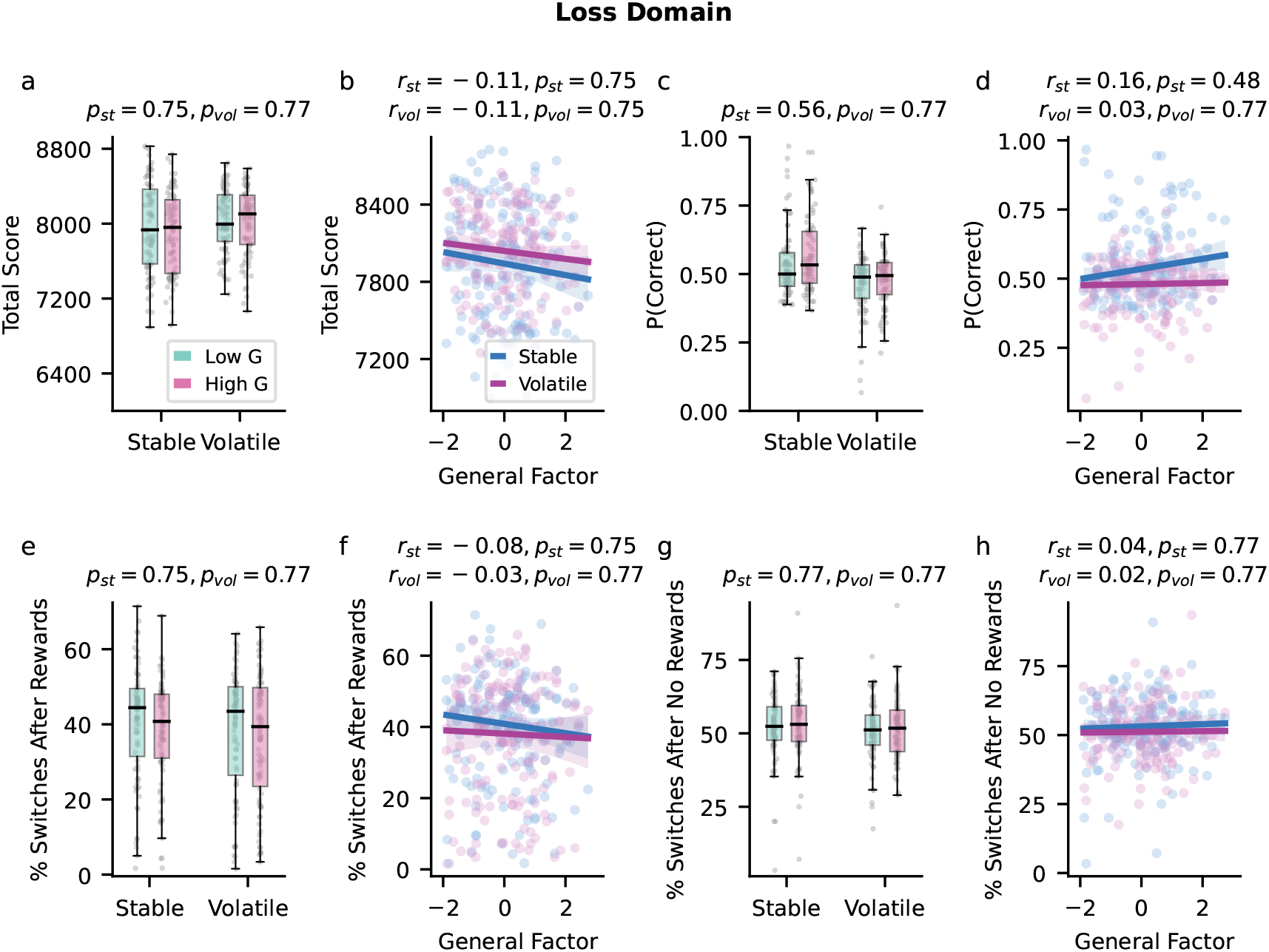
Total score, performance, and switching behavior in the loss domain of the binary reversal learning task of experiment 4. Participants began the task with an initial score of 10,000 points, which decreased after failed trials in the loss domain. Subscripts *st* and *vol* denote the stable and volatile task phases, respectively. **a**| We did not observe any significant differences in total scores between the low- and high-internalizing groups across either task phase. **b**| Regression analyses did not reveal any significant associations between total scores and internalizing in either the stable or volatile phase. **c**| Performance, measured as the proportion of trials in which participants selected the rewarding fractal, was not found to be significantly different between the low- and high-internalizing groups in either task phase. **d**| Regression analyses did not reveal any significant associations between performance and internalizing for either task phase. **e**|We did not find any significant difference in the percentage of switches following no-loss trials (successful trials) between the low- and high-internalizing groups in either the stable or volatile phase. **f**| Regression analyses did not reveal any significant associations between the percentage of switches after no-loss trials and internalizing for either phase. **g**| The percentage of switches after loss trials (failed trials) was also not found to be significantly different between the low- and high-internalizing groups in either task phase. **h**| Regression analyses did not reveal a significant association between the percentage of switches after loss trials and internalizing in either task phase. All *p*-values are corrected for multiple comparisons using false discovery rate correction.

For the loss domain, we did not find any significant associations of internalizing with total scores, performance, or switching behavior (Fig. S30). Participants started this task with an initial score of 10,000 points, which reduced after every loss. We did not find a significant difference in the final participant score between the low- and high-internalizing groups, for either task phase (stable phase: Welch’s *t*-test *t*_155.74_ = 0.99, *p* = 0.75, *BF*_01_ = 3.745, volatile phase: *t*_158.7_ = 0.31, *p* = 0.77, *BF*_01_ = 5.618; Fig. S30a). Similarly, in a regression analysis, we did not find a significant association of the final score with internalizing in the stable phase (*r*(161) = -0.11, *p* = 0.75, *BF*_01_ = 4.926) and the volatile phase (*r*(161) = -0.11, *p* = 0.75, *BF*_01_ = 4.854; Fig. S30b). The analysis also did not reveal a significant difference in performance between the two internalizing groups for either task phase (stable phase: *t*_158.65_ = -1.8, *p* = 0.56, *BF*_01_ = 1.328, volatile phase: *t*_154.43_ = -0.77, *p* = 0.77, *BF*_01_ = 4.464; Fig. S30c). Regression of performance with internalizing also did not reveal any significant association in either the stable (*r*(161) = 0.16, *p* = 0.48, *BF*_01_ = 1.139) or the volatile phase (*r*(161) = 0.03, *p* = 0.77, *BF*_01_ = 10.526; Fig. S30d). Analysis of switch percentage following no loss (successful trials) did not reveal any significant difference between the two internalizing groups, for either the stable or volatile phase (stable phase: *t*_156.01_ = 1.0, *p* = 0.75, *BF*_01_ = 3.704, volatile phase: *t*_159.0_ = 0.58, *p* = 0.77, *BF*_01_ = 5.025; Fig. S30e).

Similarly, regression analysis did not reveal any significant association between internalizing and switch percentage following no-loss trials for either the stable (*r*(161) = -0.08, *p* = 0.75, *BF*_01_ = 7.092) or the volatile phase (*r*(161) = -0.03, *p* = 0.77, *BF*_01_ = 10.638 Fig. S30f). This pattern was also found in the percentage of switches following losses, where we did not find a significant difference between the low- and high-internalizing groups, in either task phase (stable phase: *t*_158.95_ = -0.69, *p* = 0.77, *BF*_01_ = 4.717, volatile phase: *t*_158.99_ = -0.45, *p* = 0.77, *BF*_01_ = 5.348; Fig. S30g). Similarly, regression analysis did not reveal a significant association between internalizing and switch percentages following failed trials, in either the stable (*r*(161) = 0.04, *p* = 0.77, *BF*_01_ = 9.174) or the volatile phase (*r*(161) = 0.02, *p* = 0.77, *BF*_01_ = 10.87; Fig. S30h). This result contrasts with findings from the reward domain, where participants exhibited increased switching following losses during the volatile phase.

**Table S8.** Model comparison for the probabilistic reversal learning task with reward and loss domains. *α*: Learning-rate parameter, *β*: Inverse temperature parameter, *γ*: Risk parameter, *λ*: Mixture parameter, *r*: Magnitude-scaling parameter, *δ*: Decay parameter, *ϵ*: Lapse parameter. The subscript *ck* represents the choice-kernel parameter. Superscripts indicate parameter components: *sv*: Baseline component and a task phase-dependent component (stable vs. volatile). *rl*: Component encoding task domain (reward vs. loss). *gb*: Component encoding outcome valence (good vs. bad). *sv· gb*: Interaction between task phase and outcome valence. *sv · rl*: Interaction between task phase and task domain. *rl · gb*: Interaction between task domain and outcome valence. PSIS-LOO: Pareto-smoothed importance-sampling-based approximation of leave-one-out cross-validation. *SE*: standard error.

| Model Number | Parameters | # of Parameter Components | PSIS-LOO (SE) |
| --- | --- | --- | --- |
| Model 1 | $\alpha^{sv,rl,sv \cdot rl}, \gamma^{sv,rl,sv \cdot rl}, \beta^{sv,rl,sv \cdot rl}$ | 12.0 | 70878.07 (165.4) |
| Model 2 | $\alpha^{sv,rl,sv \cdot rl}, \lambda^{sv,rl,sv \cdot rl}, \beta^{sv,rl,sv \cdot rl}$ | 12.0 | 65773.8 (203.4) |
| Model 3 | $\alpha^{sv,rl,gb,sv \cdot gb,sv \cdot rl,rl \cdot gb}, \lambda^{sv,rl,sv \cdot rl}, \beta^{sv,rl,sv \cdot rl}$ | 15.0 | 64966.14 (207.99) |
| Model 4 | $\alpha^{sv,rl,sv \cdot rl}, \lambda^{sv,rl,gb,sv \cdot gb,sv \cdot rl,rl \cdot gb}, \beta^{sv,rl,gb,sv \cdot gb,sv \cdot rl,rl \cdot gb}$ | 18.0 | 65388.15 (205.33) |
| Model 5 | $\alpha^{sv,rl,gb,sv \cdot gb,sv \cdot rl,rl \cdot gb}, \lambda^{sv,rl,gb,sv \cdot gb,sv \cdot rl,rl \cdot gb}, \beta^{sv,rl,gb,sv \cdot gb,sv \cdot rl,rl \cdot gb}$ | 21.0 | 64786.26 (208.39) |
| Model 6 | $\alpha^{sv,rl,gb,sv \cdot gb,sv \cdot rl,rl \cdot gb,sv \cdot gb \cdot rl}, \lambda^{sv,rl,gb,sv \cdot gb,sv \cdot rl,rl \cdot gb,sv \cdot gb \cdot rl}, \beta^{sv,rl,gb,sv \cdot gb,sv \cdot rl,rl \cdot gb,sv \cdot gb \cdot rl}$ | 24.0 | 64803.69 (208.43) |
| Model 7 | $\alpha^{sv,rl,gb,sv \cdot gb,sv \cdot rl,rl \cdot gb}, \lambda^{sv,rl,gb,sv \cdot gb,sv \cdot rl,rl \cdot gb}, \beta^{sv,rl,gb,sv \cdot gb,sv \cdot rl,rl \cdot gb}, r^{rl}$ | 23.0 | 64171.58 (209.9) |
| Model 8 | $\alpha^{sv,rl,gb,sv \cdot gb,sv \cdot rl,rl \cdot gb}, \lambda^{sv,rl,gb,sv \cdot gb,sv \cdot rl,rl \cdot gb}, \beta^{sv,rl,gb,sv \cdot gb,sv \cdot rl,rl \cdot gb}, r^{rl}, \epsilon^{rl}$ | 25.0 | 64140.71 (209.82) |
| Model 9 | $\alpha^{sv,rl,gb,sv \cdot gb,sv \cdot rl,rl \cdot gb}, \lambda^{sv,rl,gb,sv \cdot gb,sv \cdot rl,rl \cdot gb}, \beta^{sv,rl,gb,sv \cdot gb,sv \cdot rl,rl \cdot gb}, r^{rl}, \delta^{sv,rl,sv \cdot rl}$ | 27.0 | 64132.63 (208.9) |
| Model 10 | $\alpha^{sv,rl,gb,sv \cdot gb,sv \cdot rl,rl \cdot gb}, \lambda^{sv,rl,gb,sv \cdot gb,sv \cdot rl,rl \cdot gb}, \beta^{sv,rl,gb,sv \cdot gb,sv \cdot rl,rl \cdot gb}, r^{rl}, \delta^{sv,rl,sv \cdot rl}$ | 27.0 | 64044.19 (210.17) |
| Model 11 | $\alpha^{sv,rl,gb,sv \cdot gb,sv \cdot rl,rl \cdot gb}, \alpha_{ck}, \lambda^{sv,rl,gb,sv \cdot gb,sv \cdot rl,rl \cdot gb}, \beta^{sv,rl,gb,sv \cdot gb,sv \cdot rl,rl \cdot gb}, \beta_{ck}, r^{rl}$ | 26.0 | 64062.8 (210.43) |
| Model 12 | $\alpha^{sv,rl,gb,sv \cdot gb,sv \cdot rl,rl \cdot gb}, \alpha_{ck}, \lambda^{sv,rl,gb,sv \cdot gb,sv \cdot rl,rl \cdot gb}, \beta^{sv,rl,gb,sv \cdot gb,sv \cdot rl,rl \cdot gb}, \beta_{ck}, r^{rl}, \delta^{sv,rl,sv \cdot rl}$ | 30.0 | **63977.5 (210.81)** |
| Model 13 | $\alpha^{sv,rl,gb,sv \cdot gb,sv \cdot rl,rl \cdot gb}, \alpha_{ck}, \lambda^{sv,rl,gb,sv \cdot gb,sv \cdot rl,rl \cdot gb}, \beta^{sv,rl,gb,sv \cdot gb,sv \cdot rl,rl \cdot gb}, \beta_{ck}, r^{rl}, \delta^{sv,rl,sv \cdot rl}$ | 30.0 | 63882.5 (210.52) |

#### Model Comparison

We fitted all hierarchical models from Gagne et al. (2020) to behavioral data from the reversal learning task with reward and loss domains. Model 13 achieved the best fit, with a PSIS-LOO value of 63882.5 (standard error = 210.52; Table S8). However, despite all parameters having a 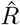 value below 1.1, the population-level parameters for this model exhibited high standard deviations. Given this, we selected

Model 12—the second-best fitting model—as our final model for analysis.

**Figure S31.**
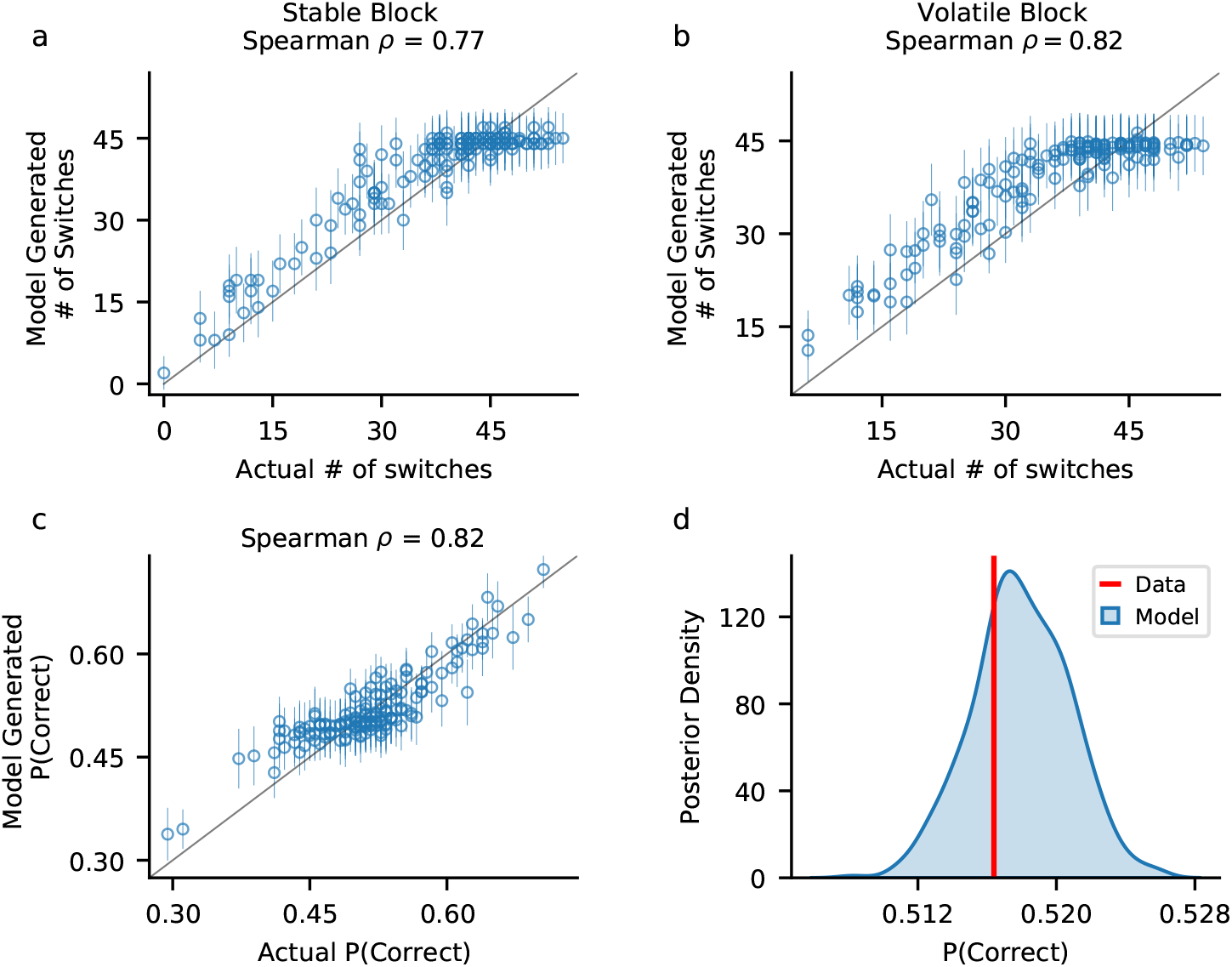
Posterior predictive checks for model 12. To assess how well the model 12 captured key qualitative features of choice behavior, we generated 500 simulated choice datasets per participant by drawing random samples from each participant’s joint posterior distribution (180 trials each for the reward and loss domain). **a-b**| Spearman correlations between the average number of switches across the task domains in simulations and actual participant switches in the stable (**a**) and volatile (**b**) task phases were strong, indicating that the model successfully reproduces switching behavior. Circles represent the average number of switches, and error bars denote the standard deviation across simulations. **c**| We found a high correlation between the proportion of trials in which the high-reward fractal was chosen (P(Correct)) in the simulated data and actual participant P(Correct). This suggests that the model captured choice accuracy accurately. **d**| Distribution of the model’s predicted P(Correct) across trials and participants. The red line represents the actual average P(Correct) across participants. The close alignment between the predicted distribution and the actual value further supports the model’s ability to replicate key behavioral patterns.

#### Model Reproduction of Basic Features

We used the procedure described in SM Model Reproduction of Basic Features to assess how well the posterior predictions from model 12 captured key qualitative aspects of participant behavior, focusing on switching patterns and choice accuracy combined across the reward and loss domains in model-generated data (Fig. S31). The number of switches in the simulated datasets closely matched the actual participant data, showing high Spearman correlations in both the stable (*ρ* = 0.78, Fig. S31a) and volatile (*ρ* = 0.82, Fig. S31b) task phases. Similarly, the model successfully reproduced choice accuracy, with a high correlation between simulated and actual P(Correct) (Spearman *ρ* = 0.82, Fig. S31c). Furthermore, the distribution of mean P(Correct) in the simulated data closely aligned with the actual average P(Correct) across participants (Fig. S31d). These results indicate that model 12 was able to effectively capture key behavioral patterns observed in participant choices.

**Figure S32.**
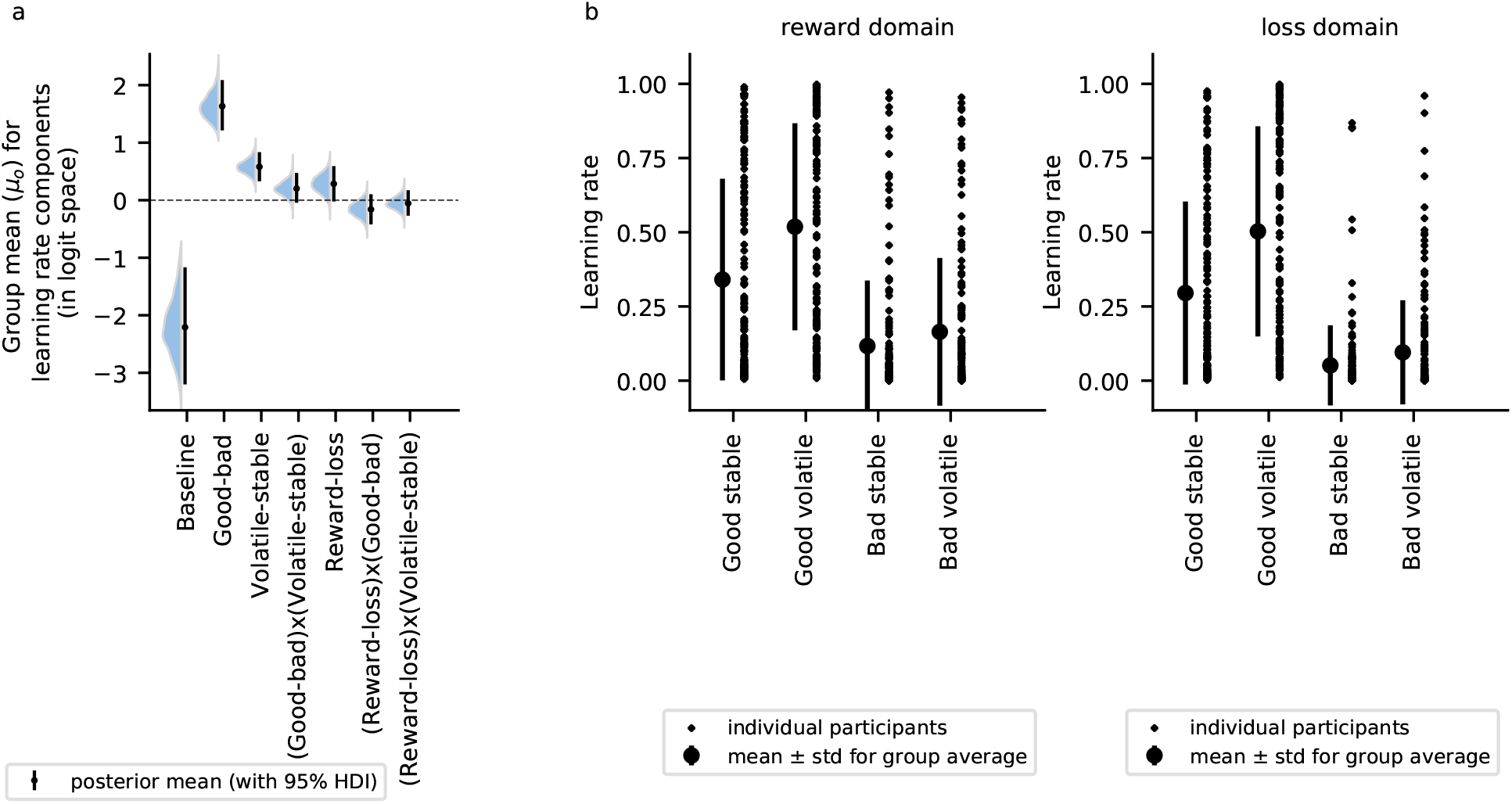
Mean learning-rate components and overall learning rates in the reversal learning task across reward and loss domains. **a**| Mean learning-rate components across all participants. Learning rates were significantly higher after positive outcomes compared to negative outcomes and during the volatile phase compared to the stable phase. **b**| Overall learning rates in the reward domain, categorized by different outcome valence and task phase conditions. The error bars represent the mean learning rate and standard error of the mean for each condition. **c**| Overall learning rates in the loss domain, categorized by different outcome valence and task phase conditions.

#### Model Results

Across participants, learning rates were significantly influenced by outcome valence (good vs. bad) and task phase (volatile vs. stable), but were not significantly affected by task domain (reward vs. loss) (Fig. S32a). Specifically, participants exhibited a higher learning rate following good outcomes compared to bad outcomes (*α*_*Good*− *bad*_ *µ*_0_ = 1.63, 95% HDI = [1.21, 2.09]). Learning rates were also higher in the volatile phase than in the stable phase (*α*_*V olatile* −*stable*_ *µ*_0_ = 0.58, 95% HDI = [0.33, 0.83]). However, we did not find systematic evidence of an interaction of outcome valence and task phase to influence the learning rate (*α*_(*Good* −*bad*)*x*(*V olatile*− *stable*)_ *µ*_0_ = 0.2, 95% HDI = [-0.04, 0.47]). Similarly, we did not find the learning rate to be significantly different between the reward and loss domains (*α*_*Reward* −*loss*_ *µ*_0_ = 0.28, 95% HDI = [-0.02, 0.59]). Additionally, we did not find significant interaction effects of outcome valence and task domain (*α*_(*Reward* −*loss*)*x*(*V olatile*− *stable*)_ *µ*_0_ = -0.16, 95% HDI = [-0.42, 0.1]) or of task phase and task domain on the learning rates (*α*_(*Reward*− *loss*)*x*(*Good*− *bad*)_ *µ*_0_ = -0.05, 95% HDI = [-0.27, 0.17]).

We did not find a significant effect of internalizing on any of the learning-rate components (Fig. S33a), with all the 95% HDI’s containing zero. Specifically, we did not find internalizing to have a significant influence on learning from outcome valence, task phase, or their interaction (*α*_*Baseline*_ *β*_*g*_ = 0.36, 95% HDI = [-0.22, 0.97], *α*_*Good*−*bad*_ *β*_*g*_ = -0.27, 95% HDI = [-0.6, 0.01], *α*_*V olatile*−*stable*_ *β*_*g*_ = -0.14, 95% HDI = [-0.36, 0.07], *α*_(*Good* −*bad*)*x*(*V olatile* −*stable*)_ *β*_*g*_ = -0.11, 95% HDI = [-0.33, 0.1]). Similarly, internalizing was not found to significantly affect learning across task domains (reward vs. loss) or their interactions with outcome valence or task phase (*α*_*Reward*− *loss*_ *β*_*g*_ = -0.06, 95% HDI = [-0.38, 0.2], *α*_(*Reward* −*loss*)*x*(*Good* −*bad*)_ *β*_*g*_ = -0.04, 95% HDI = [-0.28, 0.19], *α*_(*Reward* −*loss*)*x*(*V olatile*− *stable*)_ *β*_*g*_ = -0.2, 95% HDI = [-0.4, 0.01]). Further, dividing participants into low- and high-internalizing groups based on their general-factor scores and visualizing their learning rates across valence and task-phase conditions for each domain revealed no qualitative differences. Learning rates for good vs. bad outcomes in either the stable or volatile phase appeared comparable between groups across both task domains (Fig. S33b andS33c). These results align with findings from previous task versions, reinforcing the conclusion that internalizing does not impair learning under volatility in either the reward or loss domain.

**Figure S33.**
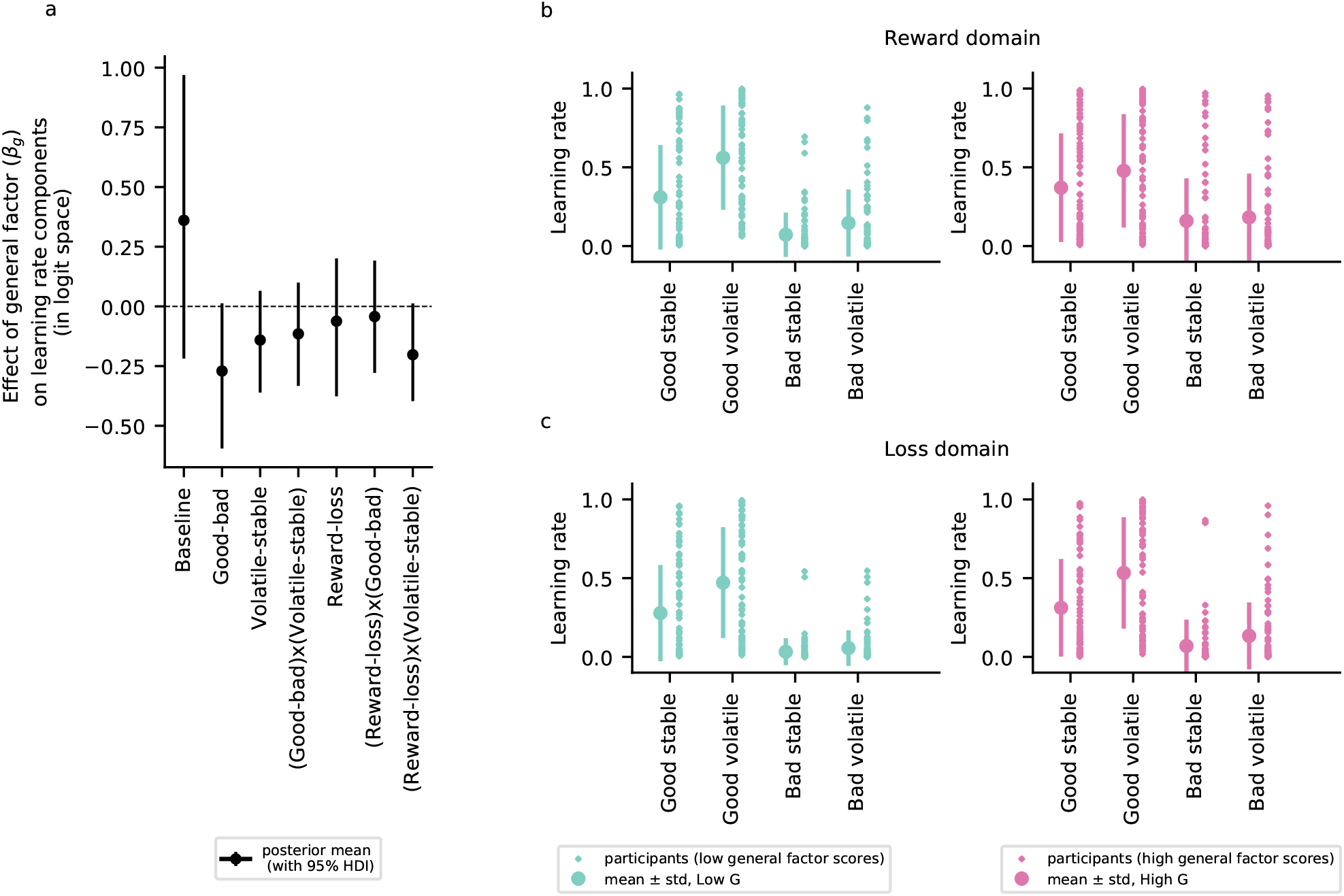
No significant effect of internalizing on the learning-rate components. **a**| We did not find a significant effect of internalizing on any of the learning-rate components, with all 95% HDIs containing zero. **b**,**c**| Participants were categorized into low- and high-internalizing groups, with their learning rates for different outcome valence and task phase conditions shown for the reward domain (**b**) and the loss domain (**c**).

**Figure S34.**
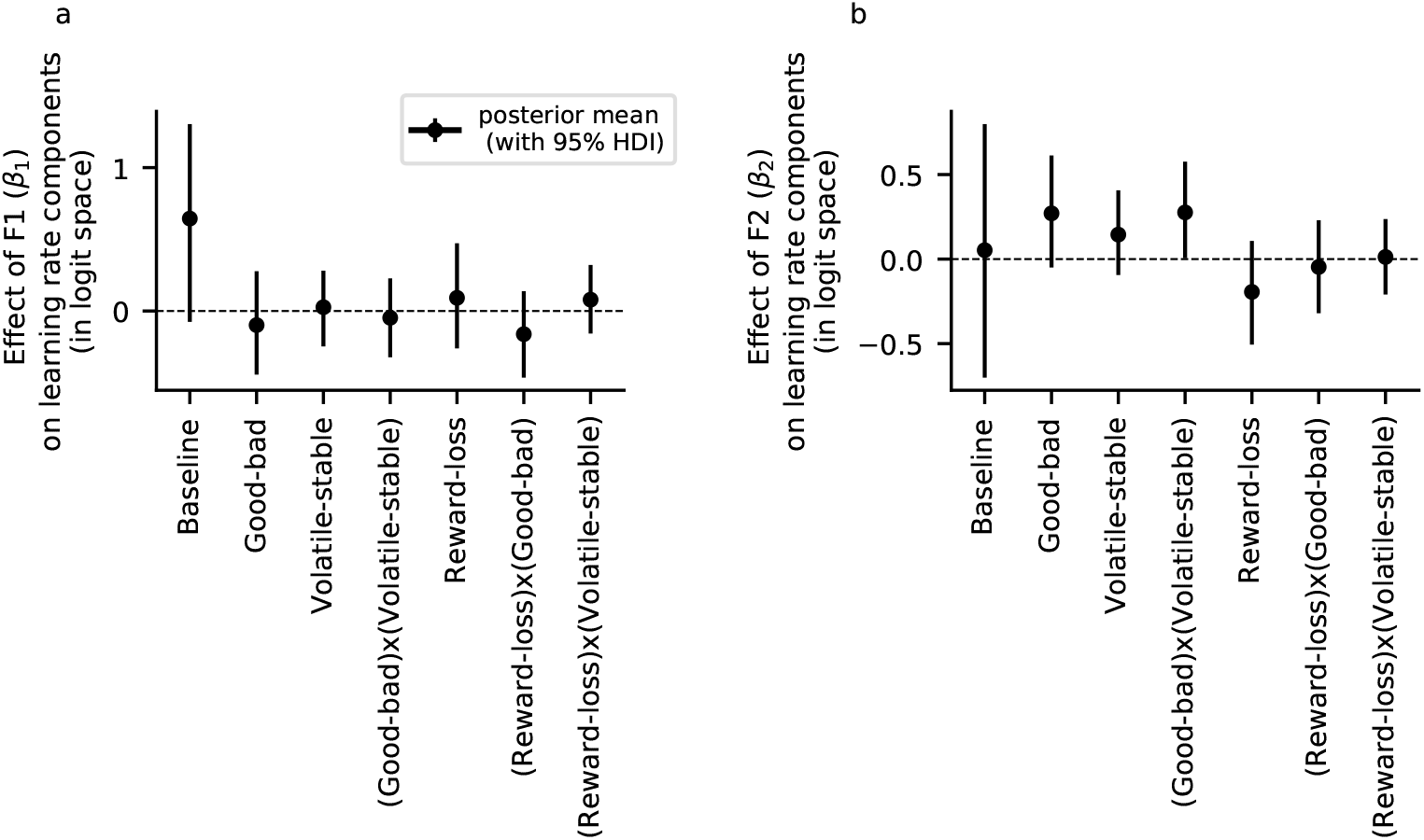
Effect of the anxiety-related factor F1 and the depression-related factor F2 on learning-rate components. **a**| We did not find a significant effect of the anxiety-related factor on any learning-rate component, with all 95% highest-density intervals (HDIs) containing zero. **b**| The depression-related factor showed a marginally significant effect on the interaction between outcome valence and task phase, suggesting higher learning from good compared to bad outcomes in the volatile phase relative to the stable phase.

**Figure S35.**
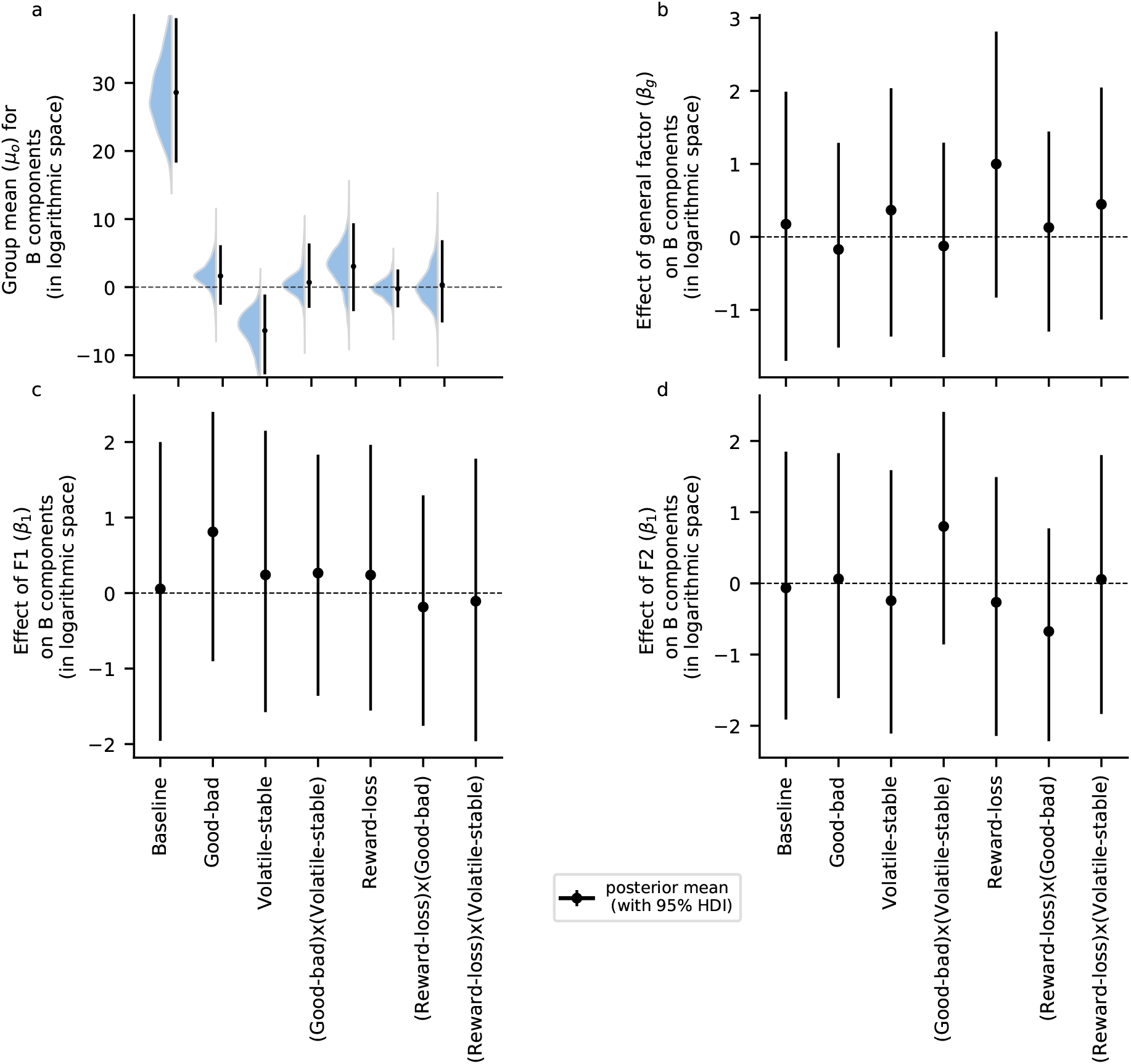
Mean inverse temperature and effects of internalizing, the anxiety-related factor F1, and the depression-related factor F2 on the inverse temperature parameter in the reversal learning task with reward and loss domains. **a**| Group mean (*µ*_0_, population-level parameter) of inverse temperature across participants revealed a significant effect of task phase, with inverse temperature being lower in the volatile phase compared to the stable phase. This suggests an increased choice stochasticity in the volatile phase compared to the stable phase. **b**| We did not find a significant effect of internalizing (*β*_*g*_, population-level parameter) on any inverse temperature components, with all corresponding 95% HDIs containing zero. **c**| We also did not find a significant effect of the anxiety-related factor (*β*_1_, population-level parameter) on any inverse temperature components, with all corresponding 95% highest-density intervals (HDIs) containing zero. **c**| Similarly, we did not find a significant effect of the depression-related factor (*β*_2_, population-level parameter) on any inverse temperature components, with all corresponding 95% HDIs containing zero.

Additionally, the analysis did not reveal a significant effect of the anxiety-related factor F1 on any learning-rate component (Fig. S34a). However, we found a marginal effect of the depression-related factor F2 on the interaction between outcome valence and task phase, with higher learning from good compared to bad outcomes in the volatile phase relative to the stable phase (*α*_(*Good* −*bad*)*x*(*V olatile*− *stable*)_ *β*_2_ = 0.28, 95% HDI = [0.01, 0.58]; Fig. S34b). While this finding suggests a tendency toward greater learning from positive outcomes in individuals with higher depression-related symptoms, it should be interpreted with caution, given the marginal nature of the effect and the lack of replication across other tasks.

Next, we examined the inverse temperature parameter and its relationship with internalizing, anxiety, and depression (Fig. S35). Across participants, we found a significant decrease in inverse temperature in the volatile phase compared to the stable phase, indicating that participants made more stochastic choices in the volatile phase (*B*_*V olatile* −*stable*_ *µ*_0_ = -6.39, 95% HDI = [-12.81, -1.09]). However, we did not observe a significant effect of outcome valence (good vs. bad), task domain (reward vs. loss), or any of the interactions on overall inverse temperature (*B*_*Good*−*bad*_ *µ*_0_ = 1.63, 95% HDI = [-2.59, 6.16], *B*_(*Good*− *bad*)*x*(*V olatile* −*stable*)_ *µ*_0_ = 0.69, 95% HDI = [-3.04, 6.42], *B*_*Reward* −*loss*_: *µ*_0_ = 3.05, 95% HDI = [- 3.53, 9.38], *B*_(*Reward*− *loss*)*x*(*Good* −*bad*)_ *µ*_0_ = -0.23, 95% HDI = [-2.97, 2.59], *B*_(*Reward* −*loss*)*x*(*V olatile*− *stable*)_ *µ*_0_ = 0.32, 95% HDI = [-5.19, 6.89]; Fig. S35a). We did not find a significant effect of internalizing (general factor G, Fig. S35b), the anxiety-related factor F1 (Fig. S35c), or the depression-related factor F2 (Fig. S35d) on any inverse temperature component. These results broadly align with findings from the previous task version with reward magnitudes, reinforcing their robustness. Furthermore, the results remained consistent when analyzing parameters from Model 13.

